# Context-dependent regulatory variants in Alzheimer’s disease

**DOI:** 10.1101/2025.07.11.659973

**Authors:** Ziheng Chen, Yaxuan Liu, Ashley R. Brown, Heather H. Sestili, Easwaran Ramamurthy, Xushen Xiong, Dmitry Prokopenko, BaDoi N. Phan, Lahari Gadey, Robert Van De Weerd, Karsen E. Shoger, Peinan Hu, Shiyu Wang, Irene Kaplow, Rachel A. Gottschalk, Li-Huei Tsai, Lars Bertram, Winston Hide, Rudolph E. Tanzi, Manolis Kellis, Andreas R. Pfenning

## Abstract

Noncoding genetic variants underlie many complex diseases, yet identifying and interpreting their functional impacts remains challenging. Late-onset Alzheimer’s disease (LOAD), a polygenic neurodegenerative disorder, exemplifies this challenge. The disease is strongly associated with noncoding variation, including common variants enriched in microglial enhancers and rare variants nominated at loci implicated in neurodevelopment and synaptic function. These variants may perturb regulatory sequences by disrupting transcription factor (TF) motifs or altering local regulatory sequence context, with potential consequences for gene expression and chromatin accessibility. However, assessing their impact is complicated by the context-dependent functions of regulatory sequences, underscoring the need to systematically examine variant effects across diverse tissues, cell types, and cellular states.

Here, we combined *in vitro* and *in vivo* massively parallel reporter assays (MPRAs) with interpretable machine-learning models to systematically characterize common and rare variants across myeloid and neuron-enriched neural contexts. Parallel profiling of variants in four immune states *in vitro* and three mouse brain regions *in vivo* revealed that individual variants can differentially and even oppositely modulate reporter activity across cellular contexts, while a subset showed immune-state-dependent effects. Within the assayed candidate set, the relative effect sizes of common and rare variants reversed between contexts, with common variants showing larger effects in THP-1 macrophages and rare variants in brain tissue. Interpretable sequence-to-function models prioritized context-biased variants and generated motif-level hypotheses, including predicted transcription-factor motif disruption and subtler changes in local motif context. To probe endogenous consequences at a prioritized locus, we used CRISPR interference to perturb a rare-variant-containing enhancer at the *SEC63–OSTM1* locus, revealing metabolic and biosynthetic transcriptional programs resembling those induced by *SEC63* promoter perturbation, together with a stimulation-specific interferon response. These findings show that LOAD-associated noncoding variants exhibit context-dependent effects on reporter activity and predicted chromatin accessibility, and provide a framework for linking genetic association to regulatory activity, sequence grammar and endogenous transcriptional state.

## Main

Late-onset Alzheimer’s disease (LOAD), the most prevalent neurodegenerative disorder worldwide, exhibits strong heritability^1^. Its genetic architecture comprises multiple components, including the strong influence of the *APOE* ε4 allele^2^, common variants with weaker effects, and rare variants. Genome-wide association studies (GWAS)^3–6^ have identified numerous common variants (minor allele frequency (MAF) ≥ 0.01) that are enriched for functions in immune cells, particularly microglia and macrophages. This genetic landscape implicates neuroinflammation and mirrors the broad observed failure of microglial functions in LOAD brains^7,8^: impaired surveillance^9,10^, reduced amyloid-β^11,12^ and tau clearance^13,14^, dysregulated injury response^15,16^, and altered inflammatory regulation^17,18^. Notably, top risk genes such as *BIN1* and *TREM2* encode proteins crucial for maintaining myeloid homeostasis; dysfunction in these genes perturbs processes like endocytosis^19,20^, lipid metabolism^19,21^, and cytokine signaling^19,22^, thereby accelerating neurodegeneration^23,24^. Human single-nucleus transcriptomic studies further delineate distinct microglial activation states, and several of these states display more prominent LOAD-associated gene expression signatures, suggesting state-specific genetic effects^25,26^. Although fine-mapping efforts are increasingly narrowing down GWAS signals^27–31^ in the immune context, pinpointing the exact causal variants and their functional mechanisms remains a major challenge.

In contrast to GWAS, whole-genome sequencing (WGS) of 2,247 individuals from 605 multiplex LOAD families uncovered 13 loci with consistent association of rare variant (MAF < 0.01) clusters, such as *CTNNA2, KIF2A, SYTL3, CLSTN2, NALCN,* and *FNBP1L*, whose products are pivotal for synaptic function and neurodevelopment^32^. These loci suggest that neural and synaptic regulatory mechanisms may represent an additional component of LOAD genetic risk. Nevertheless, limited statistical power for individual rare variants within these clusters obscures the identity of the functional candidate alleles, and their precise functional consequences therefore remain unknown. Thus, direct functional testing can prioritize individual rare alleles and compare their regulatory potential with common GWAS-derived candidates. Furthermore, immune-versus-neural cell-type divergence underscores the importance of analyzing genetic variants in disease-relevant cell types and states. Consequently, there is a critical need for systematic, large-scale functional measurements that directly compare common and rare variants, coupled with detailed mechanistic interpretations in disease-relevant cellular contexts.

Both the GWAS and WGS studies suggest that LOAD is predominantly driven by noncoding variants^3–6,32^. Noncoding genetic variants contribute to the risk of many complex diseases by disrupting gene regulation, often through interference with transcription factor (TF) binding at *cis*-regulatory elements (CREs) such as enhancers and promoters^33–35^. The activity of these CREs can be highly cell-type-specific and even state-specific, shaped by the availability and dynamics of distinct TFs^36–39^. This specificity implies that the effects of genetic variants on CRE function, and consequently on disease risk, are likewise context-dependent^40,41^. LOAD GWAS variants show significant enrichment in microglia and macrophage enhancers, potentially disrupting immune enhancer activities^42,43^. A well-characterized CRE is the microglia-specific enhancer ∼25 kb upstream of *BIN1* that harbors the lead LOAD risk variant rs6733839. CRISPR-mediated deletion of this enhancer in human iPSC-derived microglia selectively lowers *BIN1* mRNA and protein levels, suggesting rs6733839 may modulate LOAD risk by altering microglial *BIN1* expression^43^. Although epigenomic modeling that integrates open chromatin, histone marks, and three-dimensional contacts now permits prediction of variant impacts on these CREs, current models still struggle to resolve the resulting transcriptional and cellular consequences because of the versatile, combinatorial nature of TF function.

Quantifying the functional consequences of noncoding genetic variation is intrinsically difficult: regulatory elements act in a highly context-specific manner and typically elicit subtler phenotypic changes than coding mutations^44^. Massively parallel reporter assays (MPRAs) overcome many technical barriers by coupling thousands of synthesized CREs to barcoded reporter constructs, positioned either upstream or downstream of a minimal promoter, to measure sequence-dependent enhancer activity ^45^. Allelic MPRAs directly contrast CREs that differ by a single nucleotide and provide a robust, high-throughput readout of allele-specific regulatory effects^30,31,46–48^. Traditional *in vitro* MPRAs can achieve high delivery efficiency and statistical power, yet they only approximate the *in vivo* cellular identity of their target cell type. Recent *in vivo* tissue MPRAs, including our AAV-based systemic platform (sysMPRA)^49^, extended this paradigm to native neural contexts in adult brain tissue using neuron-biased AAV vectors, enabling neuron-enriched measurement of regulatory activity within intact multicellular environments despite limited cell-type resolution^49,50^. Single-cell^51,52^ and spatial^53,54^ reporter assays offer a route to assigning regulatory activity to specific cell types and states within heterogeneous tissues, although current implementations remain difficult to scale to large CRE libraries with the barcode coverage and statistical power achieved by conventional bulk MPRA. Thoughtful selection of the experimental system therefore remains critical for accurately inferring variant function. Additionally, while MPRA effectively quantifies variant impacts on transcription, its ability to resolve downstream effects on endogenous gene expression remains limited. To bridge this gap, methods such as CRISPR interference (CRISPRi)^55^ and CRISPR-based excision^43^ have been employed to perturb enhancers, enabling measurement of their effects on endogenous gene expression and LOAD-relevant transcriptional programs, though still at a relatively limited scale^28,30,43^. MPRA screens measure the sequence-intrinsic enhancer activity of CREs outside their native loci, whereas CRISPRi tests whether prioritized elements alter endogenous transcriptional states in native chromatin, thereby providing complementary rather than interchangeable evidence for regulatory variant function. Collectively, these experimental advances establish a versatile toolkit for disentangling the consequences of regulatory variants across diverse cellular contexts.

Complementing these experimental approaches, computational methodologies continue to evolve to interpret how genetic variants influence cell-type- and cell-state-dependent gene expression. Early computational frameworks relied on position weight matrices representing individual TF motifs^56^. Over the past decade, sequence-to-function models such as DeepSEA^57^, Enformer^58^, ChromBPNet^59^, DeepAllele^60^ and GET^61^ have advanced the field by using deep neural networks to learn combinatorial regulatory codes directly from DNA. Although predicting gene expression from sequence alone remains difficult, these architectures now deliver state-of-the-art accuracy on chromatin-feature prediction. Within this deep-learning landscape, convolutional neural networks (CNNs)—neural networks that apply learnable filters to scan sequence features such as TF motifs—continue to excel at regulatory genomics tasks, including cell-type-specific enhancer classification^62^ and genetic variant effect inference^27^. Although transformer-based, self-supervised genomic large language models aspire to achieve zero-shot prediction by learning the modeling genome-wide sequence context and long-range dependencies, supervised CNNs trained *de novo* and their derivatives^58^ currently deliver superior predictive performance, greater computational efficiency, and enhanced interpretability for uncovering underlying biology^63,64^. Recently, multimodal sequence-to-function models such as AlphaGenome^65^ have emerged, unifying diverse functional genomic outputs for variant-effect prediction across tissues and cell types. Although these models can capture cell-type- and tissue-specific regulatory signals, accurately predicting regulatory effects across unseen cellular states and experimental conditions remains challenging. Task-specific CNNs may therefore remain valuable when models can be trained directly on matched cellular or experimental contexts^54,66^. Together, these advances in deep learning provide a powerful, yet still evolving, framework for mechanistically informed prediction of variant effects on gene regulation, particularly when integrated with experimental assays that directly measure regulatory function.

To investigate how noncoding LOAD genetic variants affect gene regulatory activities across immune and neural cellular contexts, we assayed 599 candidate sites (588 LOAD-associated variants from GWAS^4,6^ and WGS^32^, and 11 AD-associated CpG sites from epigenome-wide association study^67^, Methods), including 175 rare variants (MAF < 0.01), 413 common variants (MAF ≥ 0.01), prioritizing sites overlapping accessible chromatin in microglia, macrophages, and neurons (Fig. 1a). We integrated chromatin-accessibility profiles across a broad panel of immune and brain-related cell types and classified sites based on the accessibility pattern of their surrounding candidate regulatory elements (Fig. 1b; see Methods). This analysis resolved five categories: Broad (n = 199), Immune (n = 246), Neuronal (n = 38), Glia (n = 22), and Low ATAC (n = 94). Across categories, sites were frequently found in intronic, intergenic, and promoter-proximal regions, consistent with localization in noncoding regulatory DNA.

**Fig. 1.**
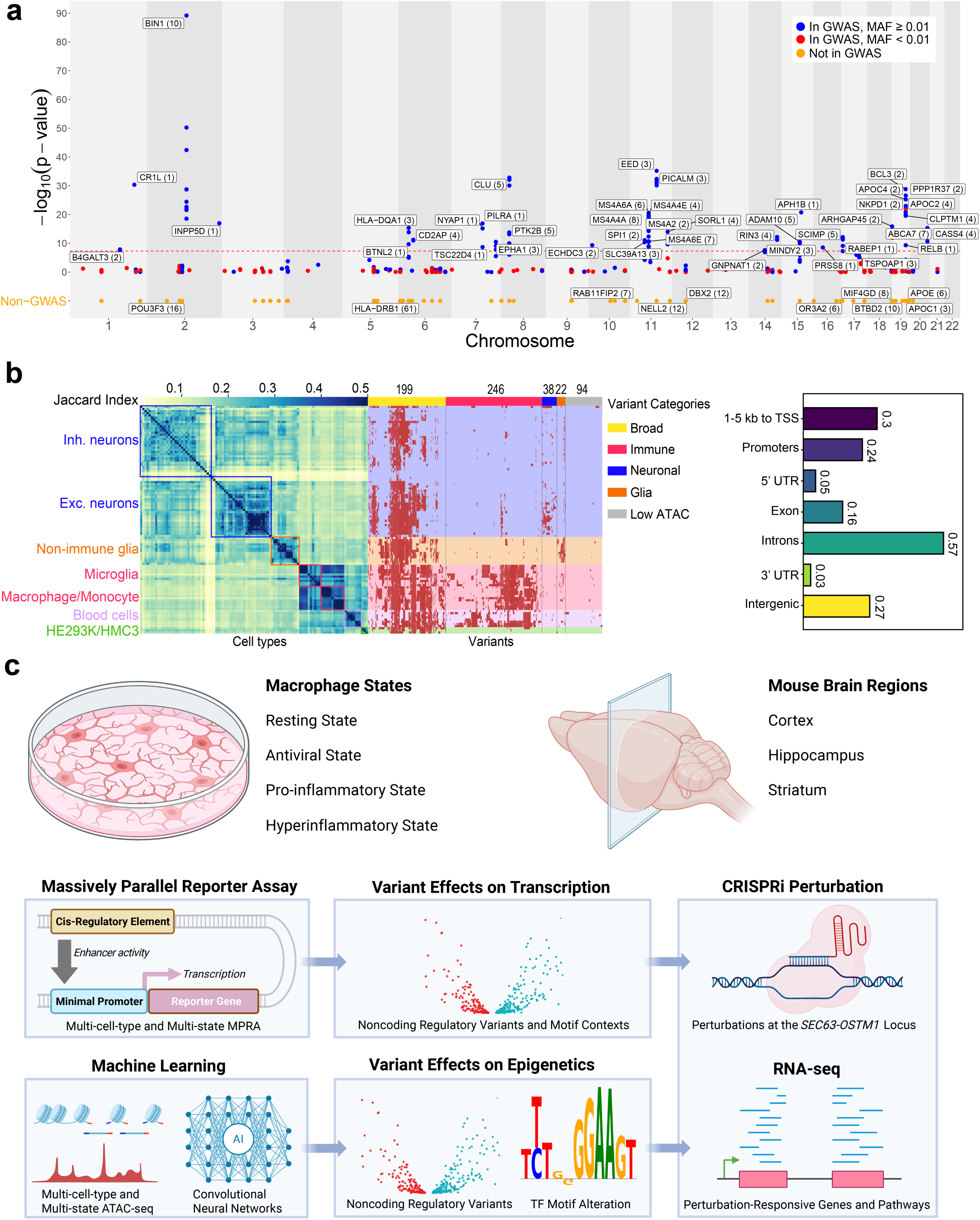
Study design and integrative framework for identifying context-dependent regulatory effects of LOAD-associated variants. **(a)** Chromosomal distribution of the candidate late-onset Alzheimer’s disease (LOAD)-associated variants assayed in this study. For variants present in the Bellenguez et al. AD GWAS^5^, the y-axis shows −log₁₀(*P*) as reported in that study; variants absent from the GWAS are plotted on a separate baseline. Blue points, variants present in the GWAS with minor allele frequency (MAF) ≥ 0.01 (n = 366); red points, variants present in the GWAS with MAF < 0.01 (n = 97); orange points, variants not present in the GWAS (n = 136). Labeled points denote selected loci or genes, with the number in parentheses indicates the number of variants in the corresponding plotted GWAS/MAF category at that locus. The dashed red line indicates the genome-wide significance threshold (*P* = 5 × 10⁻^8^). **(b)** Chromatin-accessibility-based classification and genomic annotation of candidate variants. Left, heatmap of pairwise Jaccard indices between cell-type-specific chromatin-accessibility profiles. Middle, heatmap showing the overlap of candidate variants with accessible chromatin across cell types, together with their assignment to the Broad, Immune, Neuronal, Glia, and Low ATAC categories. Numbers above the heatmap indicate the number of variants assigned to each category. Right, horizontal bar plot showing the fraction of candidate variants overlapping each genomic annotation, including regions 1–5 kb from a transcription start site (TSS), promoters, 5′ UTRs, exons, introns, 3′ UTRs, and intergenic regions. Numbers next to the bars indicate the fraction of candidate variants overlapping each annotation. Genomic annotation categories are not necessarily mutually exclusive. **(c)** Schematic overview of the experimental strategy integrating immune and neural model systems, massively parallel reporter assays (MPRAs), machine-learning predictions, and endogenous perturbation. THP-1-derived macrophages were assayed in resting, antiviral, pro-inflammatory, and hyperinflammatory states. Mouse brain MPRA measurements were obtained from the cortex, hippocampus, and striatum. MPRA was used to quantify candidate *cis*-regulatory element activity across cell types and states through reporter transcription driven by a minimal promoter. Allele-specific MPRA measurements were used to assess the effects of noncoding regulatory variants and their local sequence and transcription factor motif contexts on transcription. In parallel, convolutional neural networks trained on multi-cell-type and multi-state ATAC-seq profiles were used to predict variant effects on chromatin accessibility and identify altered transcription factor motifs. The prioritized regulatory region at the *SEC63–OSTM1* locus was further examined by CRISPR interference (CRISPRi), followed by RNA-seq analysis of perturbation-responsive genes and pathways.

In this study, we established an integrated experimental and computational framework that combines context-dependent *in vitro* and *in vivo* MPRA, interpretable CNN models, and CRISPRi perturbation (Fig. 1c). Using characterized THP-1 macrophages as a scalable and perturbable myeloid regulatory system, mouse brain tissue as a neuron-enriched neural context, HMC3 cells as an immortalized CNS-derived cell line historically used as a microglial model but exhibiting a noncanonical, largely non-myeloid phenotype, and HEK293T cells as a permissive non-immune comparison context, this approach (i) quantifies the regulatory impact of hundreds of LOAD-associated variants, (ii) prioritizes functionally impactful alleles, (iii) dissects their cell-type- and immune-state-specific mechanisms, and (iv) enables in-depth functional analysis of a disease-risk–associated enhancer harboring functional rare variants.

Together, this framework links LOAD-associated noncoding variation to context-dependent enhancer activity and candidate sequence-level regulatory mechanisms, while enabling endogenous transcriptional consequences to be tested at prioritized loci.

## Results

### Characterization of THP-1 macrophages as a LOAD-relevant immune-responsive myeloid model

Mononuclear phagocytes, including microglia, macrophages, and monocytes, are implicated in the genetic predisposition to LOAD^5,6,27,42,68^. Microglia, the resident macrophages of the central nervous system, continuously survey the brain parenchyma^9,10^, clear amyloid-β (Aβ)^11,12^ and tau aggregates^13,14^, respond to neural injury^15,16^, and modulate neuroinflammation^17,18^, all of which are processes dysregulated in LOAD^7,8^. Although microglia are the primary brain-resident immune cells, perivascular macrophages and recruited monocyte-derived macrophages also participate in plaque clearance and secrete proinflammatory cytokines^69–72^. In microglia-depleted mouse models of LOAD, monocyte-derived macrophages repopulated the brain and functionally substituted for microglia, as evidenced by their association with amyloid plaques and expression of *Trem2*^73^. Moreover, peripheral inflammation could induce microglial immune memory through epigenetic reprogramming, modifying LOAD pathology^74^. Together with microglia, peripheral immune cells are also increasingly recognized as LOAD modulators^27,75^.

To model mononuclear phagocytes and immune responses relevant to LOAD-associated microglial states^25,26^, we adopted THP-1 cells as a scalable and perturbable innate-immune system, leveraging their transcriptomic similarity to primary microglia and macrophages^76^ and their ease of genetic and pharmacological manipulation. HMC3 cells, historically used as a human microglia-like model but recently reported to exhibit more astrocyte-like than microglial features^77,78^, were retained as a complementary immortalized CNS-derived cell-line context. Using chromatin accessibility (ATAC-seq) and transcriptome profiling (RNA-seq), we characterized THP-1 macrophages throughout differentiation and after stimulation with interferon-β (IFN-β), interferon-γ (IFN-γ), or lipopolysaccharide plus IFN-γ (LPS+IFN-γ), chosen to model three mechanistically distinct innate immune programs that capture selected antiviral and inflammatory features of immune states associated with neuroinflammation and Alzheimer’s disease: a type I interferon–responsive antiviral state^79,80^, a type II interferon–responsive pro-inflammatory state^81^, and an LPS+IFN-γ–driven hyperinflammatory-like state^82^, respectively (Supplementary Tables 1 and 2).

Upon differentiation from THP-1 monocytes, THP-1 macrophages showed increased immune-response programs and reduced cell-cycle activity, consistent with successful differentiation (Extended Data Fig. 1a). Consistent with rapid immune signaling, nuclear pSTAT1 and pSTAT3 showed early and condition-specific activation before the 4 h profiling time point, supporting the selection of this time point to capture early immune-response programs before the accumulation of later secondary cytokine-mediated responses (Extended Data Fig. 2b). At 4 h, promoter-level analysis showed that IFN-β and IFN-γ treatments predominantly increased chromatin accessibility, whereas LPS+IFN-γ produced broader bidirectional changes involving both accessibility gains and losses. Parallel RNA-seq analyses showed a similar directional pattern: IFN-β and IFN-γ predominantly induced gene expression, whereas LPS+IFN-γ elicited broader transcriptional changes involving both gene activation and repression (Extended Data Fig. 1b,c and 2a,c).

Promoters gaining accessibility after IFN-β treatment were associated with genes enriched for antiviral-defense and leukocyte-migration biological processes, whereas those gaining accessibility after IFN-γ treatment were associated with responses to molecules of bacterial origin and lipids, as well as protein transport and localization. Promoters gaining accessibility after LPS+IFN-γ treatment were associated with immune-regulatory processes, including defense responses, leukocyte activation, and cytokine production. RNA-seq gene set enrichment analysis showed that all three treatments activated overlapping interferon-associated programs, with LPS+IFN-γ eliciting the broadest inflammatory transcriptional response across the pathways examined (Extended Data Fig. 1d and 2d). Across the IFN-β, IFN-γ, and LPS+IFN-γ time courses, differential chromatin accessibility was already evident at 4 h and continued to evolve at later time points (Supplementary Fig. 1). Together, these chromatin-accessibility and transcriptional profiles supported the intended distinction among the three stimulation conditions.

**Fig. 2.**
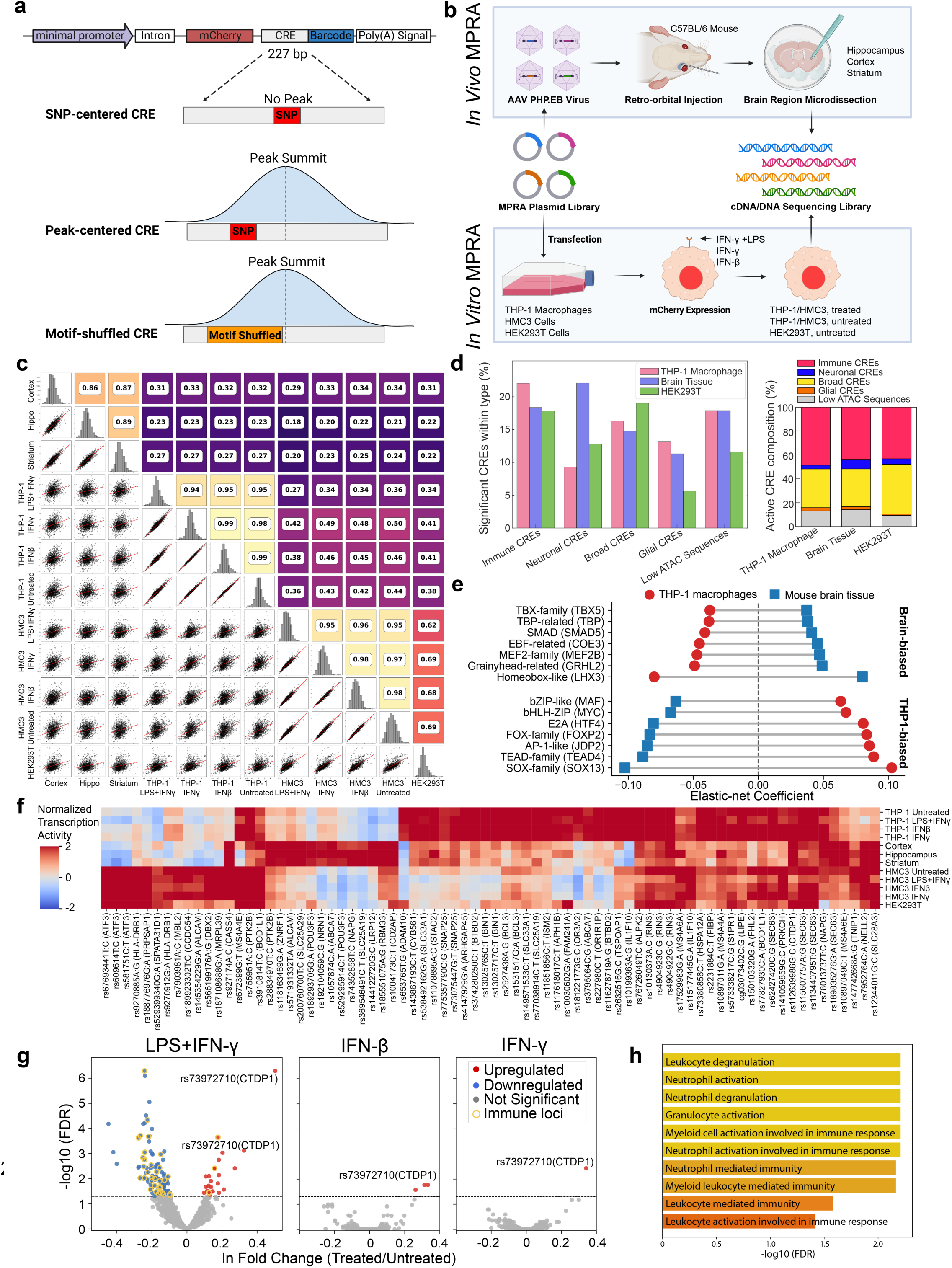
Context-dependent MPRA reveals regulatory activity of LOAD-associated candidate elements across immune-cell and brain-tissue contexts. **(a)** MPRA reporter construct and candidate *cis*-regulatory element (CRE) design strategy. A minimal promoter drove mCherry reporter expression, with synthesized 227-bp sequences representing candidate CREs and associated barcode sequences positioned downstream of the reporter. SNP-centered CREs positioned the candidate variant at the center of the assayed sequence. Peak-centered CREs were centered on the summit of the corresponding open-chromatin peak. Motif-shuffled CREs contained a shuffled 25-bp sequence surrounding selected variants. **(b)** Experimental workflows for *in vivo* and *in vitro* MPRA. For *in vivo* MPRA, the plasmid library was packaged into AAV.PHP.eB and delivered to C57BL/6 mice by retro-orbital injection. The cortex, hippocampus, and striatum were subsequently microdissected, and matched cDNA and DNA sequencing libraries were prepared. For *in vitro* MPRA, the plasmid library was transfected into THP-1 macrophages, HMC3 cells, and HEK293T cells. THP-1 macrophages and HMC3 cells were analyzed under untreated conditions and following IFN-β, IFN-γ, or LPS + IFN-γ treatment, whereas HEK293T cells were analyzed under untreated conditions. Reporter activity was quantified using matched cDNA and DNA barcode sequencing libraries. **(c)** Pairwise comparison of candidate CRE activity across mouse brain regions, THP-1 macrophage conditions, HMC3 conditions, and HEK293T cells. MPRA transcriptional activity was estimated from cDNA:DNA barcode ratios using one-sided MPRAnalyze analysis and transformed into median absolute deviation (MAD) scores. Cortex, hippocampus, and striatum indicate *in vivo* mouse brain MPRA samples. THP-1 Untreated, THP-1 IFNβ, THP-1 IFNγ, and THP-1 LPS+IFNγ indicate untreated, IFN-β-treated, IFN-γ-treated, and LPS+IFN-γ-treated THP-1 macrophages, respectively. HMC3 Untreated, HMC3 IFNβ, HMC3 IFNγ, and HMC3 LPS+IFNγ indicate the corresponding HMC3 conditions. HEK293T represents the non-immune control context. Diagonal panels show the distribution of MAD scores in each context. Lower-triangle panels show pairwise comparisons of MAD scores, with red lines indicating linear fits. Upper-triangle panels show Pearson correlation coefficients, with tile color indicating correlation strength. **(d)** Left, percentage of significant candidate CREs within each predefined CRE category in THP-1 macrophages, mouse brain tissue, and HEK293T cells. Right, category composition of active CREs in THP-1 macrophages, mouse brain regions, and HEK293T cells. CRE categories include Immune, Neuronal, Broad, Glia, and Low ATAC. **(e)** Selected transcription factor motif-family coefficients derived from elastic-net models of context-specific MPRA enhancer activity in THP-1 macrophages and mouse brain tissue. Red circles indicate coefficients for THP-1 macrophages, blue squares indicate coefficients for mouse brain tissue, and gray lines connect coefficients for the same motif group across the two contexts. Motif groups are ordered according to their brain-or THP-1-biased associations. **(f)** Heatmap of Z-normalized MPRA transcriptional activity for selected CREs across THP-1 macrophage conditions, mouse brain regions, HMC3 conditions, and HEK293T cells. Rows indicate assay contexts, and columns indicate CREs labeled by variant and nearby gene. Color indicates normalized transcriptional activity. **(g)** Volcano plots showing stimulation-responsive CRE activity in THP-1 macrophages following LPS + IFN-γ, IFN-β, or IFN-γ treatment relative to untreated cells. The x-axis shows the natural logarithm of the fold change in MPRA activity between treated and untreated cells, and the y-axis shows −log10(FDR). Red points indicate upregulated CREs, blue points indicate downregulated CREs, gray points indicate nonsignificant CREs, and yellow-outlined points indicate immune loci. The dashed horizontal line denotes FDR = 0.05. **(h)** Gene ontology enrichment analysis of genes nearest to CREs repressed following LPS + IFN-γ treatment in THP-1 macrophages (FDR < 0.05 and log fold change < 0). Bars show −log10(FDR) values for enriched immune-related biological processes.

Representative mononuclear phagocyte-associated genes, including *FCER1G*, *TREM2*, *CSF1R*, *AIF1*, and *TYROBP*, were detectably expressed across THP-1 monocyte and macrophage conditions, consistent with a mononuclear phagocyte-like cellular context (Extended Data Fig. 2e). Interferon-response and inflammatory marker genes also showed condition-consistent expression patterns across biological replicates, supporting the reproducibility of the stimulation responses (Supplementary Fig. 2).

Previous studies have shown that THP-1 macrophages exhibit transcriptomic similarity to primary and stem-cell-derived microglia^76^; consistent with this, their chromatin accessibility profiles clustered with or adjacent to those of iPSC- and hESC-derived microglia. In contrast, our HMC3 ATAC-seq profiles were separated from the myeloid and microglial samples and occupied a region of PCA space closer to astrocytes and several immortalized non-myeloid cell lines, including HEK293 and SK-N-SH cells (Extended Data Fig. 1e).

Human single-nucleus transcriptomic studies of post-mortem brain tissue and cortical biopsies from living individuals have defined approximately 12 distinct microglial states, several of which are altered in LOAD^25,26^. In the Sun et al. post-mortem brain study^25^, three inflammatory states, MG10, MG2, and MG8, were ordered by decreasing inflammatory strength, while AD genetic enrichment was strongest in MG8 and MG2, with additional enrichment in MG0 (Homeostatic), MG4 (Lipid Processing), and MG11 (Antiviral). We next asked whether THP-1 macrophages under different stimulation conditions exhibited transcriptional programs resembling the signatures of these reported human microglial states, focusing on inflammatory and antiviral axes most directly related to our stimulation design. Gene set enrichment analysis showed that LPS+IFN-γ produced the broadest and strongest enrichment of human microglial state signatures in THP-1 macrophages (Extended Data Fig. 2f). Notably, LPS+IFN-γ enriched signatures of inflammatory microglial states, including MG10, MG2, and MG8. Enrichment was strongest for MG10 and MG2, indicating that the LPS+IFN-γ response aligned most closely with strongly inflammatory microglial-state programs. LPS+IFN-γ also enriched ribosome-biogenesis MG3, stress-related MG6, and antiviral MG11 signatures, together with negative enrichment of the cycling MG12 state. By comparison, IFN-β and IFN-γ showed more restricted enrichment patterns. IFN-β prominently enriched antiviral MG11 and phagocytic MG5 signatures, together with inflammatory MG2, whereas IFN-γ most prominently enriched inflammatory MG2 and antiviral MG11. Inflammatory microglial states are closely associated with amyloid pathology in human AD brains^25,26^. Sun et al. showed that amyloid-β fibrils induce inflammatory responses in iPSC-derived microglia^25^. Consistent with this observation, the 4 h LPS+IFN-γ response in THP-1 macrophages most closely matched the late inflammatory response (24 h) induced by amyloid-β fibrils (Extended Data Fig. 2g), supporting the use of LPS+IFN-γ as an experimentally controlled, AD-relevant inflammatory context. Together, these findings indicate that stimulated THP-1 macrophages capture selected transcriptional features of microglial states identified in human brains, with LPS+IFN-γ producing the strongest disease-relevant inflammatory-state-like response in this system.

Inflammatory and antiviral microglial states in human brains are enriched for LOAD genetic risk^25,26^. We performed stratified linkage disequilibrium score regression (S-LDSC) and observed significant enrichment of LOAD GWAS heritability in open chromatin from THP-1 macrophage states, along with primary microglia, macrophages, and iPSC/hESC-derived microglia, whereas no significant enrichment was detected in the neurons, HMC3 cells, and HEK293T cells tested (Extended Data Fig. 1f).

Collectively, these data indicate that THP-1 macrophages capture selected myeloid marker expression, activation-state signatures, and epigenomic features linked to LOAD, supporting their use as a practical model for studying LOAD-associated immune regulatory mechanisms.

### CREs harboring LOAD-associated variants exhibit context- and stimulation-dependent enhancer activity

To systematically evaluate common and rare LOAD variant effects on *cis*-regulatory activity, we designed STARR-seq-style MPRA constructs containing barcoded 227-bp candidate CREs carrying either reference or alternate alleles downstream of a minimal promoter, thereby enabling quantification of sequence-dependent enhancer activity (Fig. 2a). CREs were designed as open chromatin peak-centered (variants close to open chromatin peak summit), variant-centered (distal variants), or motif-shuffled (disrupting local TF motifs), providing multiple readouts of variant effects (Fig. 2a; Supplementary Table 3).

We assayed 1,600 CREs in cultured THP-1 macrophages and HMC3 cells, with untreated HEK293T cells included as a permissive non-immune comparison context. To capture immune-state-specific CRE activity, we assayed THP-1 macrophages and HMC3 cells under untreated, IFN-β-, IFN-γ-, and LPS+IFN-γ-stimulated conditions. In THP-1 macrophages, although MPRA plasmid-library nucleofection itself altered selected immune-related transcriptional programs, cytokine stimulation produced broader transcriptomic responses (Methods, Supplementary Fig. 3). To capture an *in vivo* neural regulatory context, we delivered the same MPRA library to the adult mouse brain using AAV-PHP.eB, a blood–brain barrier-penetrant capsid with strong CNS tropism and a predominantly neuronal transduction profile after systemic delivery^49,83,84^. We quantified CRE-driven enhancer activity from bulk cortex, hippocampus, and striatum to evaluate library activity across anatomically and cellularly distinct brain regions, including a well-characterized cortical context, the hippocampus as an AD-relevant region, and the striatum as a neuroanatomically distinct region enriched for GABAergic inhibitory neurons (Fig. 2b). Because AAV-PHP.eB transduction is predominantly neuronal, with limited glial and minimal microglial contributions^49,83–86^, but reporter RNA was collected from bulk tissue, we refer to this assay as a neuron-enriched neural-context assay.

We used MPRAnalyze^87^ to model CRE-driven reporter transcription by jointly analyzing DNA and RNA-derived cDNA counts across the 15 barcodes assigned to each LOAD-associated variant-containing candidate CRE. To obtain a conservative and consistently applicable activity criterion across assay contexts, the resulting activity estimates were standardized within each assay using a control-independent median absolute deviation (MAD) framework (Methods) derived from the broader within-assay activity distribution. Candidate CREs with a one-sided p.MAD < 0.10 were classified as active; this primary threshold was selected to balance sensitivity and stringency while remaining substantially more conservative than scrambled-control-based classification (Methods). Because the assay directly quantifies CRE-dependent reporter transcription, we hereafter refer to these reporter-based measurements as enhancer activity. Using the within-assay standardized MAD activity scores, we observed strong reproducibility among biological replicates (Pearson’s *r* ≈ 0.8–0.9 *in vitro* and *r* ≈ 0.6–0.7 *in vivo*; *P* < 0.01; Extended Data Fig. 3a). Enhancer-level DNA and RNA-derived cDNA read counts likewise showed strong within-context reproducibility (Supplementary Fig. 4). Activity profiles clustered by assay context, with strong correlations within immune or neural contexts and weaker correlations across them (Fig. 2c; Supplementary Table 4).

Given that the *in vivo* assay tested human regulatory sequences in mouse brain tissue, cross-species conservation was an important consideration. Previous comparative genomic studies have shown that tissue- and cell type-specific regulatory sequence features are partially conserved across mammals^88–90^. To obtain complementary evidence in a human neuronal context, we performed a pilot AAV-PHP.eB MPRA in human iPSC-derived excitatory neurons that expressed neuronal and excitatory/synaptic marker genes (Supplementary Fig. 5). Using the same within-assay MAD-based activity framework, mouse cortical positive-control sequences from Nguyen et al.^91^ showed higher reporter activity than negative controls in these human neurons, consistent with their activity pattern in the mouse brain tissue MPRA and supporting preservation of reporter activity for selected cortical CREs across species (Supplementary Fig. 6). Because barcode recovery was insufficient for robust enhancer-level and allele-specific inference, these pilot data were used as complementary cross-species evidence and were not included in the primary enhancer- or variant-effect analyses.

Approximately 18% of tested CREs were called significantly active in macrophages or brain tissue (Extended Data Fig. 3b, Supplementary Table 4). We first tested whether significant CREs (p.MAD < 0.1) differed in annotation composition from non-significant CREs (p.MAD ≥ 0.1) within each assay context. In THP-1 macrophages and HEK293T cells, significant and non-significant CREs showed significantly different distributions across the five predefined CRE categories (Immune, Neuronal, Glial, Broad, and Low-ATAC; χ^2^ test, FDR < 0.05; Fig. 2d), indicating that active MPRA hits were selectively enriched in specific CRE classes rather than sampled uniformly from the tested set. The corresponding comparison in brain tissue did not reach significance. We next compared the category composition of significant CRE hits across assay contexts. Among CREs with p.MAD < 0.1, THP-1 macrophage hits contained a higher fraction of Immune CREs than brain-tissue and HEK293T hits, whereas brain-tissue hits contained a higher fraction of Neuronal CREs and Low-ATAC sequences. HEK293T hits showed a prominent contribution from Broad CREs (Fig. 2d). The distribution of active hits reflected both the baseline representation of CRE categories in the library and category-specific activity across assay contexts. These categories capture endogenous accessibility patterns rather than absolute context specificity, and MPRA measures CRE activity outside the native genomic environment, allowing CREs from each category to respond to shared *trans*-regulatory factors across multiple assay contexts. Together, these results indicate that both the probability of a CRE being called active and the composition of significant hits varied across assay contexts.

Enhancer activity was also associated with sequence-encoded TF motif content. Elastic-net regression using aggregated FIMO motif-match scores as features provided a strong *in-sample* fit to enhancer activity in THP-1 macrophages and a more moderate fit in brain tissue (*R*^2^ = 0.90 and 0.48, respectively; permutation *P* = 0.001 for both), consistent with the greater variability of the *in vivo* brain measurements. Motif coefficients differed markedly between THP-1 macrophages and mouse brain tissue, consistent with context-biased regulatory logic. Among the top context-biased features, brain-biased coefficients included MEF2-, homeobox-, COE/EBF-, TBX-, SMAD-, Grainyhead- and TBP-related motifs, several of which are associated with neural and developmental regulation, whereas THP-1-biased coefficients included stimulus-responsive AP-1- and other bZIP-related motifs, together with FOX-, E2A-, SOX-, TEAD- and bHLH-ZIP-related families involved more broadly in lineage and cell-state regulation (Fig. 2e). At the level of individual CREs, we identified both broadly active and strongly context-biased enhancers; for example, a distal enhancer at the *BIN1* locus harboring rs13025765 and rs13025717 showed macrophage-biased activity, whereas a segment of the *NRN1* promoter harboring rs192104059 showed brain-tissue-biased activity (Fig. 2f).

Among the immune stimulation conditions, LPS+IFN-γ produced the most distinct enhancer responses. In THP-1 macrophages, 146 CREs (10%) showed significant activity changes after LPS+IFN-γ treatment (MPRAnalyze comparative analysis, FDR < 0.05), nearly half of which overlapped regions that were differentially accessible after LPS+IFN-γ stimulation, whereas IFN-β and IFN-γ alone affected only three and one CREs, respectively (Fig. 2g; Supplementary Table 5). Notably, multiple segments spanning exon 1 and intron 1 of *CTDP1* showed consistent increases in enhancer activity across all three stimulation conditions, indicating that this promoter-proximal region exhibits shared interferon-associated responsiveness, although *CTDP1* is not classically recognized as an immune gene. In HMC3 cells, 96 CRE sequences showed significant activity changes after 24 h of LPS+IFN-γ stimulation, compared with two after IFN-β stimulation and none after IFN-γ stimulation (FDR < 0.05; Supplementary Table 5).

LPS+IFN-γ predominantly reduced enhancer activity among responsive CREs in THP-1 macrophages, while increasing activity at only a smaller subset (Fig. 2g). Nearest-gene annotation linked LPS+IFN-γ-repressed CREs to genes enriched in immune-activation pathways, suggesting that these elements may help constrain excessive immune activation and further supporting the biological relevance of the identified enhancer responses (Fig. 2h; Supplementary Table 5). Elastic-nets trained independently on LPS+IFN-γ-treated and untreated enhancer activity showed that motif coefficients were broadly conserved (Pearson’s *r* = 0.91; Spearman’s *ρ* = 0.89), with selected motifs showing stimulation-dependent shifts. The FOSL2/AP-1- and ELK1-related motif coefficients shifted in the positive direction under LPS+IFN-γ, whereas the NRF2 (*NFE2L2*) motif coefficient became more negative, consistent with selective remodeling of motif-associated enhancer activity during inflammatory stimulation (Extended Data Fig. 3c). Together, these analyses identified LOAD-associated, variant-containing CREs with context- and stimulation-dependent enhancer activity across THP-1 macrophage and HMC3 cell states, a neuron-enriched brain-tissue context and the non-immune HEK293T comparison context.

### MPRA reveals context- and immune-state-dependent regulatory effects of LOAD-associated variants

Having established a context-resolved atlas of enhancer activity across immune and neural contexts, we next asked how LOAD variants modulate those activities. Screening 599 candidate variants represented by 855 allele-specific construct pairs, we identified expression-modulating variants (emVars; FDR < 0.05) in THP-1 macrophages and HMC3 cells under four immune states and *in vivo* in the cortex, hippocampus, and striatum (Extended Data Fig. 4a; Supplementary Table 6). Across all variants, 104 emVars were detected in both THP-1 macrophages and brain tissue, whereas 188 were detected in THP-1 but not brain tissue and 72 in brain tissue but not THP-1 (Extended Data Fig. 4b). Some variants in the latter two groups were also detected in HMC3 cells. These categories denote context-dependent detection rather than absence of an allelic effect in the other assay context. Across the four THP-1 activation states, allelic effects were moderately to strongly concordant. Pairwise SNP effect-size correlations ranged from 0.64 to 0.88, with the strongest concordance observed among untreated, IFN-γ-treated and IFN-β-treated THP-1 macrophages, whereas LPS+IFN-γ-treated THP-1 macrophages represented the most distinct condition (Extended Data Fig. 4d). Using FDR < 0.05, 229 SNPs showed significant allelic effects in at least one THP-1 condition, including 134 SNPs significant in at least two conditions, 93 in at least three conditions and 36 in all four conditions. Notably, all SNPs significant in multiple THP-1 conditions showed concordant effect directions across those conditions, indicating that allelic-effect directions were largely stable across THP-1 activation states (Extended Data Fig. 4e).

In contrast, among 79 tested elements exhibiting significant allelic effects in both THP-1 macrophages and brain tissue, and a significant THP-1 × brain context-by-allele interaction (corresponding to 77 unique SNPs), 40 elements (50.6%) displayed opposite allelic directions between the two contexts (Supplementary Fig. 7). More broadly, variant effects showed context-dependent patterns (Extended Data Fig. 4c). When aggregated across immune states or brain regions, emVars within active enhancers segregate into brain-biased, immune-biased and broadly shared clusters (Fig. 3a), highlighting substantial divergence in allelic effects between immune and neural contexts. Among 146 variants shared with the Cooper et al. MPRA dataset^30^, statistical significance and absolute allelic-effect size were positively correlated in the matched HEK293T context, but not between the Cooper et al. HEK293T assay and our brain-tissue assay, further supporting context dependence (Supplementary Fig. 8).

**Fig. 3.**
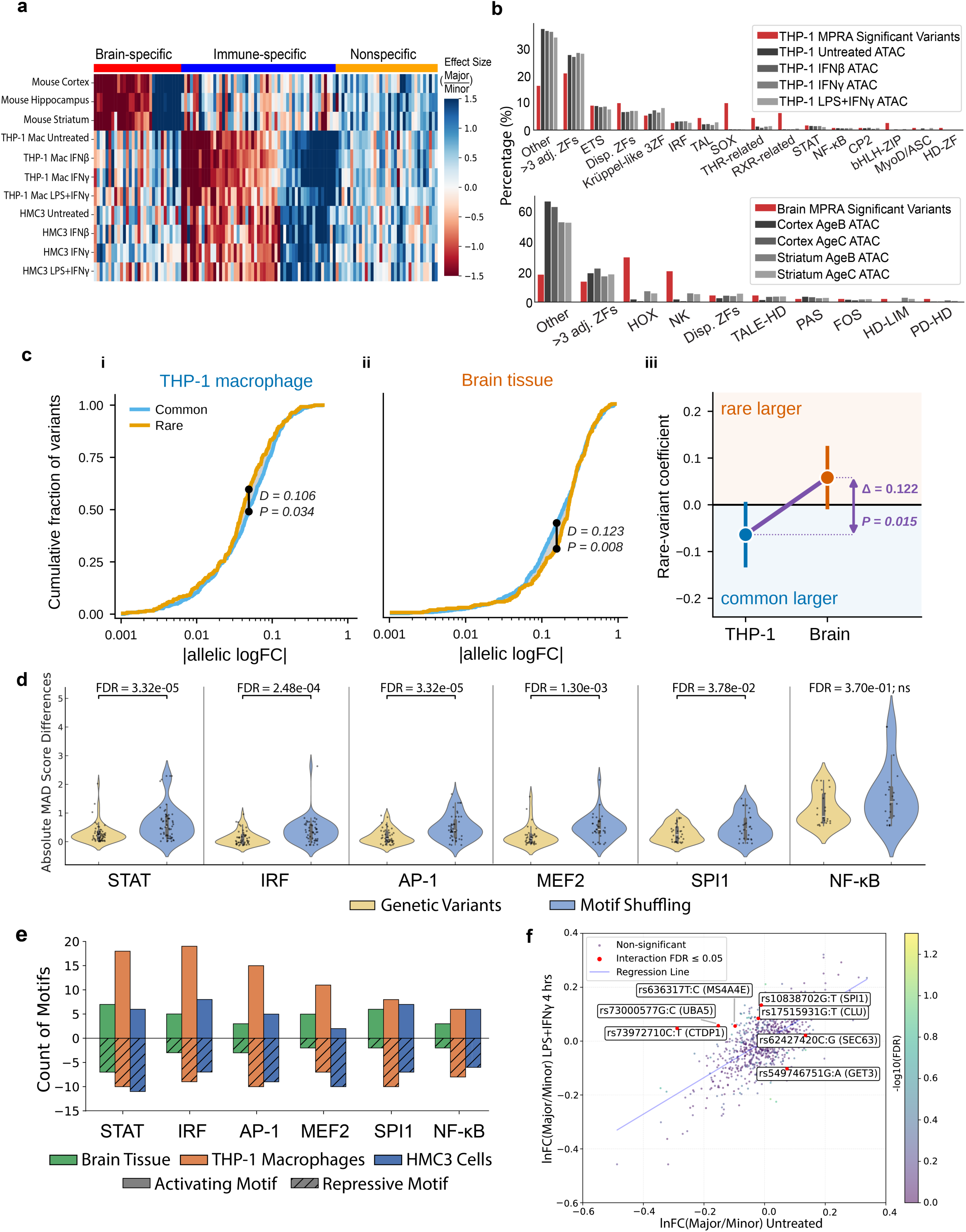
MPRA variant effects differ across allele-frequency classes, cellular contexts, immune states, and local TF-motif contexts. **(a)** Heatmap of allelic effect sizes for expression-modulating variants (emVars) across mouse brain regions, THP-1 macrophage conditions, and HMC3 conditions. Columns indicate emVars grouped into brain-specific, immune-specific, and nonspecific clusters, and rows indicate assay contexts. Color indicates the major-to-minor allele effect size. **(b)** Transcription factor (TF) motif-family composition of significant MPRA variants and open-chromatin peaks. Top, percentages of TF motif families associated with significant THP-1 MPRA variants and THP-1 ATAC-seq peaks from untreated, IFN-β-treated, IFN-γ-treated, and LPS + IFN-γ-treated conditions. Bottom, percentages of TF motif families associated with significant brain MPRA variants and ATAC-seq peaks from the cortex and striatum in the AgeB and AgeC groups. **(c)** Comparison of allelic effect sizes between common and rare LOAD-associated variants in THP-1 macrophages and brain tissue. Common, MAF ≥ 0.01 (n = 591 constructs); rare, MAF < 0.01 (n = 253 constructs). **i**, **ii**, Empirical cumulative distributions of absolute allelic log fold changes for common (blue) and rare (orange) variants in THP-1 macrophages (**i**) and brain tissue (**ii**); the x-axis is on a log₁₀ scale with untransformed tick labels. Black points indicate the maximum separation between the distributions; *D* and *P* values are from two-sided Kolmogorov–Smirnov tests. **iii**, Rare-variant coefficients from ordinary least-squares models fitted separately in each context (n = 844 constructs assayed in both contexts), adjusting for mean allele reporter activity, ATAC-defined variant category, construct design, GC content and distance to the transcription start site (Extended Data Fig. 5e). Positive coefficients indicate larger allelic effects for rare variants, whereas negative coefficients indicate larger effects for common variants. Points indicate coefficient estimates and error bars indicate 95% confidence intervals. The purple arrow spans the difference between contexts (Δ = 0.122, 95% CI 0.024 to 0.220, *P* = 0.015), estimated as the allele-frequency × context interaction in a pooled fully stratified model with construct-clustered standard errors. All *P* values are two-sided. **(d)** Violin plots comparing absolute MAD-score differences between genetic-variant constructs and motif-shuffled constructs across STAT, IRF, AP-1, MEF2, SPI1, and NF-κB motif contexts in THP-1 macrophages. Points indicate individual constructs, and FDR-adjusted *P* values are shown above each comparison; ns, not significant. **(e)** Numbers of activating and repressive motif-context effects across STAT, IRF, AP-1, MEF2, SPI1, and NF-κB motif groups in brain tissue, THP-1 macrophages, and HMC3 cells. Solid bars above zero indicate activating motif effects, whereas hatched bars below zero indicate repressive motif effects. **(f)** Comparison of allelic effects in untreated and LPS + IFN-γ-treated THP-1 macrophages. The x-axis shows the natural-log fold change between the major- and minor-allele constructs in untreated THP-1 macrophages, and the y-axis shows the corresponding natural-log fold change after 4 h of LPS + IFN-γ stimulation. Point color indicates −log10(FDR) for the treatment-by-allele interaction. Red points indicate variants with an interaction FDR ≤ 0.05; all other points indicate nonsignificant interactions; the line indicates the linear regression fit. Labels identify selected emVars with treatment-dependent allelic effects.

This divergence prompted us to examine whether common and rare LOAD risk variants differ in how they act across cellular contexts. Rare variants came almost entirely from the WGS family-based study, whereas common variants came from GWAS and fine-mapping resources together with the same WGS study, in which association was tested within genomic regions and both common and rare variants within those regions entered our library (Methods).

Comparing the two classes within each context revealed a reversal. Common variants were shifted towards larger effects in THP-1 macrophages (Kolmogorov–Smirnov *D* = 0.106, *P* = 0.034), whereas rare variants were shifted towards larger effects in brain tissue (*D* = 0.123, *P* = 0.008; Fig. 3c i,ii). A frequency × context interaction, adjusting for mean allele reporter activity, ATAC-based CRE category, construct design, GC content and distance to TSS, placed this reversal on a formal footing (Δ = 0.122, 95% CI 0.024–0.220, *P* = 0.015, construct-clustered standard errors; n = 844 constructs assayed in both contexts; Fig. 3c iii). The WGS study provides an internal control that sharpens this result. Because it contributed both common (n = 234) and rare (n = 225) variant constructs drawn from a single set of nominated genomic regions, and because 42 of its 64 loci contain both classes, the two groups can be compared without any difference in source or genomic location. Within this study the interaction is not merely preserved but stronger (Δ = 0.162, 95% CI 0.047–0.276, *P* = 0.006), and it survives restriction to the 42 shared loci with locus fixed effects, where common and rare variants are compared within the same locus (Δ = 0.148, 95% CI 0.037–0.259, *P* = 0.009; Extended Data Fig. 5d). Within this selected set of AD-associated regions, allele-frequency class was associated with the assay context in which variants showed larger reporter effects, independently of several measured features of study origin and CRE annotation.

The interaction was stable across method and specification, holding without covariates, under construct-level bootstrap and label permutation, and within each ATAC-defined CRE category; the direction of the contrast was preserved at every common-group MAF threshold from 0.01 to 0.40, and the two frequency classes overlapped immune open-chromatin peaks at similar rates (85% versus 81%), differing mainly in neuronal peak overlap (32% versus 49%) (Extended Data Fig. 5a–e). Modelling MAF as a continuous variable did not detect a significant MAF × context interaction (β = −0.034, P = 0.13), providing no evidence for a monotonic linear gradient across allele frequency.

We next oriented MPRA allelic effects according to the Bellenguez β-defined risk allele for GWAS variants and the sign of the locus-level W statistic for WGS-derived variants (Methods). WGS-derived allelic effects oriented by the locus-level W statistic (Supplementary Table 1) showed a significant aggregate shift toward reduced reporter transcription in brain tissue, whereas THP-1 macrophages did not show this pattern (Extended Data Fig. 4f). Because W is defined at the locus level, this analysis does not assign genetic-risk direction to individual alleles. In contrast, Bellenguez^5^ β-aligned effects for genome-wide significant GWAS variants (*P* < 5 × 10⁻^8^) were not directionally shifted in either context (Extended Data Fig. 4f).

To investigate the regulatory basis of this context-dependent divergence, we next examined TF-motif enrichment near emVars, comparing it directly to motif enrichment within open chromatin regions in THP-1 macrophages and brain tissue. Motifs enriched in open chromatin were also preferentially enriched near emVars within the same assay context, with consistent representation across TF families. Brain-active emVars were predominantly associated with motifs from TF families involved in neural development and function, including TALE-type homeodomain, HOX-related, NK-related, and Fos-related families. In contrast, emVars active in THP-1 macrophages were enriched for immune-related TF families such as ETS-related, IRF, STAT, and NF-κB. Together, these analyses identify context-biased motif environments associated with the regulatory effects of LOAD variants (Fig. 3b).

We next directly tested whether the TF-motif contexts surrounding these variants functionally modulate enhancer activity. Given that LOAD variants are often predicted to disrupt immune-related TF motifs, we analyzed a set of motif-shuffled enhancers in which a 25-bp window surrounding each variant was fully shuffled to disrupt the predicted immune-related TF-motif context, a design we refer to as motif-context shuffling. Because single-nucleotide variants often produce more limited sequence perturbations, we hypothesized that broader disruption of the surrounding motif context would elicit larger transcriptional shifts. In both THP-1 macrophages and HMC3 cells, most motif-context shuffling events caused significant changes in enhancer activity (FDR < 0.05, Supplementary Table 7) and produced markedly larger changes in reporter expression than single-nucleotide substitutions, with the strongest effects observed at motif contexts containing STAT, MEF2, IRF, AP-1, and SPI1 motifs (Fig. 3d). Motif-context shuffling either increased or decreased transcription, though certain TF-motif contexts showed consistent directional biases. For example, IRF-, STAT-, and AP-1-related motif contexts preferentially decreased transcription upon disruption in THP-1 macrophages, consistent with an activating contribution of these local sequence contexts (Fig. 3e). These findings show that many assayed variants reside within functionally active TF-motif contexts.

TFs involved in immune-signaling pathways, such as the interferon response, are often transiently activated; thus, variants altering these TF-motif contexts may exert regulatory effects only under specific immune conditions. To explore this possibility, we analyzed variant effects separately under IFN-β, IFN-γ, and LPS+IFN-γ stimulation in THP-1 macrophages and HMC3 cells. We detected significant state-specific emVars (FDR < 0.05) in all conditions except IFN-β-stimulated HMC3 cells. Under LPS+IFN-γ stimulation, we identified seven state-specific emVars in THP-1 macrophages and four in HMC3 cells (Fig. 3f, Supplementary Table 8). Notably, rs10838702-T (*SPI1*) and rs17515931-T (*CLU*) reduced transcription exclusively under hyperinflammatory conditions, whereas rs62427420-G (*SEC63*) reduced transcription only in untreated cells. Several variants exhibited more complex patterns: rs73972710-T (*CTDP1*), rs73000577-C (*UBA5*), and rs636317-C (*MS4A4E*) increased transcription in both untreated and stimulated states, but showed stronger activation upon stimulation, whereas rs549746751-A (*GET3*) exerted opposing effects depending on the immune state. These examples are consistent with state-dependent regulatory mechanisms in which variants alter the activity of conditionally available TFs, modulate motif-dependent activity to different degrees across states, or affect sequences interpreted by different TF environments under distinct immune conditions.

In summary, our MPRA identified context-biased and immune-state-specific effects of LOAD-associated variants on enhancer activity, with regulatory effects associated with local TF-motif contexts and differing between common GWAS-derived and rare WGS-derived variant groups. WGS-derived risk-direction alleles additionally showed a directional bias toward reduced reporter transcription in brain tissue.

### Deep-learning models recover context-specific regulatory sequence features and prioritize LOAD-associated variants

To characterize sequence features underlying context-dependent chromatin accessibility and facilitate interpretation of MPRA variant effects, we trained separate CNN regression models on ATAC-seq data from multiple cell types, including stimulated and unstimulated THP-1 macrophages, hESC/iPSC-derived microglia, THP-1 monocytes, human brain-region-derived neurons, mouse cortex and striatum, human astrocytes and oligodendrocytes, and HEK293(T) cells (Methods). These models accurately predicted chromatin accessibility in held-out data from matched or related cell types (Spearman’s ρ ≈ 0.8), whereas predictions across divergent cell types yielded moderate performance (Spearman’s ρ ≈ 0.5), demonstrating generalization to unseen sequences while preserving biologically meaningful differences among cellular contexts (Fig. 4a). Notably, the cross-prediction matrix revealed a clear separation between neural and immune/myeloid regulatory programs. Models trained on human neurons or mouse brain tissue generalized preferentially across neural datasets and showed substantially weaker performance on immune/myeloid datasets. Human neuronal models also showed stronger cross-prediction with mouse cortex and striatum than models trained on human astrocytes or oligodendrocytes, supporting a predominantly neuronal rather than broadly glial regulatory profile of the mouse brain datasets. By contrast, iPSC- and hESC-derived microglia showed prediction patterns more similar to THP-1 and other myeloid models. Within the neural datasets, the models showed substantial cross-species generalization with regionally coherent patterns: human hippocampal and DLPFC neuronal models cross-predicted mouse cortex accessibility, whereas the human putamen model showed stronger correspondence with mouse striatum. These results indicate partial conservation of neural regulatory sequence features across species and support the use of mouse brain as an *in vivo* context for assaying human CREs (Fig. 4a).

**Fig. 4.**
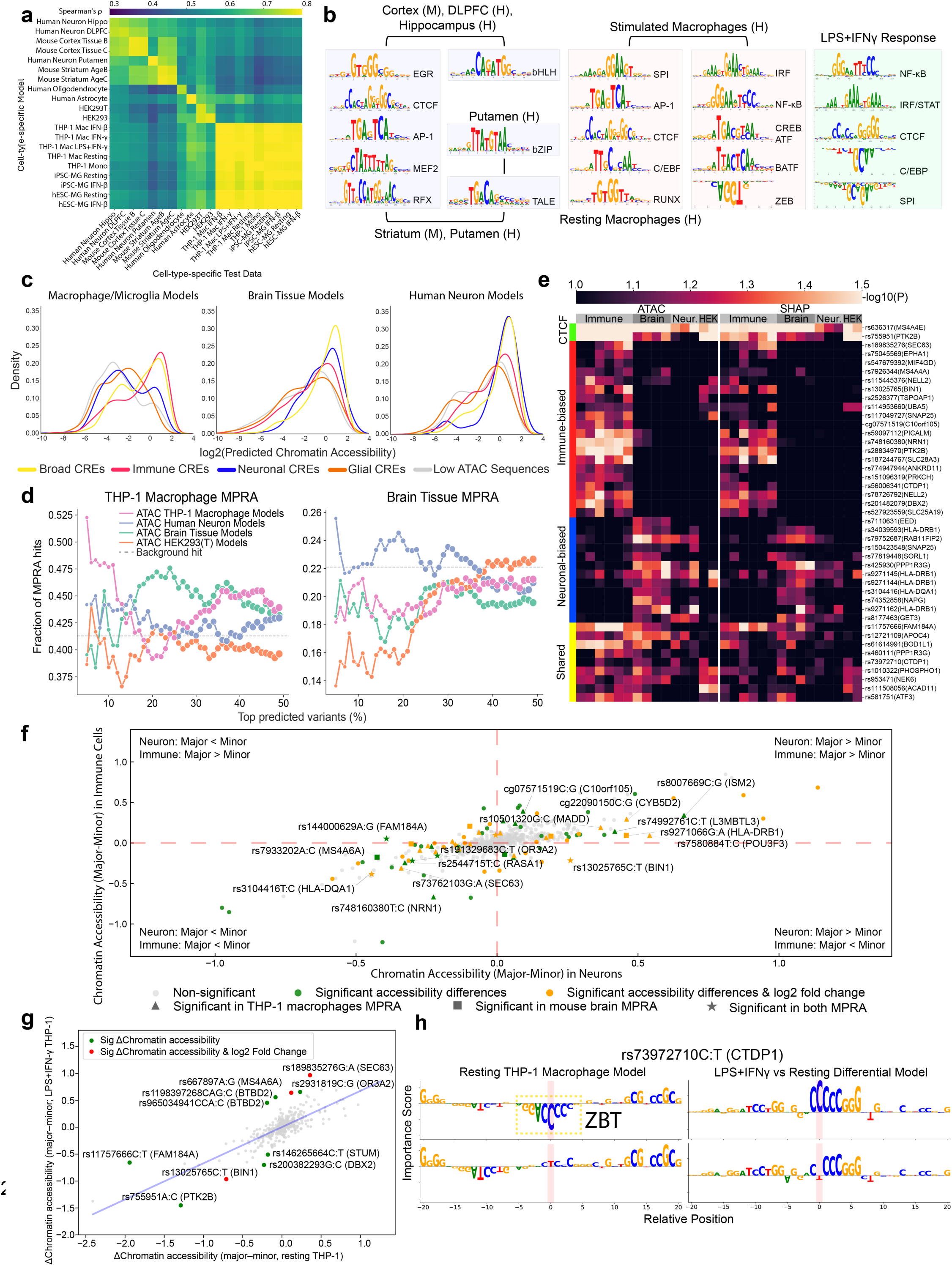
Accessibility-trained sequence models prioritize LOAD-associated variants and nominate cell-type- and state-associated regulatory features. **(a)** Spearman correlation heatmap comparing model-predicted and observed chromatin accessibility across cell-type- and state-specific held-out test datasets. Rows indicate trained models, and columns indicate held-out test datasets. Color indicates the Spearman correlation coefficient (ρ). **(b)** Representative transcription factor (TF) motifs identified by TF-MoDISco from context-specific sequence-to-function models. Motifs are shown for mouse and human brain-region models, resting and stimulated macrophage models, and the lipopolysaccharide plus interferon-γ (LPS + IFN-γ) response model. DLPFC denotes dorsolateral prefrontal cortex; M and H denote mouse and human models, respectively. **(c)** Density distributions of log₂-transformed predicted chromatin accessibility across candidate *cis*-regulatory element (CRE) categories in macrophage/microglia, brain-tissue, and human-neuron models. CRE categories include Broad, Immune, Neuronal, Glial, and Low ATAC sequences. **(d)** Fraction of massively parallel reporter assay (MPRA) hits among candidate variants ranked by predictions from different assay for transposase-accessible chromatin using sequencing (ATAC-seq) model groups. Left, enrichment of THP-1 macrophage MPRA hits. Right, enrichment of mouse brain-tissue MPRA hits. The x-axis indicates the percentage of top-ranked candidate variants, and the y-axis indicates the fraction identified as MPRA hits. Lines represent ATAC-seq models trained on THP-1 macrophages, human neurons, brain tissue, or HEK293T cells. Dashed horizontal lines indicate the background MPRA hit rate. **(e)** Heatmap of −log₁₀(*P*) values for variant-level allelic effects estimated from the null distribution of ATAC-model predictions and SHapley Additive exPlanations (SHAP) attribution scores. Columns represent immune, brain-tissue, neuronal, and HEK293T model groups. Rows indicate variants grouped according to CTCF-associated, immune-biased, neuronal-biased, or shared MPRA effects. **(f)** Comparison of predicted major-minus-minor allelic chromatin-accessibility differences in neuronal and immune models. The x-axis indicates the predicted allelic difference in neuronal models, and the y-axis indicates the corresponding difference in immune-cell models. Gray points indicate nonsignificant variants, green points indicate variants with significant predicted accessibility differences, and orange points indicate variants with significant predicted accessibility differences and log₂ fold-change differences. Triangles indicate variants significant by THP-1 macrophage MPRA, squares indicate variants significant by mouse brain MPRA, and stars indicate variants significant in both MPRA contexts. Dashed lines indicate zero allelic difference. Labels identify selected variants and nearby genes. **(g)** Comparison of predicted allelic chromatin-accessibility differences in resting and LPS + IFN-γ-treated THP-1 macrophage models. The x-axis indicates the major-minus-minor allelic difference in the resting model, and the y-axis indicates the corresponding difference in the LPS + IFN-γ-treated model. Gray points indicate nonsignificant variants, green points indicate variants with significant predicted accessibility differences, and red points indicate variants with significant accessibility differences and log₂ fold-change differences. The blue line indicates the linear regression fit. Labels identify selected variants and nearby genes. **(h)** Allele-specific, base-resolution importance-score logos for rs73972710:T near *CTDP1* in the resting THP-1 macrophage model and the LPS + IFN-γ-versus-resting differential model. The x-axis indicates position relative to the variant, and the y-axis indicates the predicted importance score. Paired tracks show allele-specific importance profiles. Vertical pink shading marks the variant position, and the dashed yellow box highlights the local ZBT motif identified in the resting-state model.

Using SHAP^92^-based importance scores and TF-MoDISco^93^, which identifies recurring sequence patterns contributing to model predictions, we identified context-associated TF-motif features learned by the models. Immune-related motifs, including SPI-, C/EBP-and RUNX-related motifs, were enriched in macrophage and microglia models, whereas neuron-associated motifs, including EGR-, MEF2- and RFX-related motifs, were enriched in neuron and bulk-brain models (Fig. 4b). Models trained on stimulated immune cells additionally recovered IRF-, NF-κB-, CREB/ATF- and BATF-related motifs with positive contributions to predicted accessibility, whereas ZEB-related motifs showed negative contributions in interferon-treated models.

To resolve sequence features associated with hyperinflammatory chromatin remodeling, we trained a model specifically on regions that were differentially accessible between LPS+IFN-γ-treated and untreated THP-1 macrophages (Supplementary Fig. 9). Model interpretation identified NF-κB-, IRF/STAT- and CTCF-related motifs as features associated with chromatin opening, whereas SPI- and C/EBP-related motifs were unexpectedly associated with chromatin closing (Fig. 4b). Previous single-nucleus multiomic work suggested that SPI1 (PU.1) can function as a transcriptional repressor in late-stage LOAD microglia^94^, a disease context marked by neuroinflammation^95^, raising the possibility that SPI-family motif effects are context dependent and may contribute to repression under strong inflammatory stimulation. Together, these results indicate that the models recovered interpretable regulatory sequence features associated with cell-type- and state-dependent patterns of chromatin accessibility.

We next applied these independently trained sequence models to LOAD variant-containing MPRA CREs to assess whether their predicted accessibility patterns were concordant with the ATAC-seq-based CRE annotations. When grouped post hoc by ATAC-defined CRE categories, immune-cell models assigned relatively higher predicted accessibility to Immune and Broad CREs and lower scores to Neuronal and Glial CREs and Low-ATAC sequences, whereas mouse-brain-tissue and human-neuron models assigned higher scores to Neuronal and Broad CREs than to Glial CREs and Low-ATAC sequences, with intermediate predictions for some Immune CREs (Fig. 4c). The models also recovered motif features consistent with TF motifs enriched in the matched ATAC-seq datasets, including context-associated and LPS+IFN-γ-responsive features (Extended Data Fig. 3d,e). Together, these findings indicate that sequence features learned from endogenous ATAC-seq retained expected immune-versus-neural and stimulation-associated differences when applied to the MPRA library sequences.

We then applied *in silico* mutagenesis to assess predicted allelic effects of LOAD variants (Supplementary Table 9), using GC-matched non-peak control sequences to define null distributions for predicted accessibility differences and SHAP contrasts, allowing cross-context comparisons, and Monte Carlo dropout to estimate uncertainty in individual predictions. Across both accessibility-difference- and SHAP-based rankings, context-matched models preferentially prioritized variants with experimentally detected allelic effects, indicating context-specific alignment between model predictions and MPRA outcomes. THP-1 macrophage models showed the strongest enrichment for THP-1 macrophage MPRA hits, whereas human neuronal models most effectively prioritized brain-tissue MPRA hits, showing stronger enrichment than astrocyte, oligodendrocyte, THP-1 macrophage, HEK293(T), and bulk mouse-brain models, particularly among the most highly ranked variants (Fig. 4d; Extended Data Fig. 6a,b).

Human neuronal models prioritized brain-tissue MPRA hits more strongly than glial or myeloid models. The stronger prioritization by human neuronal than bulk mouse-brain models may reflect the predominantly neuronal transduction profile of AAV-PHP.eB and the human origin of the assayed CREs, whereas bulk mouse-brain ATAC-seq integrates accessibility signals across heterogeneous neuronal and non-neuronal cell populations. By integrating predicted accessibility changes and SHAP importance scores, which were significantly correlated (Spearman’s ρ = 0.8; Extended Data Fig. 6c), the models identified variants that exhibit effects across cell types as well as variants with effects biased toward the immune or neural contexts (P < 0.1; Fig. 4e). Among the strongest outliers were rs636317 (*MS4A4E*) and rs755951 (*PTK2B*), both of which alter CTCF motifs. Immune-biased variants were identified near genes implicated in microglial biology, including *EPHA1, MS4A4A, BIN1, PICALM* and *PTK2B*, while brain-biased hits included variants near *SORL1*, which is involved in APP trafficking and processing, and near *SNAP25* and *NAPG*, which participate in synaptic or vesicular trafficking.

Interestingly, some variants showed consistent directional effects across cell types, while others displayed opposing effects (Extended Data Fig. 6d). These results show that the models prioritize MPRA-supported regulatory variants and recover context-dependent effect patterns.

To identify context-dependent predicted allelic effects, we used linear regression with cell type as an explanatory variable to test whether CNN-predicted allelic effects differed between human immune and neuronal contexts. (Supplementary Fig. 10). This analysis highlighted two major classes of variants: those with the same predicted direction but different effect magnitudes across contexts, and those with opposite predicted effects between immune and neuronal models (Fig. 4f; Extended Data Fig. 6e). Notably, several common variants at the immune-associated *HLA-DRB1* and *HLA-DQA1* loci showed stronger predicted effects in neuronal than immune models (Fig. 4e,f). MPRA measurements showed that HLA-region variants—including rs3104416, rs9271162, rs9271144 and rs9271145—had larger normalized absolute allelic effects in brain tissue than in THP-1 macrophages (Extended Data Fig. 4g). In neuronal models, the minor G allele of rs9271162 near *HLA-DRB1* was predicted to weaken a TALE-type homeodomain motif, the minor T allele of rs9270833 near *HLA-DRB1* was associated with reduced MEF2-related sequence attribution, and the minor C allele of rs3104416 near *HLA-DQA1* was predicted to create or strengthen an EGR-related motif (Extended Data Fig. 6f). Several context-biased predictions also corresponded to MPRA-defined emVars. For example, rs13025765 near *BIN1* was detected as an emVar in both THP-1 macrophages and brain tissue, with opposing allelic effects across the two assays.

Consistent with this pattern, the minor T allele was predicted to increase accessibility in immune models but decrease accessibility in neural models. Together, these findings support the use of accessibility-trained sequence models to prioritize and interpret context-dependent allelic effects and to generate hypotheses about the sequence features associated with those effects.

We next explored immune-state-dependent predicted allelic effects using models trained on chromatin-accessibility profiles from LPS+IFN-γ-, IFN-γ- and IFN-β-treated or untreated THP-1 macrophages (Fig. 4g; Extended Data Fig. 6g). Several variants showed state-dependent predictions, including rs13025765 near *BIN1*, which showed a larger predicted allelic effect under LPS+IFN-γ stimulation than under the untreated condition, and rs189835276 near *SEC63*, which showed larger predicted effects under all three stimulated conditions. A model trained on regions differentially accessible between LPS+IFN-γ-treated and untreated THP-1 macrophages provided additional sequence-level interpretation (Extended Data Fig. 3d, Supplementary Table 9).

For rs73972710 near *CTDP1*, the minor T allele increased reporter transcription relative to the major C allele in the untreated MPRA condition. Consistent with this effect, the model trained on untreated THP-1 macrophages predicted that the C allele preserved a ZBT/ZBTB-associated repressive sequence feature that was weakened by the T allele. In the LPS+IFN-γ-versus-untreated differential model, however, the T allele showed a smaller predicted inducible effect than the C allele, potentially offsetting its effect under the untreated condition and consistent with the attenuated MPRA allelic difference after stimulation (Fig. 4h; Fig. 3f).

Collectively, these analyses show that context-specific accessibility models recover interpretable cell-type- and immune-state-associated regulatory sequence features, prioritize LOAD-associated variants with context-biased predicted allelic effects, and generate sequence-level hypotheses that align with selected MPRA measurements. Accessibility-trained models therefore provide a complementary approach for prioritizing candidate regulatory variants and investigating sequence features associated with context-specific effects.

### Integrated analyses prioritize context-dependent regulatory variants at LOAD-associated loci

To integrate experimental and sequence-model evidence at the locus level, we examined representative LOAD-associated GWAS- and WGS-derived regions containing multiple emVars detected by MPRA in THP-1 macrophages and neuron-enriched brain tissue (Fig. 5a). A subset of these emVars was additionally supported by at least one relevant sequence model that predicted a significant difference in chromatin accessibility between the major and minor alleles. Within individual regions, some minor alleles increased reporter transcription, whereas others decreased it, and a small number showed construct-discordant effects, indicating context- and sequence-dependent allelic regulatory activity (Fig. 5a). The WGS-derived *SEC63–OSTM1* and *NELL2–DBX2* regions contained multiple assayed emVars with effects detected in both THP-1 macrophages and brain tissue, whereas a subset of *NELL2* intronic variants was detected only in brain tissue under the conditions tested (Extended Data Fig. 8).

**Fig. 5.**
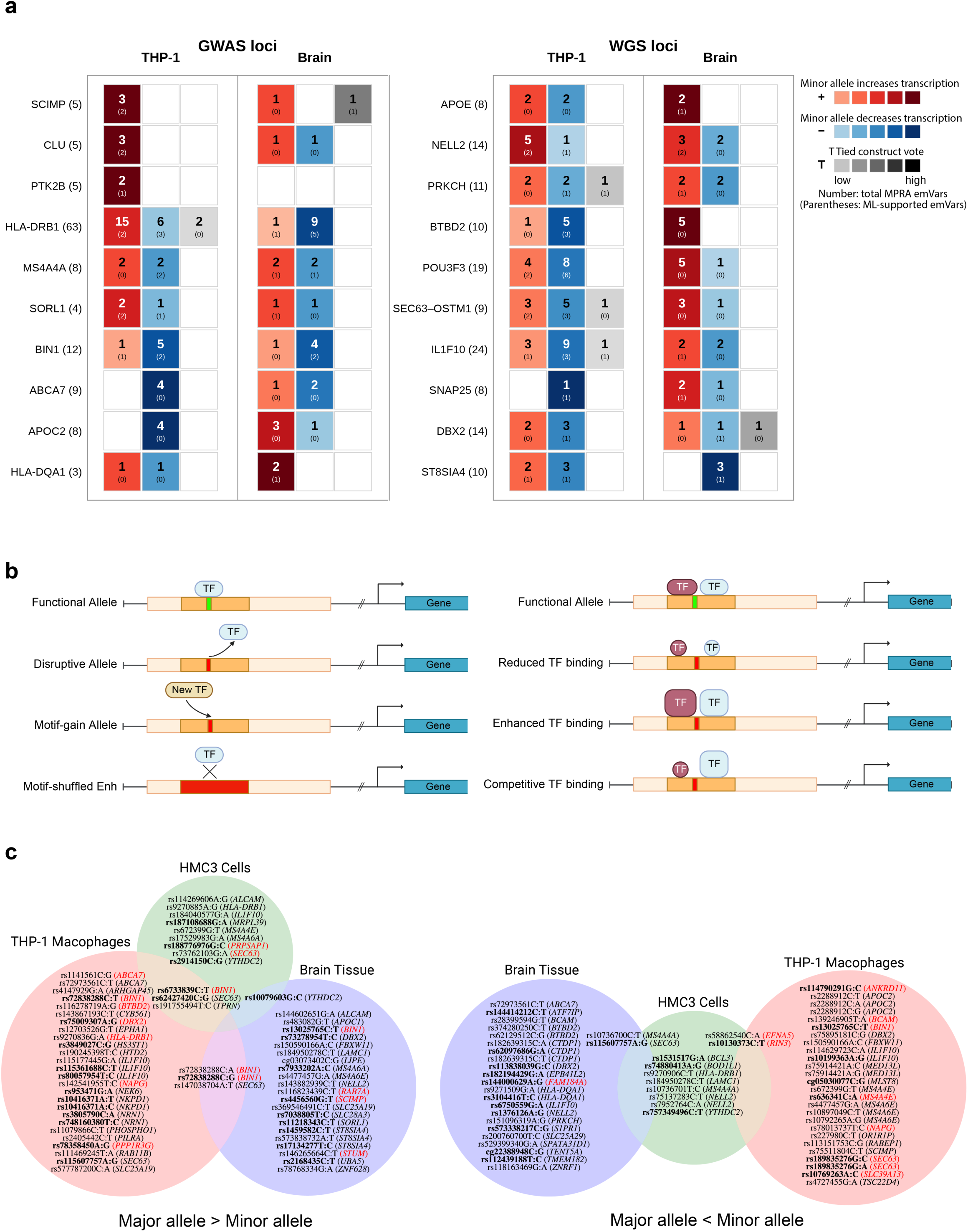
Integrated prioritization of candidate regulatory variants across LOAD-associated GWAS and WGS loci. **(a)** Locus-level summary of expression-modulating variants (emVars) identified by MPRA in THP-1 macrophages and mouse brain tissue. Loci are grouped according to whether they were identified by genome-wide association studies (GWAS) or whole-genome sequencing (WGS). Numbers in parentheses after locus names indicate the number of candidate variants assayed at each locus. Red cells indicate emVars for which the minor allele increased transcription relative to the major allele, whereas blue cells indicate emVars for which the minor allele decreased transcription. Gray cells indicate tied construct-level votes. Color intensity reflects the strength of the construct-level vote. Numbers within cells indicate the total number of MPRA emVars, and numbers in parentheses indicate the subset supported by machine-learning (ML) predictions. **(b)** Proposed mechanisms through which regulatory variants alter transcription factor (TF) binding and transcriptional output. Variants may disrupt an existing TF motif, create a new motif, reduce or enhance TF binding, or alter competitive binding between TFs. Motif-shuffled enhancer constructs disrupt the local motif context and were used to assess the contribution of local TF binding to enhancer activity. **(c)** Context-specific distribution of candidate functional regulatory variants identified in THP-1 macrophages, HMC3 cells, and mouse brain tissue. Variants are separated according to whether the major allele exhibited higher transcriptional activity than the minor allele (left) or lower transcriptional activity than the minor allele (right). Candidate variants were required to show a significant allelic effect in the MPRAnalyze comparative analysis (FDR < 0.05) and significant enhancer activity for at least one allele in the one-sided MPRAnalyze quantitative analysis (*P*.MAD < 0.1). Variant labels shown in red indicate support from motif-shuffling analysis, defined by a significant difference between an allelic construct and its corresponding motif-shuffled construct in the MPRAnalyze comparative analysis (FDR < 0.05). Bold labels indicate variants predicted by ML models to significantly alter chromatin accessibility. Variants identified in THP-1 macrophages or HMC3 cells were evaluated using THP-1 macrophage models, whereas variants identified in mouse brain tissue were evaluated using human neuronal models. ML-supported variants met the following criteria in at least one corresponding model: Benjamini–Hochberg-adjusted FDR ≤ 0.05 based on a two-sided Wilcoxon signed-rank test of Bayesian predictions generated using Monte Carlo dropout; either an absolute predicted natural-log fold change in accessibility between alleles > 0.1 or an absolute allelic difference in predicted accessibility score > 0.5; and a predicted accessibility score ≥ 1 for the allele with higher predicted accessibility.

Variants assigned to the *BIN1* region also showed significant allelic effects in both assay contexts (Fig. 5a). We further detected noncoding emVars assigned to the *APOE* region and *APOC2* across immune and neural contexts. Together, these results show that selected variants within individual LOAD-associated regions can produce distinct effects across assay contexts, with a subset also captured by independently trained accessibility models.

To examine regulatory-effect direction relative to genetic association, we oriented common-variant effects to the Bellenguez β-defined risk allele and WGS-derived effects according to the sign of the locus-level W statistic (Extended Data Fig. 7). Among the assayed variants with available directional information, β-aligned risk-allele effects assigned to *MS4A4A* and *CLU* were predominantly transcription-decreasing in THP-1 macrophages. Variants assigned to *BIN1* showed both transcription-increasing and transcription-decreasing effects, with an overall bias toward decreased reporter transcription in THP-1 macrophages, whereas their effects were evenly distributed between the two directions in brain tissue. Variants assigned to *CLU* showed opposite predominant directions across contexts, decreasing reporter transcription in THP-1 macrophages and increasing it in brain tissue. Among the WGS-derived variants, W-aligned effects assigned to *APOE*, *POU3F3* and *BTBD2* also showed context-dependent directionality, with opposite predominant directions between the THP-1 macrophage and brain assays. These selected examples illustrate that association-aligned enhancer effects among the assayed variants can differ between immune and neural contexts.

Integrating multidimensional measurements of genetic variant effects, sequence-level regulatory activity, and motif context function, we synthesized a conceptual framework of regulatory variant actions (Fig. 5b). Variants can (i) destroy or create a core TF site, (ii) fine-tune the strength of adjacent TF motifs, or (iii) modify the balance among neighboring activating and repressive sequence features; motif shuffling represents the extreme case of class (i). Guided by this framework, we prioritized candidate functional regulatory variants that met stringent criteria, including significant enhancer activity, allele-specific transcriptional differences, and, where applicable, disruption of a motif context (Fig. 5c, Supplementary Table 10). A subset of these variants also showed significant predicted allelic differences in chromatin accessibility in at least one relevant sequence model.

To explore candidate enhancer–promoter relationships beyond nearest-gene annotation, we applied the Activity-by-Contact (ABC) model^96^ to distal emVar-containing CREs in THP-1 macrophage contexts^97^. ABC nominated non-nearest candidate target promoters for 7 of 10 unique CREs, indicating that linear genomic proximity alone may not capture all potential enhancer–promoter relationships (Supplementary Fig. 11 and Supplementary Table 11).

The integrated MPRA–machine-learning framework prioritized context-dependent candidate regulatory variants and generated sequence-level hypotheses about their regulatory mechanisms. We next examined representative loci using model interpretation and locus-specific analyses, followed by CRISPRi perturbation of a prioritized enhancer at the *SEC63–OSTM1* locus in THP-1 macrophages to assess its endogenous transcriptional consequences.

### Mechanistic interpretation of context-dependent regulatory variants at the *BIN1*, *MS4A*, and *SEC63–OSTM1* loci

Among the prioritized candidate regulatory variants, we selected three LOAD-associated regions containing multiple variants with divergent reporter effects for detailed analysis (Fig. 6a). These include (i) the *BIN1* regulatory region containing two adjacent enhancer CREs; (ii) an enhancer within the critical *MS4A* locus, where several gene products have been implicated in regulating *TREM2* expression and microglial state^98,99^; and (iii) a newly identified enhancer overlapping a rare variant cluster at the *SEC63-OSTM1* locus. We compared allelic effects on reporter transcription with context-specific accessibility predictions and model-derived motif annotations across immune and neural assay contexts.

**Fig. 6.**
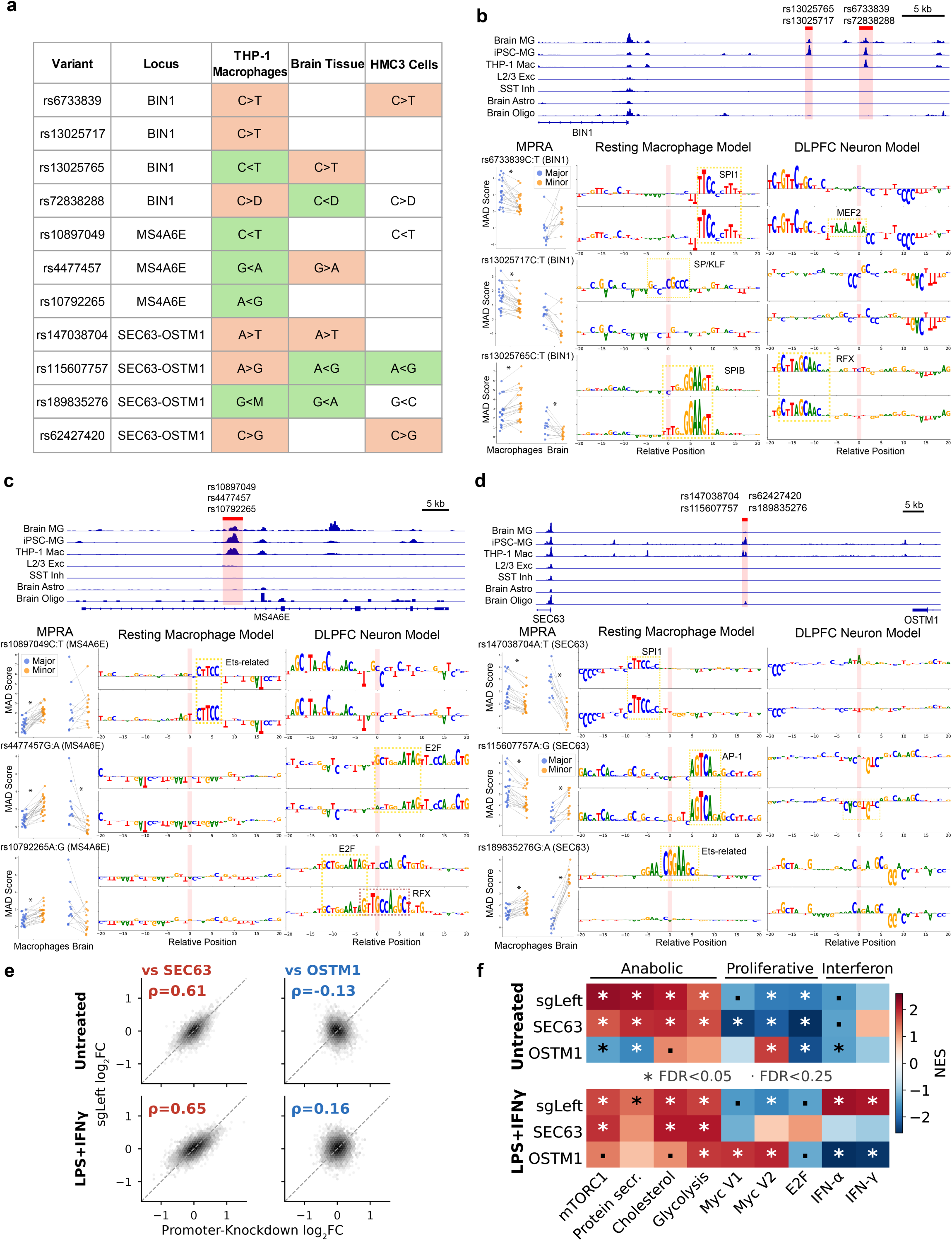
Context-dependent mechanistic analysis of candidate regulatory variants at the BIN1, MS4A6E, and SEC63–OSTM1 loci. **(a)** Summary of selected expression-modulating variants at the *BIN1*, *MS4A6E*, and *SEC63–OSTM1* loci across THP-1 macrophages, mouse brain tissue, and HMC3 cells. For each entry, the first symbol denotes the major allele and the second denotes the minor allele or, for multiallelic variants, a group of non-major alleles. IUPAC nucleotide ambiguity codes are used to collectively represent multiple non-major alleles: D denotes A, G, or T, and M denotes A or C. The inequality sign indicates the direction of allelic transcriptional activity measured by massively parallel reporter assay (MPRA); for example, C > T indicates that the major-allele C construct exhibited greater transcriptional activity than the minor-allele T construct, whereas C < T indicates greater activity of the minor-allele T construct. Orange cells indicate significant effects with greater activity of the major allele, whereas green cells indicate significant effects with greater activity of the minor allele. Uncolored entries indicate the estimated allelic direction without a significant allelic difference, and blank cells indicate contexts without a statistically significant effect. **(b)** Chromatin accessibility, MPRA allelic activity, and model-based sequence interpretation for selected variants at the *BIN1* locus. Top, chromatin-accessibility profiles from brain microglia, induced pluripotent stem cell-derived microglia (iPSC-MG), THP-1 macrophages, layer 2/3 excitatory neurons, somatostatin-positive inhibitory neurons, brain astrocytes, and brain oligodendrocytes. Shaded regions mark the positions of the indicated variants. Bottom left, MPRA median absolute deviation (MAD) scores for major- and minor-allele constructs in macrophage and brain contexts. Bottom middle and right, allele-specific, base-resolution importance-score logos generated using the resting macrophage and dorsolateral prefrontal cortex (DLPFC) neuron models, respectively. Pink shading marks the variant position, and dashed yellow boxes highlight selected transcription factor (TF) motif contexts. **(c)** Chromatin accessibility, MPRA allelic activity, and model-based sequence interpretation for selected variants at the *MS4A6E* locus. Chromatin-accessibility tracks, MPRA MAD scores, and allele-specific importance-score logos are displayed as in **b**. Dashed yellow boxes highlight selected ETS-related, E2F, and RFX motif contexts. **(d)** Chromatin accessibility, MPRA allelic activity, and model-based sequence interpretation for selected variants at the *SEC63–OSTM1* locus. Chromatin-accessibility tracks, MPRA MAD scores, and allele-specific importance-score logos are displayed as in **b**. Dashed yellow boxes highlight selected SPI1, AP-1, and ETS-related motif contexts. **(e)** Comparison of gene-expression changes following enhancer and promoter perturbation at the *SEC63–OSTM1* locus. Each point represents a gene. The x-axis shows the log₂ fold change following CRISPR interference (CRISPRi) targeting of the *SEC63* or *OSTM1* promoter, and the y-axis shows the corresponding log₂ fold change following perturbation with the enhancer-targeting sgLeft guide. Columns compare sgLeft perturbation with *SEC63* or *OSTM1* promoter perturbation, and rows show untreated and LPS + IFN-γ-treated THP-1 macrophages. Dashed diagonal lines indicate equal expression changes between perturbations. Spearman correlation coefficients (ρ) are shown; all correlations were statistically significant by a two-sided Spearman correlation test (all P < 1 × 10⁻^53^) **(f)** Gene set enrichment analysis of transcriptional responses to sgLeft enhancer perturbation and *SEC63* or *OSTM1* promoter perturbation in untreated and LPS + IFN-γ-treated THP-1 macrophages. Columns indicate Hallmark gene sets grouped into anabolic, proliferative, and interferon-response programs, and rows indicate perturbation targets under each treatment condition. Color indicates the normalized enrichment score (NES). Asterisks indicate false discovery rate (FDR) < 0.05, and dots indicate FDR < 0.25. Asterisks in the MPRA plots in **b–d** indicate significant allelic differences at FDR < 0.05.

Fine-mapping at *BIN1* has prioritized rs6733839, a SNP within a microglia-specific enhancer near a *SPI1* motif. The minor T allele has been proposed to create an *MEF2* site and is associated with increased chromatin accessibility in iPSC-derived microglia, potentially elevating *BIN1* transcription^3–6,100^. Previous MPRA assays in HEK293T and THP-1 cells showed no significant allele-specific effects, although subtle upregulation was noted^30,31^.

We evaluated rs6733839 using complementary reporter assays and context-specific sequence models (Fig. 6a,b; Extended Data Fig. 9). SNP-centered MPRA constructs again showed no significant allele-specific effects in THP-1 macrophages, brain tissue or HEK293T cells (Extended Data Fig. 9; Supplementary Table 6). In contrast, peak-centered constructs with rs6733839 near the 5′ or 3′ ends showed lower reporter transcription from the T allele in THP-1 macrophages and HMC3 cells (Fig. 6a,b; Extended Data Fig. 9). Machine-learning models predicted increased chromatin accessibility for the T allele in both resting THP-1 macrophage and DLPFC neuron contexts. Feature-attribution profiles implicated a nearby *SPI1*-like sequence feature in the macrophage model and an *MEF2*-like feature in the neuron model, providing candidate sequence-level interpretations of the predicted accessibility effects. Synthetic enhancer tiling and motif-shuffling analyses showed that the regulatory contribution of the rs6733839-centered motif context differed across overlapping sequence tiles (Extended Data Fig. 9). The intact motif context increased reporter activity when paired predominantly with upstream sequence features, retained a partial activating contribution when combined with both upstream and downstream features, and contributed net repressive activity when paired predominantly with downstream features. Thus, the rs6733839-centered sequence context did not have a fixed regulatory effect but instead depended on the surrounding enhancer sequence. We also characterized rs13025717, a variant in the adjacent *BIN1* enhancer whose T allele was previously predicted to disrupt an SP/KLF-like motif and indirectly alter nearby PU.1 (*SPI1*) binding^100,101^ (Fig. 6b). In the THP-1 MPRA, the T allele reduced reporter transcription, and models predicted decreased chromatin accessibility, with feature-attribution profiles implicating disruption of the SP/KLF-like sequence feature. This variant has also been reported as a *BIN1* eQTL in monocytes and a *SPI1* binding QTL in the B-lymphoblastoid cell line GM12878^101^. Another variant, rs13025765, showed opposing effects between THP-1 macrophages and brain tissue, with model-attribution profiles implicating *SPIB*-like features in the immune context and *RFX*-like features in the neuronal context (Fig. 6b). Together, these results nominate context-dependent sequence features that may underlie the divergent reporter effects of selected variants at the *BIN1* locus.

At the *MS4A* locus, a CRE within intron 1 of *MS4A6E* showed strong accessibility in microglial and myeloid contexts, weaker but detectable signals in human hippocampal and DLPFC NeuN+ samples, and a called peak in striatal medium spiny neurons, while its accessibility was reduced following LPS+IFN-γ stimulation in THP-1 macrophages (Fig. 6c and Supplementary Fig. 12a,b). Variants within this CRE exhibited context-dependent allelic effects on reporter transcription accompanied by distinct model-attributed sequence features. The T allele of rs10897049 increased reporter transcription in THP-1 macrophages but showed no significant allelic effect in brain tissue, and model-attribution profiles suggested strengthening of an ETS-like sequence feature. The minor A allele of rs4477457 increased reporter transcription in THP-1 macrophages but decreased it in brain tissue, with neuronal model-attribution profiles implicating disruption of an E2F-like sequence feature. At rs10792265, the minor G allele increased reporter transcription in THP-1 macrophages but showed a non­significant effect in brain tissue. Neuronal model-attribution profiles suggested an allele-dependent shift from an E2F-like sequence feature for the A allele to an RFX-like sequence feature for the G allele. Together, these variants show that allelic changes within the same CRE can produce divergent reporter effects across immune and neural assay contexts.

In the *SEC63–OSTM1* intergenic region, four rare variants within a shared enhancer exhibited distinct allelic effects in the THP-1 macrophage MPRA: the minor alleles of three variants reduced reporter transcription, whereas that of the fourth increased it (Fig. 6a,d). These variants were not in linkage disequilibrium, supporting their interpretation as separate candidate regulatory variants rather than a single linked signal. Model-derived sequence attributions provided interpretable motif changes for three variants: rs147038704-T increased the predicted contribution of an SPI1-related motif despite reducing reporter activity, whereas rs115607757-G weakened a AP-1 site and was associated with reduced reporter activity in macrophages but increased activity in brain tissue. The rs189835276-A allele disrupted an ETS-related motif and increased reporter activity in macrophages (Fig. 6d).

Chromatin analyses using publicly available Hi-C, ATAC-seq and H3K27ac ChIP–seq datasets from THP-1 macrophages^97^ placed this CRE within an accessible, H3K27ac-marked contact domain encompassing the *SEC63–OSTM1* region. Local chromatin contacts connected the CRE to the *OSTM1* promoter and a distal CTCF site near *AFG1L*, while the broader contact domain also encompassed the *SEC63* promoter (Extended Data Fig. 10a,c). Single-nucleus chromatin co-accessibility analysis further predicted an enhancer–*SEC63* promoter connection in microglia (Extended Data Fig. 10b). Together, these analyses nominated both *SEC63* and *OSTM1* as candidate *cis*-linked genes, with the microglial co-accessibility signal preferentially connecting the CRE to the *SEC63* promoter. Both genes were expressed in THP-1 macrophages (Supplementary Table 12).

To investigate the endogenous function of this variant-containing CRE, we performed CRISPR interference (CRISPRi) in THP-1 macrophages under untreated and LPS+IFN-γ-stimulated conditions using three independent sgRNAs tiling the CRE: sgLeft, sgCenter and sgRight, which targeted the left ATAC-seq subpeak, the central accessibility valley and the right subpeak, respectively (Extended Data Fig. 10c). We also targeted the promoters of *SEC63* and *OSTM1*. Promoter-targeting CRISPRi reduced *SEC63* and *OSTM1* mRNA levels in a target-specific manner, whereas CRE-targeting CRISPRi did not produce a consistent steady-state reduction of either transcript across the three guides (Extended Data Fig. 10d,e).

Despite the absence of a reproducible effect on steady-state *SEC63* or *OSTM1* expression, CRE repression produced genome-wide transcriptional changes that resembled those induced by *SEC63* promoter CRISPRi. The genome-wide log₂ fold-change profile induced by sgLeft was strongly correlated with that induced by *SEC63* promoter CRISPRi (Spearman’s ρ = 0.65 following LPS+IFN-γ stimulation and ρ = 0.61 under untreated conditions) but showed little correlation with *OSTM1* promoter CRISPRi (ρ = 0.16 and −0.13, respectively; Fig. 6e). These correlations exceeded the concordance between *SEC63* and *OSTM1* promoter perturbations themselves (ρ = 0.35 and 0.02, respectively). Concordance with *SEC63* perturbation was detected for all three CRE-targeting guides but followed a graded order (sgLeft > sgCenter > sgRight), with non-overlapping bootstrap confidence intervals in both conditions. Following LPS+IFN-γ stimulation, only sgLeft retained a clear preferential correlation with *SEC63* rather than *OSTM1* perturbation (Extended Data Fig. 10f). sgLeft targets the left ATAC-seq subpeak, whose accessibility was selectively reduced following LPS+IFN-γ stimulation, linking the strongest *SEC63*-like transcriptional signature to a stimulus-responsive chromatin subelement (Extended Data Fig. 10c).

At the pathway level, *SEC63* promoter perturbation increased anabolic and biosynthetic gene programs — including mTORC1 signaling, cholesterol homeostasis, glycolysis and protein secretion — with most of these programs showing their strongest enrichment following LPS+IFN-γ stimulation, and reduced Myc- and E2F-associated proliferative programs under untreated conditions. sgLeft CRE repression recapitulated these *SEC63*-associated changes and shared leading-edge genes across the concordant gene sets, whereas *OSTM1* promoter CRISPRi produced a distinct transcriptional profile with limited overlap (Fig. 6f and Supplementary Fig. 13a). CRE repression additionally produced a transcriptional output not shared with *SEC63*: following LPS+IFN-γ stimulation, sgLeft selectively induced interferon-α and interferon-γ response programs (NES = +2.2 and +2.1; FDR < 0.001), whereas *SEC63* promoter perturbation had no significant effect on these programs and *OSTM1* promoter CRISPRi repressed them (NES = −2.8 and −2.5; Supplementary Fig. 13a). This interferon induction was condition-dependent: it was absent under untreated conditions and emerged only after stimulation, a difference we confirmed directly using a guide × stimulation interaction model, in which the interferon-α and interferon-γ response programs were the leading gene sets induced by sgLeft specifically upon stimulation (interaction NES = +2.0 and +1.6; FDR = 0.003 and 0.040; Supplementary Fig. 13b,c). This stimulus-dependent interferon response defines a regulatory output of the CRE that is distinct from its *SEC63*-like metabolic signature. Together, the *SEC63*-like metabolic resemblance and the enhancer-specific interferon response were consistent with the preferential CRE–*SEC63* connection suggested by the locus-level chromatin and co-accessibility analyses, while indicating that the CRE is not a simple transcriptional phenocopy of *SEC63*.

To determine whether these relationships extended to disease-relevant cell states, we projected each perturbation onto marker gene signatures of twelve microglial states and assessed their enrichment by GSEA (Extended Data Fig. 10g). CRE and *SEC63* promoter repression shifted the same microglial-state programs in the same direction — concordantly increasing phagocytic, glycolytic and inflammatory signatures and decreasing the cycling signature under untreated conditions (ρ = 0.76 across co-significant states; 100% sign agreement) — whereas *OSTM1* promoter repression shifted these programs in the opposite direction (ρ = 0.31). Mirroring the interferon result, CRE repression selectively induced the antiviral microglial state (MG11) following LPS+IFN-γ stimulation (NES = +2.2), a response that *SEC63* promoter perturbation did not produce and that *OSTM1* promoter perturbation instead strongly repressed (NES = −3.0). Thus, in the language of microglial states, CRE perturbation reproduced both its *SEC63*-like resemblance and its enhancer-specific, stimulus-dependent antiviral output.

These data link the CRE to an *SEC63*-associated metabolic and biosynthetic transcriptional program, with the strongest concordance arising from perturbation of the stimulus-responsive left subpeak, and identify an additional CRE-specific, stimulation-dependent interferon response. Because CRE repression did not reproducibly alter steady-state *SEC63* expression, the data establish functional convergence with an *SEC63*-associated program but not direct steady-state cis regulation of *SEC63*.

Together, these findings show that variant-containing CREs at the *BIN1*, *MS4A* and *SEC63–OSTM1* loci exhibit sequence- and context-dependent regulatory effects and link the *SEC63–OSTM1* CRE to an endogenous transcriptional program that converges with *SEC63* perturbation.

## Discussion

GWAS and rare-variant sequencing studies continue to expand the catalogue of LOAD risk variants, yet moving from genetic association to molecular mechanism remains challenging. In GWAS, tightly linked variants often share indistinguishable association statistics, making it difficult to distinguish one or more causal or functionally relevant alleles within each LD block. Conversely, WGS nominates rare and ultra-rare variants for which individual-variant association evidence is often limited, requiring complementary functional evidence. These challenges are compounded by the multicellular and state-dependent nature of LOAD: genetic association alone does not reveal which variant is functional, what molecular process it perturbs, or in which cell type or state its consequences emerge. Different classes of variants may act through distinct molecular mechanisms, including effects on protein sequence or RNA processing; for the noncoding variants examined here, the principal mechanisms are expected to involve perturbation of *cis*-regulatory element activity through altered TF binding, local sequence grammar, chromatin accessibility, or enhancer–promoter communication. Functional interpretation must therefore consider both the molecular class of the variant and the disease-relevant cellular context in which its effects may emerge.

Here, we functionally evaluated 599 LOAD-associated noncoding candidate variants by integrating context-resolved *in vitro* and *in vivo* MPRA, interpretable sequence-to-function models, and CRISPRi perturbation. MPRA measured sequence-intrinsic enhancer and allelic activity across immune and neural contexts, the sequence models identified regulatory features associated with these effects, and CRISPRi tested the endogenous consequences of a prioritized element in native chromatin. By examining both common and rare variants across THP-1 macrophage states, HMC3 cells as a complementary CNS-derived cell-line context, and neuron-enriched brain tissue, we identified variants whose allelic effects differed across assay contexts, together with a more limited set of stimulation-dependent effects. These findings show that regulatory effects measured by MPRA can depend on the cellular environment in which a sequence is assayed, although differences in species, delivery and measurement variability limit quantitative comparisons across the experimental systems. Together, this framework connects noncoding LOAD-associated variation to enhancer activity, local regulatory grammar, and endogenous transcriptional responses across disease-relevant contexts.

Previous genetic and epigenomic studies have localized common LOAD risk preferentially to microglial and macrophage regulatory elements^3–6,29,42,43,102^. The WGS-derived rare variants^32^ were included based on the complementary hypothesis that variants nominated near genes involved in neurodevelopment and synaptic function may preferentially affect regulatory activity in neural contexts. Within the assayed candidate set, the relative allelic effect sizes of common and rare variants differed in direction between the two contexts, with common variants showing larger effects in THP-1 macrophages and rare variants larger effects in brain tissue (frequency × context interaction *P* = 0.015). The direction of this contrast was preserved at every MAF threshold used to define the common group, so it does not depend on a particular cutoff. A recent context-resolved modeling study by Marderstein et al. provides a complementary perspective, showing that common noncoding variants tend to have more cell-type-restricted predicted effects, whereas ultra-rare variants have larger and more broadly distributed effects, with particularly strong selective constraint on fetal-neuronal regulatory effects^103^. Their AD-focused analysis further showed that common AD-associated variants were preferentially predicted to exert regulatory effects in microglial contexts. Our experiments extend this framework by directly measuring LOAD-associated variant effects in myeloid and neuron-enriched adult brain environments.

A concern for any such comparison is that common and rare candidate variants were nominated by different studies using different prioritization strategies, so that an apparent frequency effect could instead reflect how each set was assembled. Two features of our library allow this to be addressed rather than only acknowledged. First, the region-based WGS analysis contributed both common and rare variants through a single test applied within nominated genomic regions, and 42 of its 64 loci contain both classes; restricting the comparison to this analysis, and further to variants sharing the same locus, reproduced the interaction and strengthened it (*P* = 0.006 and *P* = 0.009). Second, the two frequency classes overlapped immune open-chromatin peaks at similar rates, and the contrast persisted within each chromatin-accessibility category, indicating that it does not follow the chromatin context from which a variant was nominated. Within the AD-associated regions sampled by this library, allele-frequency class was associated with the assay context in which variants showed larger reporter effects, independently of several measured features of study origin and CRE annotation. We nonetheless regard this as a property of the AD-associated regions sampled by this library rather than of common and rare noncoding variation genome-wide, since every variant assayed here was selected because it lies in a region already implicated in AD. WGS-derived effects oriented by the sign of the locus-level W statistic also showed an aggregate bias shift toward lower reporter transcription in brain tissue; because W is a locus-level statistic, this analysis provides directional evidence at the variant-set level and does not assign risk direction to individual alleles.

Integrating MPRA with sequence-model interpretation revealed LOAD-associated regions containing multiple variants with nonuniform regulatory effects. Such regions may represent distributed regulatory architectures rather than loci that can necessarily be reduced to a single functional lead variant. Variants assigned to the same locus differed in effect magnitude, direction and context dependence and may perturb distinct sequence features within one CRE, affect neighboring CREs or operate preferentially in different cellular environments. The presence of multiple MPRA-defined emVars does not establish that every tested allele is causal, but it indicates that functional prioritization should not necessarily terminate after identification of one plausible candidate.

The motif-context experiments further suggest that regulatory variants cannot always be interpreted as simple disruptions of isolated TF motifs. Shuffling the local sequence surrounding selected variants generally produced larger and often bidirectional transcriptional effects compared with single-nucleotide substitutions, indicating that these variants are embedded within combinations of activating and repressive sequence features. A variant may therefore destroy or create a core TF site, fine-tune motif affinity, or alter the balance among neighboring regulatory inputs. Importantly, the regulatory direction of a TF-associated sequence feature is not fixed: the same TF can exert activating, repressive or context-dependent effects depending on cellular state, cofactor availability, motif arrangement and the surrounding sequence grammar.

The accessibility-trained sequence-to-function models complemented the MPRA measurements by preferentially prioritizing experimentally supported variants when the model-training and assay contexts were matched. THP-1-trained models most effectively prioritized THP-1 MPRA hits, whereas human neuronal models preferentially prioritized brain-tissue hits. Accordingly, the models were most informative for variant ranking and mechanistic interpretation rather than quantitative prediction of MPRA effect size, consistent with the distinct regulatory outputs measured by endogenous chromatin accessibility and reporter transcription. In addition, increased motif affinity, predicted TF occupancy or accessibility does not necessarily imply increased transcription, because the associated TF may act as an activator or repressor depending on its regulatory context. Model-derived motif features should consequently be viewed as mechanistic hypotheses rather than definitive assignments of TF binding or regulatory direction.

Detailed analyses of representative loci illustrated the importance of local sequence context and reporter design for interpreting regulatory effects (Fig. 6; Extended Data Fig. 8). At *BIN1*, allelic effects depended on the sequence window and construct architecture represented in the reporter assay, while motif tiling showed that the contribution of the rs6733839-centered sequence changed with its surrounding enhancer context. These findings indicate that individual motifs should not be interpreted in isolation, because their net contribution can be modified by neighboring activating and repressive features. The imperfect correspondence between accessibility-model predictions and reporter transcription at this locus may therefore reflect both the distinction between the two regulatory readouts and the context-dependent activating or repressive functions of the implicated TF-associated sequence features. Together with the results of Ernst et al.^45^ and Klein et al.^104^ these observations highlight how variant positioning, CRE length and local sequence grammar influence the detection and interpretation of regulatory effects. More broadly, findings across *BIN1*, *MS4A* and other loci indicate that a single reporter window may capture only part of a variant’s regulatory behavior.

Previous MPRA work by Cooper et al. screened dementia-associated variants in HEK293T cells, a system selected partly for its reported epigenomic similarities to neuronal cells^30^. Allelic effects in our HEK293T MPRA were significantly correlated with those reported by Cooper et al., whereas no significant correlation was observed with our *in vivo* brain MPRA, suggesting that this concordance primarily reflects the shared HEK293T assay context and that neural-context effects require direct evaluation in neural tissue. In parallel, Bond et al. examined a larger set of Alzheimer’s disease-associated variants in resting and pro-inflammatory THP-1 macrophages and identified stimulation-dependent allelic effects^31^. Our study complements these studies by testing the same candidate library across multiple myeloid states and a neuron-enriched *in vivo* brain environment, while additionally incorporating WGS-derived rare variants and perturbations of local motif context.

In parallel with the recent work of Bond et al.^31^, we examined variant effects under immune stimulation. LPS+IFN-γ produced broad chromatin, transcriptional and CRE-level changes, but allele-by-stimulation interactions were confined to a limited subset of variants, with allelic directions remaining largely stable across THP-1 activation states. We therefore treat LPS+IFN-γ as a controlled inflammatory perturbation rather than a direct model of LOAD pathology. More broadly, the results show that cellular state can reveal or modify selected regulatory effects that may not be apparent under a single baseline condition.

At the *SEC63*–*OSTM1* intergenic enhancer, we identified four rare variants with distinct allele-specific effects on reporter transcription in THP-1 macrophages. Chromatin-contact and co-accessibility analyses nominated both *SEC63* and *OSTM1* as candidate cis-linked genes, with microglial co-accessibility preferentially connecting the CRE to *SEC63*. However, CRISPRi-mediated repression of the CRE did not consistently alter steady-state expression of either nearby gene. Instead, CRE perturbation produced genome-wide transcriptional and pathway-level responses that more closely resembled *SEC63* promoter repression than *OSTM1* promoter repression, with the strongest concordance observed for sgLeft, which targeted the LPS+IFN-γ-responsive accessibility subpeak. Beyond this *SEC63*-like resemblance, CRE repression—but not *SEC63* promoter repression—engaged an interferon response that emerged only under LPS+IFN-γ stimulation, pointing to a condition-dependent regulatory output of the enhancer that is distinct from, rather than a copy of, the *SEC63*-associated program.

The absence of a reproducible steady-state change in *SEC63* expression leaves open whether this relationship reflects transient or buffered cis regulation or indirect convergence on a shared downstream state. Time-resolved perturbation and endogenous allele editing will distinguish these possibilities and determine how the rare variants contribute to this regulatory program.

These mechanistic analyses were enabled by a shared experimental and computational framework. Assaying the same MPRA library across distinct cellular environments allowed context comparisons without changing candidate selection or construct composition. Optimized electroporation enabled reproducible delivery of the library into THP-1 macrophages, while pseudo-barcode aggregation improved enhancer-level estimates in the lower-coverage *in vivo* datasets. Independently trained accessibility models provided an orthogonal means of prioritizing variants and nominating candidate sequence features. Together, these approaches establish a scalable framework for context-resolved functional characterization of selected noncoding LOAD variants, while retaining complementary experimental readouts of sequence activity, motif context and endogenous function.

This study has four principal limitations. First, the focused candidate library does not comprehensively represent the current LOAD genetic landscape, and the functional assays prioritize regulatory candidates rather than establish genetic causality. The limited sample size of the family-based WGS study leaves uncertainty around individual ultra-rare variant associations, some of which may represent false positives. Second, THP-1 macrophages and HMC3 cells do not fully reproduce primary human microglial identity. Third, the neuron-enriched brain MPRA provides a bulk, cross-species readout that does not model aging or AD pathology. Finally, MPRA and CRISPRi assay complementary aspects of regulatory function but do not reproduce endogenous allele-specific effects. These limitations motivate cell-type-resolved assays and endogenous editing experiments to test the prioritized variants.

Our findings open key avenues for future research, including systematic enhancer–gene mapping, investigation of variant effects in motif–motif interactions, functional studies of prioritized LOAD-associated loci, and defining the cellular states where variants become active. Expanded libraries guided by updated association and fine-mapping resources will enable more complete evaluation of the evolving LOAD genetic landscape and better-matched comparisons of common and rare variants. Single-cell^51,52^ and spatially resolved^54^ reporter assays may resolve effects restricted to specific neural or glial populations, while endogenous base editing, prime editing or allele replacement will be important for determining whether MPRA-prioritized variants reproduce their effects in native chromatin. Together, these efforts will deepen our understanding of noncoding genetic risks and their cellular consequences in LOAD.

Together, our results show that the functional consequences of LOAD-associated noncoding variants depend on both the local sequence in which an allele is embedded and the cellular environment in which that sequence is read. By combining a shared MPRA library with context-specific sequence models and endogenous perturbation, this study establishes a framework for prioritizing candidate regulatory variants, interpreting their local regulatory grammar and testing prioritized elements in native chromatin.

## Methods

### Cell Culture and Animals

#### THP-1 Macrophage Differentiation

THP-1 monocytes (ATCC TIB-202) were cultured in RPMI-1640 supplemented with 10% fetal bovine serum (FBS) at 37°C in 5% CO₂, with a density of under 1 million cells/mL. Differentiation into macrophages was induced by treating cells with 50 nM phorbol 12-myristate 13-acetate (PMA, Sigma-Aldrich) for 72 h, followed by a 24 h resting period in PMA-free medium.

#### HMC3 Cell Culture

HMC3 cells (ATCC CRL-3304) were cultured using Microglial Cell Medium Kit (Accegen, ABI-TM009) and maintained at 37°C with 5% CO₂. Cells were passaged before reaching confluence and seeded at appropriate densities for downstream assays.

#### HEK293T Cell Culture

HEK293T cells (AAVpro(R), Clontech, Cat. No. 632273) were cultured in Dulbecco’s Modified Eagle Medium (DMEM) supplemented with 10% fetal bovine serum (FBS) and maintained at 37°C with 5% CO₂. Cells were passaged into fresh medium before reaching confluence and seeded at appropriate densities for downstream assays.

#### Human iPSC-derived Excitatory Neuron Cell Culture

Human iPSC-derived excitatory neurons (iCell Induced Excitatory Neurons, FUJIFILM Cellular Dynamics, Cat. No. 01279) were cultured on tissue-culture plates sequentially coated with 0.01% poly-L-ornithine and growth-factor-reduced Matrigel. Plates were coated with poly-L-ornithine overnight at 4 °C, washed three times with DPBS without calcium or magnesium, and subsequently coated with Matrigel at a final concentration of approximately 0.083 mg mL⁻^1^ for 1 h at 37 °C. Coated wells were not allowed to dry before cell plating.

Neurons were maintained at 37 °C with 5% CO₂ in complete BrainPhys medium containing iCell Neural Supplement B, iCell Nervous System Supplement, N-2 Supplement and laminin. iCell Plating Supplement B was included during initial cell plating. Cells were thawed according to the manufacturer’s recommendations, recovered by gradual addition of plating medium and centrifuged at 300 × *g* for 3 min at room temperature before plating. Medium was added or replaced gently along the side of each well to minimize neuronal detachment. Cell attachment, soma morphology, neurite integrity and cellular debris were monitored throughout the experiment.

#### Stimulation of THP-1 Macrophages and HMC3 cells

To model diverse immune states, THP-1 macrophages were stimulated for 4 h and HMC3 cells for 24 h with interferon-β (IFN-β, 200 U/mL; PBL Assay Science, Cat. No. 11415-1), interferon-γ (IFN-γ, 10 ng/mL; R&D Systems, Cat. No. 10067-IF-025), or lipopolysaccharide (LPS, 50 ng/mL; Sigma-Aldrich, Cat. No. L2630-25MG) plus IFN-γ (10 ng/mL) to induce antiviral, proinflammatory, and hyperinflammatory states, respectively. Cytokines and LPS were added as small-volume (<20 µL) supplements in DPBS to culture media. Untreated THP-1 macrophages or HMC3 cells received the same culture media and handling. THP-1 signaling responses were assessed by quantification of STAT1 and STAT3 phosphorylation by immunofluorescence imaging using the CellInsight CX5 High Content Screening Platform and HCS Studio software (Thermo Fisher Scientific), as previously described in Cheemalavagu et al^105^.

Additionally, THP-1 macrophages stimulated for 24 h and 72 h were collected for the ATAC-seq characterization of THP-1 macrophages to assess longer-term transcriptional and chromatin accessibility changes.

#### Animals

All procedures were approved by the Carnegie Mellon University Institutional Animal Care and Use Committee (IACUC). Experiments were conducted on 10-week-old C57BL/6J mice (Jackson Laboratory, strain 000664; n = 4, 2 males and 2 females). Animals were housed under specific-pathogen-free conditions in individually ventilated cages (≤ 5 mice per cage) containing corncob bedding and nestlet enrichment. The animal room was maintained at 20-22 °C with 45 % relative humidity and a 12 h:12 h light-dark cycle. Autoclaved standard chow (LabDiet 5001) and UV-irradiated water were provided *ad libitum*. Detailed surgical procedures are described in the section ‘Plasmid Library Transduction of Mouse Brain Regions via AAV’.

### ATAC-seq

#### Cell Preparation and Lysis

THP-1 monocytes were centrifuged and pelleted, lysed using pre-chilled Buenrostro lysis buffer^106^, homogenized with a Dounce homogenizer on ice, and then debris was removed by filtration through a 70 µm mesh. Similarly, THP-1-derived macrophages, adhered to 6-well plates, were lysed with pre-chilled Buenrostro lysis buffer on ice for 10 minutes. Nuclei were dislodged using cell scrapers, filtered through a 70 µm mesh, and pelleted by centrifugation at 500 x g for 5 minutes at 4°C.

#### Nuclei Isolation and DNA Tagmentation

Isolated nuclei were resuspended in nuclease-free water. DAPI-stained nuclei were quantified to approximately 50,000 using a Countess II cell counter (Thermo Fisher Scientific). These nuclei underwent DNA tagmentation using the Illumina Tagment DNA TDE1 Enzyme and Buffer Kits, incubated at 37°C with shaking at 300 rpm for 30 minutes.

#### Library Preparation and Amplification

Tagmented DNA was purified using the Qiagen MinElute PCR Purification Kit. The sequencing library was then prepared using NEBNext High-Fidelity PCR Master Mix and primers (Eurofins Genomics) following the Illumina Nextera tagmentation format as previously published^107^, with the optimal number of amplification cycles determined via qPCR side reactions to be an average of 9 cycles. Final PCR products were purified using the NEB Monarch Nucleic Acid Purification Kit (# T1030).

#### ATAC-seq Library Sequencing

ATAC-seq sequencing libraries were pooled and sequenced on the Illumina NovaSeq™ 6000 platform at GENEWIZ from Azenta Life Sciences using paired-end 150 bp reads. The libraries achieved an average sequencing depth of 68 million reads per sample.

#### ATAC-Seq Data Processing

FASTQ files were processed using the ENCODE ATAC-Seq Pipeline v2.2.3. Paired-end sequencing reads from biological replicates in fastq format were analyzed with the pipeline’s default settings. The Irreproducible Discovery Rate (IDR) optimal peaks, derived from these analyses, were utilized for both differential analyses and machine learning model training.

#### Differential ATAC-Seq Analysis of THP-1 Macrophages upon Stimulation

Common open chromatin regions were identified by merging IDR optimal peaks from both resting and stimulated THP-1 macrophages within a 200 bp proximity. This merging was performed using the bedtools v2.31.1 with the command: bedtools sort -i {input} | bedtools merge -d 200 -i - > {output}. Read counts at each peak for each cell type were quantified using the featureCounts^108^ function from the Rsubread package, utilizing the bam files and bed files of common chromatin peaks. Differential accessibility analysis was executed using the DESeq2^109^ v1.48, with filtering criteria set to include peaks that had at least one read in a minimum of two replicates. Peaks with significantly different accessibility were determined using a false discovery rate (FDR) threshold of <0.05. Differentially accessible promoter regions were annotated using the hg38 human genome assembly, focusing on known genes within a 500 bp range of the identified open chromatin regions.

#### Genetic Variants Enrichment Analysis

To pinpoint cell types enriched for LOAD risk, we applied stratified LD score regression (S-LDSC)^110^ v1.0.1 on open chromatin data from various sources. We obtained open chromatin promoter, enhancer and dyadic peaks for 111 reference epigenomes from the Epigenome Roadmap database^36^. These peaks were merged using bedtools^111^ and treated as background peaks. Following the original S-LDSC methodology, we tested LOAD GWAS heritability enrichment in the open-chromatin peaks of each candidate cell type relative to the Roadmap background. Enrichment p-values were then adjusted across all tests using the Benjamini-Hochberg FDR correction, with enriched cell types defined by a q-value below 0.05. This analysis was performed separately using each GWAS summary-statistics dataset^3–6^. Public open chromatin datasets were used for comparison^28,33,112–119^.

### RNA-seq

#### RNA-seq Library Preparation and Sequencing

Total RNA was isolated with the RNeasy Mini Kit (Qiagen) and RNA integrity was verified on an Agilent 4200 TapeStation. For THP-1 characterization, three biological replicate batches of THP-1 macrophages (± stimulation) and THP-1 monocytes were processed for unstranded poly(A)+ RNA-seq. For CRISPRi samples, total RNA was harvested 4 h after ±stimulation; two biological replicate batches were prepared for (i) poly(A)+ stranded RNA-seq and (ii) rRNA-depleted stranded RNA-seq. All libraries were sequenced by GENEWIZ from Azenta Life Sciences (paired-end 2 × 150 bp).

#### Poly(A)+ Libraries

Total RNA (500 ng per sample) was quantified on a Qubit 2.0 Fluorometer (Thermo Fisher Scientific) and integrity assessed on a 4200 TapeStation (Agilent). External RNA Controls Consortium spike-ins (ERCC Mix 1, Thermo Fisher Scientific, cat. 4456740) were added to the normalized RNA before library construction.

##### Strand-specific libraries

Polyadenylated RNA was enriched and strand-specific libraries were prepared with the NEBNext Ultra II Directional RNA Library Prep Kit for Illumina (New England Biolabs) according to the manufacturer’s guidelines. mRNA was fragmented for 8 min at 94 °C, first-strand cDNA was synthesized, and dUTP was incorporated during second-strand synthesis. After end repair/A-tailing, indexed adapters were ligated and libraries were amplified by limited-cycle PCR.

##### Unstranded libraries

RNA-seq libraries lacking strand information were generated with the NEBNext Ultra II RNA Library Prep Kit for Illumina (New England Biolabs). Poly(A)+ RNA was captured with Oligo(dT) beads and fragmented for 15 min at 94 °C, followed by first- and second-strand cDNA synthesis. Subsequent end repair/A-tailing, adapter ligation, and limited-cycle PCR amplification mirrored the strand-specific workflow.

Library size profiles were confirmed on the TapeStation, and concentrations were determined by Qubit and KAPA Library Quantification qPCR. Equimolar pools were sequenced on an Illumina NovaSeq 6000 in paired-end mode (2 × 150 bp), targeting ∼30 million read pairs per sample.

#### rRNA-depleted Libraries

Total RNA (500 ng) was treated with TURBO DNase (Thermo Fisher Scientific) and rRNA removed using the QIAseq FastSelect-rRNA HMR Kit (Qiagen). Strand-specific libraries were prepared with the same NEBNext Ultra II kit. rRNA-depleted RNA was fragmented for 8 min at 94 °C, followed by first- and dUTP-labelled second-strand synthesis, 3′ adenylation, adapter ligation and limited-cycle PCR (second-strand amplification suppressed by dUTP). Library size was verified on the TapeStation and quantified by Qubit 4.0 and KAPA qPCR. Equimolar pools were clustered on an Illumina NovaSeq X and sequenced in paired-end mode (2 × 150 bp).

#### RNA-seq Data Processing

Raw paired-end reads were trimmed of adapter sequences and quality-checked using FastQC^120^ v0.12.0. Reads were then aligned to the human reference genome (Ensembl GRCh38) using STAR^121^ v2.7.10. Gene-level counts were obtained with featureCounts (Subread package v2.12.2) using ‘--largestOverlap -s 0’ parameters for unstranded RNA-seq and using ‘--largestOverlap -s 2’ for stranded RNA-seq. The resulting count matrices were imported into R, filtered to remove lowly expressed genes, and analyzed with DESeq2^109^ v1.48, adjusting for batch effects where applicable. lfcShrink using Approximate posterior estimation for GLM coefficients (apeglm)^122^ v1.30.0 was used to shrink log2 (fold change) for genes with low read counts. Independent Hypothesis Weighting (IHW)^123^ v1.36.0 was used for multiple testing corrections. Gene annotations were retrieved via biomaRt v2.64.0 to map Ensembl gene IDs to standard gene symbols.

#### Differential RNA-seq Analysis of THP-1 Macrophages upon Stimulation

Untreated and IFN-β-, IFN-γ- and LPS + IFN-γ-treated THP-1 macrophages were analyzed jointly using DESeq2 with stimulation condition and experimental batch included in the design (∼Treatment + Batch). The untreated condition was used as the reference, and contrasts were extracted separately for each stimulation condition versus untreated macrophages. Differential-expression estimates were subjected to the log₂-fold-change shrinkage, annotation and multiple-testing procedures described above.

#### Differential Analysis of THP-1 Macrophage Nucleofection-control RNA-seq

Nucleofection-control samples were analyzed using DESeq2 with experimental batch and delivery condition included in the design (∼Batch + condition). The nucleofection-reagent-only group was used as the reference, and nucleofection-only, pMaxGFP and MPRA-backbone conditions were each contrasted with this control. Previous cytokine-treated THP-1 macrophages were additionally contrasted with the unstimulated MPRA-backbone condition. Differential-expression estimates were processed using the log₂-fold-change shrinkage, annotation and multiple-testing procedures described above. Hallmark pathway enrichment was performed with fgsea using genes ranked by the unshrunk DESeq2 Wald statistic.

#### Analysis of RNA-seq from iCell Human iPSC-derived Excitatory Neurons

Whole-transcriptome RNA-seq data from two biological replicates of the human iPSC-derived excitatory neurons described above (iCell Induced Excitatory Neurons, FUJIFILM Cellular Dynamics) were processed using the general RNA-seq alignment and gene-counting workflow. Gene-level counts generated by featureCounts were converted to transcripts per million (TPM) using annotated gene lengths. Ensembl gene-version suffixes were removed before matching gene identifiers across replicates. The neuronal identity of the iCell neurons and the potential expression of markers associated with non-neuronal cell types were assessed using a predefined marker panel. The panel included pan-neuronal markers (*NEFM*, *STMN2*, *TUBB3*, *MAP2*, *DCX* and *ELAVL4*); excitatory-neuronal and synaptic markers (*SLC17A6*, *SLC17A7*, *SNAP25*, *SYT1*, *SYN1*, *DLG4* and *GRIA1*); pluripotency markers (*POU5F1* and *NANOG*); inhibitory-neuronal markers (*GAD1*, *GAD2* and *SLC32A1*); astrocyte markers (*GFAP*, *AQP4*, *SLC1A3* and *ALDH1L1*); oligodendroglial markers (*OLIG2*, *PDGFRA*, *MBP* and *MOG*); and microglial or immune markers (*PTPRC*, *AIF1* and *CX3CR1*). Marker expression was visualized separately for the two biological replicates after transformation as log₂(TPM + 1). The heat map displayed absolute transformed expression values without gene-wise scaling or differential-expression testing.

#### Differential analysis of CRISPRi RNA-seq

Genes were retained if they had at least 10 reads in at least 16 of the 32 samples. Untreated and LPS + IFN-γ-treated samples were analyzed separately using DESeq2. *SEC63*- and *OSTM1*-promoter perturbations were each tested in isolated promoter-versus-control models after excluding the other promoter-targeting and all enhancer-targeting samples. Enhancer-targeting sgRNAs directed to the left, center and right regions were analyzed together with shared control samples after excluding promoter-targeting samples. Models included perturbation identity and library-preparation method (∼sgRNA + Method) when both poly(A)+ and rRNA-depleted libraries were present. Control sgRNA libraries were used as reference levels. Differential expression was estimated separately for each perturbation relative to control within the untreated and LPS + IFN-γ-treated conditions, followed by the log₂-fold-change shrinkage, annotation and multiple-testing procedures described above.

#### Gene Set Enrichment Analysis (GSEA)

Hallmark gene sets were obtained from the msigdbr package v25.1.1 by filtering for “H” category (Hallmark). Genes were ranked based on a statistical measure derived from their differential expression results ‘unshrunk log2FoldChange / lfcSE’. After removing duplicates and low-confidence entries, fgsea^124^ v1.34.2 was performed on the ranked list, examining Hallmark pathways of size 15-500 genes. Significant pathways (adjusted *p* < 0.05) were visualized in bar plots of Normalized Enrichment Scores.

#### Microglia-state Concordance

Marker-gene sets for human microglial states and amyloid-fibril-treated iPSC-derived microglia were obtained from Sun et al.^25^ For each THP-1 stimulation or CRISPRi contrast, genes were ranked by unshrunk log2FoldChange/lfcSE after removal of duplicate and low-confidence entries. Enrichment was tested using fgsea, and signatures with FDR < 0.05 were considered significant.

### MPRA Plasmid Library Design

#### Candidate variant selection and library composition

To prioritize LOAD-associated variants for evaluation with both machine learning models and reporter assays, we first assembled a collection of sentinel LOAD GWAS variants from Jansen et al.^4^ and Kunkle et al.^6^ that met a significance threshold of 5×10⁻^8^. We then broadened this collection by including variants in linkage disequilibrium with the sentinel variants, using HaploReg (v4.1)^125^, which leverages data from the 1000 Genomes Project^126^. Applying an LD r^2^ threshold of 0.5 resulted in an expanded set of 1,209 variants, of which 205 constructs representing 163 variants were carried into the library.

Additional candidate variants were contributed through the Cure Alzheimer’s Fund CIRCUITS consortium from three further analyses. First, cell-type-resolved fine-mapping of AD GWAS loci by peak-to-gene linking and aQTL–GWAS colocalization^29^ contributed 114 constructs representing 67 variants; we used a pre-publication version of these analyses, which at that stage covered microglia and excitatory neurons for the linking analysis and applied a colocalization posterior probability threshold of PP4 ≥ 0.1.

Second, region-based association testing was performed in the family-based NIMH and NIA ADSP whole-genome sequencing cohorts using the spatial-clustering approach described previously^32^, including the same two-pass design in which each region is tested using either all variants regardless of allele frequency or rare variants only, and the same *P* < 5 × 10⁻^4^ threshold. Regions were defined differently here: rather than genome-wide, clustering was restricted to variants overlapping microglial, monocyte and neuronal open-chromatin peaks. Seventy-six regions comprising 379 variants were carried forward, contributing 459 constructs. Third, 66 constructs representing 48 variants were contributed from a further selection of the single-variant association results from the same study^32^.

Variants from the GWAS, LD-expansion and fine-mapping sources were filtered to retain those within 113 bp of an open-chromatin peak summit, allowing each variant to be centred within a 227-bp synthesized enhancer. Variants from the region-based whole-genome sequencing analysis were retained in full, having already been selected within microglial, monocyte and neuronal open-chromatin peaks. Variants from the single-variant analysis were retained without open-chromatin filtering. In total, 844 allele-pair constructs were analysed; individual variants may be represented by more than one construct where peak-centred and SNP-centred designs, distal placements, or multiple alternate alleles were included (Supplementary Table 3). A further 163 constructs carried a 25-bp shuffled window centred on the variant position, designed to disrupt the local transcription-factor motif context; these do not form allele pairs and were analysed separately (Supplementary Table 7).

A further 11 constructs targeted CpG sites nominated by epigenome-wide association analyse^67^s of AD contributed through the CIRCUITS consortium, on the rationale that a CpG showing differential methylation may also fall within a transcription-factor recognition sequence. For each site we substituted the cytosine of the CpG dinucleotide with guanine. These are not naturally occurring variants and carry no allele frequency; they were excluded from all analyses comparing common and rare variants.

#### Annotation of LOAD Genetic Variants

Open-chromatin peak sets from diverse human cell types were assembled from the human brain cATLAS and additional in-house and public bulk and single-cell ATAC-seq datasets^28,29,33,112–119,127–130^. Bulk ATAC-seq narrowPeak files were processed with the ENCODE ATAC-seq pipeline v2.2.3, whereas published single-cell ATAC-seq peak sets were used as provided. Variants in GRCh38/hg38 coordinates were intersected with each peak set using a minimum overlap of 1 bp to construct a binary variant-by-cell-type accessibility matrix.

Cell types were assigned to cell-subclass and cell-class annotations using the accompanying metadata. Within each cell subclass, a variant was classified as accessible if it overlapped an open-chromatin peak in at least one constituent cell type. Cell-subclass accessibility values were then summarized into neuronal, immune and non-immune glial scores. The neuronal score was calculated across GABAergic and glutamatergic neuronal subclasses; the immune score was calculated across central-immune, peripheral-immune and blood-cell subclasses; and the glial score was calculated across non-immune glial subclasses. Cell types assigned to the Other class contributed to the assessment of broad accessibility but were not used to define an Other-enriched category.

Variants were assigned to five predefined accessibility categories using fixed rule-based thresholds. Variants accessible in no more than one cell subclass were classified as Low-ATAC. Variants were classified as Broad when at least two cell classes had an accessibility fraction of at least 0.10 and the highest-scoring class accounted for no more than 0.65 of the summed class score. Otherwise, a variant was classified as Immune, Neuronal or Glial when the corresponding class had a score of at least 0.16 and exceeded the second-highest class by either a ratio of at least 1.5 or an absolute difference of at least 0.08. Remaining variants accessible across multiple classes were classified as Broad, whereas the remaining variants were assigned to their highest-scoring class. The resulting categories were termed Immune, Neuronal, Glial, Broad and Low-ATAC.

For visualization, pairwise Jaccard distances were calculated from the binary accessibility matrix, and average-linkage hierarchical clustering was performed within predefined cell-class and variant-category blocks. Clustering was used only to order rows and columns in the heat map and did not determine variant-category assignment. The R package annotatr^131^ v1.34.0 was used to annotate variants with genomic features.

#### Design of *Cis*-regulatory Elements (CREs)

A total of 599 unique common and rare variants from LOAD GWAS and WGS were incorporated into 227 base pair synthetic CREs. Variants overlapping open chromatin peak summits in microglia and monocytes, which represent regions most likely to serve as TF binding sites, were prioritized. When a variant overlapped a peak in immune cells or neurons, the summit served as the center of the enhancer. For variants not overlapping open chromatin peaks, SNP-centered enhancers were designed. Additional constructs were generated for variants positioned distal (83 bp) to peak summits and for those with multiple alternate alleles. In addition, 25 base pair motif-shuffled versions of the CREs were produced as positive controls to disrupt TF binding, particularly for variants predicted to affect immune- and neuron-related factors in haploR^56,132^ (MEF2, SPI1, IRF, STAT, NF-κB, AP-1).

#### MPRA Experiments Plasmid Library Assembly

A pool of 24,000 unique 300-nt oligonucleotides (Twist Bioscience, San Francisco, CA, USA) was PCR-amplified (12 cycles; 66 °C annealing) with KAPA HiFi HotStart Polymerase (Roche) using primer (5′-GGCTACCGTCTCACATGGAGTCTGAACCTGTGTGCTA-3′) and primer (5′-GGCTACCGTCTCGACCACTACGATGCTCTTCCGATCT-3′). Amplicons were verified on a D1000 ScreenTape system (Agilent) and purified with the Monarch PCR & DNA Cleanup Kit (NEB).

The MPRA backbone (pAAV-Hsp68-nls/mCherry-MPRAe) was linearized by PCR (35 cycles; 72 °C annealing) with Q5 High-Fidelity DNA Polymerase (NEB) using primer (5′-GGCTACCGTCTCCCATGTTTCTCCATTTGGCGC-3′) and primer (5′-GGCTACCGTCTCGTGGTACTTAGCATCTCAC-3′). The product was gel-separated and extracted with the Monarch Gel Extraction Kit (NEB).

Purified insert and the linearized vector were assembled at a 2:1 molar ratio using the NEB Golden Gate Assembly Kit (BsmBI-v2) under the following thermal program: 42 °C for 1 h, 60 °C for 5 min, then 4 °C hold. 4 µL of the assembly reaction was electroporated into 100 µL of ElectroMAX 10-beta T1 E. coli (Thermo Fisher Scientific). Transformants were expanded in 3 L Terrific Broth at 37 °C, and plasmid DNA was purified with the QIAGEN Plasmid Plus Giga Kit.

#### Plasmid Library Transfection of Stimulated THP-1 Macrophages

On the day of transfection, the culture supernatant was collected for later use. The cells were detached from the dish with Invitrogen Accumax at 37 °C for 30 min. Transfection was performed using the Nucleofection P3 Primary Cell 4D-Nucleofector kit (Lonza Bioscience) with the DP-148 program. In each electroporation, 2 µg of the MPRA plasmid library was used to transfect 3 million THP-1 macrophages. Cells from five electroporations were pooled and seeded in 6-well plates, then incubated at 37 °C for 1 hour to allow attachment. Next, the medium containing the nucleofection reagent was replaced with the previously collected supernatant, and interferons were added. 4 hours post-electroporation, cells were lysed directly in the wells, and lysates were snap-frozen and stored at -80 °C for RNA extraction. A replicate of each condition was scraped from the plates, snap-frozen in a separate tube, and stored at -80 °C for genomic DNA extraction.

Untreated MPRA samples underwent nucleofection with 2 µg of the MPRA plasmid library per 3 × 10^6^ cells and the same post-nucleofection handling as stimulated samples but received no cytokine or LPS treatment. Thus, the untreated MPRA condition represents a delivery-exposed, unstimulated baseline rather than a non-nucleofected resting state.

#### Plasmid Library Transfection of THP-1 Monocytes

THP-1 monocytes in suspension were pelleted by centrifugation at 300 × g for 5 min. Transfection was conducted using a LONZA Nucleofector 4D device with the SG Cell Line 4D-Nucleofector kit with the FF-100 program. In each electroporation, 1.5 µg of the MPRA plasmid library was used to transfect 3 million THP-1 monocytes. 4 hours post-electroporation, the cells were pelleted, washed with DPBS, and lysed. The resulting lysates were snap-frozen and stored at -80 °C. In parallel, a portion of each cell pellet was snap-frozen in a separate tube for genomic DNA extraction.

#### Plasmid Library Transfection of HMC3 cells and HEK293T Cells

3 million HMC3 or HEK293T cells were seeded in 100 mm dishes 24 hours before transfection. For each dish, 18 µg of the MPRA plasmid library was transfected using 54 µL of Promega FuGene 6. At 24 hours post-transfection, HEK293T cells were washed with DPBS and lysed in the dish for total RNA extraction, and HMC3 cells were treated with interferons. 24 hours after interferon treatment, HMC3 cells were washed with DPBS and lysed in the dish for total RNA extraction. Immediately before cell lysis, 25% of each sample was scraped from the plate, snap-frozen in a tube, and stored at - 80 °C for genomic DNA extraction.

#### Plasmid Library Transduction of Human iPSC-derived Excitatory Neurons via AAV

Following optimization of AAV delivery conditions, approximately six million human iPSC-derived excitatory neurons across four independent biological replicates were transduced with the MPRA library packaged in AAV-PHP.eB at a multiplicity of infection of 2.5 × 10^6^ vector genomes per cell. Parallel cultures were transduced with an AAV-PHP.eB single-stranded GFP reporter at the same or comparable multiplicity of infection to confirm that the neurons were permissive to AAV-PHP.eB transduction. Untransduced cultures were maintained as controls.

Following transduction, cultures were monitored for cell attachment, soma morphology, neurite integrity, reporter fluorescence, cellular debris and overt toxicity. RNA and DNA were collected six days after transduction. Total RNA and genomic DNA were processed using the same MPRA cDNA, genomic-DNA and sequencing-library preparation procedures described above.

#### Plasmid Library Transduction of Mouse Brain Regions via AAV

The plasmid pool was packaged into adeno-associated virus PHP.eB (AAV-PHP.eB) following Brown et al.^49^. Adult C57BL/6J mice (n = 4; two males, two females; 10 weeks old) were anesthetized with 3-4 % isoflurane in oxygen until respiration slowed and the pedal-withdrawal reflex was lost. Each mouse was maintained in 1-2% isoflurane and received two retro-orbital injections of 2 × 10^12 vector genomes (vg) in 85 µl phosphate-buffered saline (PBS), administered 48 h apart. Immediately after each injection, 0.5 % proparacaine hydrochloride ophthalmic solution was applied to the treated eye for comfort, and the animals were monitored continuously for signs of pain or distress. weeks after the second injection, mice were deeply re-anesthetized with isoflurane to the point of absent reflexes and euthanized by decapitation in compliance with institutional guidelines. The cortex, hippocampus and striatum were rapidly micro­dissected from 300 µm coronal brain sections on an ice-chilled stage and snap-frozen on a −80 °C cooling block for subsequent extraction of total RNA and genomic DNA.

#### cDNA MPRA Sequencing Library Preparation

cDNA libraries were prepared according to the sysMPRA^49^ protocol. Cell pellets or frozen tissues were used to extract total RNA following the standard protocol of the Qiagen RNeasy kit, including on-column DNase digestion. The resulting RNA samples were further treated with Invitrogen TURBO DNase to remove residual genomic DNA, followed by a cleanup step using the Qiagen MinElute kit. First-strand cDNA was synthesized using Invitrogen SuperScript IV reverse transcriptase and a customized plasmid-sequence-specific primer. MPRA sequencing libraries were then prepared by PCR amplification (26 cycles) with NEBNext High-Fidelity 2X PCR Master Mix and Nextera primers. The PCR products were purified with the NEB Monarch Nucleic Acid Purification kit, quantified using Qubit, and assessed for purity on a TapeStation.

#### Genomic DNA MPRA Sequencing Library Preparation

Genomic DNA was extracted from cell pellets or frozen tissues using the Qiagen DNeasy Blood & Tissue Kit. MPRA sequencing libraries were then generated by PCR amplification (26 cycles) with NEBNext High-Fidelity 2X PCR Master Mix and the same plasmid-sequence-specific primers from Brown et al.^49^. For cell-derived samples, 20 ng of genomic DNA was used as the template; for brain tissue, 200 ng of genomic DNA was used. The PCR products were purified using the NEB Monarch Nucleic Acid Purification kit, quantified by Qubit, and evaluated on a TapeStation for product purity.

#### MPRA Library Sequencing

MPRA sequencing libraries were pooled with ATAC-seq libraries to increase library diversity and sequenced on the Illumina NovaSeq™ 6000 platform at GENEWIZ from Azenta Life Sciences using paired-end 150 bp reads. The cDNA libraries achieved an average sequencing depth of 16 million reads per sample, while the genomic DNA (gDNA) libraries received an average of 4 million reads per sample.

#### MPRA Biological Replicates

All biological replicates were generated from independent experimental batches. For cultured-cell MPRAs, cells were cultured, transfected or nucleofected, treated where applicable and harvested independently on different days. The final datasets comprised eight untreated, five IFN-β-treated, five IFN-γ-treated and five LPS+IFN-γ-treated THP-1 macrophage replicates; five untreated, five IFN-β-treated, four IFN-γ-treated and four LPS+IFN-γ-treated HMC3 replicates; and three HEK293T replicates. The brain-tissue MPRA comprised cortex, hippocampus and striatum samples from four independent mice. The pilot human iPSC-derived excitatory-neuron MPRA comprised four biological replicates but was not included in primary enhancer-level or allele-specific analyses because of extensive RNA barcode dropout.

#### THP-1 Macrophage Nucleofection Controls

To evaluate transcriptional responses induced by nucleofection and plasmid-DNA delivery, differentiated THP-1 macrophages were subjected to four parallel conditions: a nucleofection-reagent control without electrical pulsing or plasmid DNA, nucleofection without plasmid DNA, nucleofection with the pMaxGFP control plasmid, and nucleofection with the MPRA backbone plasmid. For each condition, 3 × 10^6^ THP-1 macrophages were processed using the P3 Primary Cell 4D-Nucleofector Kit and subjected to the same downstream handling used for the THP-1 macrophage MPRA experiments. Electrical pulsing was performed using the DP-148 program. Plasmid-containing conditions received 2 µg of either pMaxGFP or MPRA backbone plasmid DNA. Cells in the nucleofection-reagent control were exposed to the nucleofection solution and handled in parallel but were not subjected to an electrical pulse and did not receive plasmid DNA.

Following nucleofection, cells were seeded in 6-well plates. The nucleofection medium was then replaced with the previously collected culture supernatant, as described for the MPRA experiments. No cytokines or LPS were added to the delivery-control conditions. Cells were harvested 4 h after nucleofection, and total RNA was extracted and processed for RNA-seq using the procedures described below. Each condition included two independent biological replicates generated in separate experimental batches. These samples were used to assess delivery-associated transcriptional responses and were not included in MPRA enhancer- or allele-specific quantification.

### MPRA Data Analysis

#### Read Mapping and Preprocessing

The paired-end sequencing reads of the cDNA and DNA prepared from each sample were mapped to generate a read count matrix according to both 10 bp sequences at 5’ end and 3’ end of the enhancer and the 8 bp barcodes associated with each enhancer. 21 enhancers that have barcode read counts underrepresented in the DNA read count matrix were filtered out in the analysis. The read count matrix was transformed into an enhancer table and allele table. The enhancer table contains one unique enhancer per row and individual barcode counts for each sample in separate columns. The allele table has each ALT-allele:REF-allele comparison or MOTIFDISRUPT:REF allele comparison in each row and barcodes unique to each allele in a specific sample in each column.

#### Pseudo-barcodes for Sparse Data

Read-count matrices from mouse brain tissue and THP-1 monocytes were sparse and showed frequent dropouts in both RNA and DNA, introducing noise in quantification. To mitigate this, read counts of similarly represented barcodes were aggregated into pseudo-barcodes.

DNA counts were used to evaluate the method because barcode distributions are expected and found to remain constant across experiments and cell types, whereas RNA counts vary with enhancer activity. Thus, increased DNA-count correlation between replicates was interpreted as reduced technical noise.

For each enhancer, 15 barcodes were ranked by total read counts within brain tissue or THP-1 monocytes; every three adjacent barcodes were summed to generate five pseudo-barcodes per enhancer. This transformation improved DNA read-count correlation (Spearman ρ ≈ 0.75) between biological replicates and between sparse and dense datasets.

Based on this validation, pseudo-barcodes were applied to RNA read counts of brain tissue and THP-1 monocytes, while original barcodes were retained for other cell types. Sparse DNA read counts in these tissues were further stabilized by substituting raw tissue DNA counts with pooled, normalized DNA counts derived from high-depth non-tissue DNA libraries from the same plasmid pool. Pseudo-barcodes were then generated from these substitute DNA counts to match the RNA pseudo-barcodes. For non-tissue samples, observed DNA counts were used directly.

#### Quantification of Enhancer Activities

We used the quantitative analysis function of MPRAnalyze^87^ v1.26.0 to estimate transcriptional activity for each candidate CRE from paired cDNA and DNA counts. Each LOAD-associated candidate CRE was represented by 15 preassigned barcodes. For each assay context, the regression design was specified as dnaDesign = ∼Barcode + Replicate and rnaDesign = ∼1. Enhancer activity was summarized using the one-sided median absolute deviation (MAD)-based activity score and corresponding p.MAD.Score returned by MPRAnalyze. Candidate CREs with p.MAD.Score < 0.10 were classified as active. For visualization of selected examples, transcriptional activity was estimated separately for each biological replicate.

The library included nine scrambled negative-control sequences previously used in Brown et al.^49^, with each scrambled sequence represented by 31–32 barcodes. These sequences served as empirical references but were not used as the basis for formal enhancer-activity classification.

#### Differential Enhancer Activity Analysis

We tested differential CRE activity using MPRAnalyze^87^ (v1.26.0). For each cell type (THP-1 macrophages; HMC3 cells) and contrast (stimulated vs untreated), we fit a comparative framework with dnaDesign = ∼Barcode + Replicate, rnaDesign = ∼Stimulation and, reducedDesign = ∼1. CREs with FDR < 0.05 were called differential, and direction was assigned by the sign of the logFC (stimulated vs resting).

#### Pathway Enrichment Analysis

We analyzed differential CREs with Genomic Regions Enrichment of Annotations Tool (GREAT)^133^ v4.0.4, running up-regulated and down-regulated sets separately as foregrounds and using the complete MPRA library as background. Genomic coordinates of test variants within these CREs (GRCh38/hg38) were supplied to GREAT using the default setting; enrichment was evaluated using GREAT’s region-based binomial and gene-based hypergeometric tests. Terms were reported at FDR < 0.05.

#### Quantification of Allelic Differences and Motif Disruption

To compare transcriptional differences attributable to allelic variants and motif shuffling, we used the comparative analysis function of MPRAnalyze^87^ v1.26.0 with at least 4 biological replicates per condition. In these analyses, we compared constructs harboring the CRE with the major versus minor allele, as well as constructs in which the motif was shuffled. The regression design was specified as dnaDesign = ∼Barcode + Replicate, rnaDesign = ∼Allele, reducedDesign = ∼1.

To examine the effect of interferon stimulation on allelic differences, we used the design dnaDesign = ∼Barcode + Replicate, rnaDesign = ∼Allele + Stimulation + Allele:Stimulation, reducedDesign = ∼Allele + Stimulation. For aggregated analyses across stimulated THP-1 macrophages, HMC3 cells, and mouse brain regions, we included an additional covariate “Condition” in both the rnaDesign and reducedDesign to account for the different conditions.

Pairwise Spearman correlations were calculated between MPRAnalyze allelic-effect estimates across untreated, IFN-β-, IFN-γ- and LPS + IFN-γ-treated THP-1 macrophages. Variants with allelic FDR < 0.05 were summarized by the number of THP-1 conditions in which they were significant, and effect-direction concordance was determined from the sign of the allelic log fold change. Differences in allelic effects between THP-1 macrophages and brain tissue were tested using an allele-by-context interaction model. Counts were reported at the tested-element level unless otherwise specified; unique-variant counts were obtained by collapsing multiple construct configurations representing the same variant.

#### Comparison of common and rare variant effect sizes across assay contexts

Variants were classified as rare (EUR MAF < 0.01) or common (EUR MAF ≥ 0.01) using the frequencies in Supplementary Table 3. Effect size was the absolute allelic log fold change from MPRAnalyze. Because the two assays differ in overall effect-size scale, distributions were compared within context using two-sided Kolmogorov–Smirnov tests and Cliff’s δ, a rank-based directional effect size independent of sample size; confidence intervals for δ were obtained by bootstrap resampling of constructs (6,000 resamples for the MAF-threshold analysis, 10,000 for the category analysis).

Covariate-adjusted analyses used ordinary least squares with outcome log₁₀(|allelic logFC| + 0.001) and the following covariates: mean reporter activity of the reference and alternate constructs (MPRAnalyze MAD score; reference constructs matched to alternate constructs using the library design lookup table), the five-level ATAC-based variant category (Broad as reference), construct design (peak-centred versus SNP-centred), GC content, and log₁₀ of the absolute distance to the nearest transcription start site. Continuous covariates were z-scored. Models were fitted separately within each context (n = 844 constructs assayed in both contexts).

The frequency × context interaction was estimated in a pooled model in which every covariate was interacted with context, which is algebraically equivalent to the two per-context fits; the interaction coefficient therefore equals the difference between the two context-specific rare-variant coefficients. Standard errors were clustered on construct because each construct contributes one observation per context. Because residuals departed from normality, the interaction was additionally assessed by a construct-level bootstrap (5,000 resamples) and by permutation of the rare-variant label (5,000 permutations). Analyses used statsmodels v0.14 and scipy.

#### Motif-based Prediction of Enhancer Activity

MPRA sequences were scanned with FIMO v5.5.5 (MEME-Suite; http://meme-suite.org) against the HOCOMOCO v12 CORE motif library using a threshold of *P* < 1 × 10^−4. For each enhancer, hits sharing the same HOCOMOCO accession (for example, MA0139) were merged by averaging log-likelihood scores, yielding a motif-by-enhancer score matrix.

Condition-specific enhancer activities (MAD Scores) from THP-1 macrophages (Resting, LPS+IFN-γ) and mouse brain tissue were joined to the motif matrix on enhancer ID; absent motifs were set to zero.

To identify motifs associated with activity, we fitted an Elastic Net model to each condition. Motif scores were z-standardized, and ElasticNetCV from scikit-learn v1.5.2 (https://scikit-learn.org) was run with the default l₁ ratio of 0.5 and the package’s built-in grid of 100 logarithmically spaced α values (five-fold cross-validation, max_iter = 10^5^, random_state = 0). After selecting the best hyper-parameters, we refitted the model once and estimated the significance of the in-sample R^2^ by permuting the response vector 1,000 times; the empirical P-value equals the fraction of permuted R^2^ values greater than or equal to the observed R^2^. Model fit was reported as R^2^ and its permutation-based P. Non-zero coefficients were retained for downstream interpretation.

#### Motif Enrichment Analysis

For each cell type we extracted 25-bp sequences centred on MPRA-validated expression-modulating variants (emVars). Motif enrichment was assessed with AME v5.5.6 (MEME-Suite) using the HOCOMOCO v12 core mouse–human TF motif collection. Enrichment was calculated with the Wilcoxon rank-sum test (--method ranksum) and average log-odds scoring (--scoring avg); only the top 25 % of motif hits in the target set were considered (--hit-lo-fraction 0.25). The THP-1 emVar sequences served as the control set for brain emVars (and vice-versa) via the --control option. Motifs with an E-value < 10 and Benjamini-Hochberg–adjusted q < 0.05 were deemed significantly enriched.

#### Analysis of the Human iPSC-derived Excitatory Neuron MPRA

Enhancer activity was estimated using the same within-assay MPRAnalyze quantitative framework and one-sided MAD-based activity statistic used for the primary MPRA experiments. Activity of nine scrambled negative-control sequences was compared with that of seven mouse cortical and three human cortical positive-control sequences from Nguyen et al.^90. Statistical comparisons were performed using one-sided Mann–Whitney U tests followed by Benjamini–Hochberg correction. Because RNA barcode recovery showed extensive dropout, the iPSC-neuron dataset was used only to assess activity of the cortical positive-control sequences and was excluded from the primary enhancer-level and allele-specific variant analyses.

#### Association-aligned MPRA effects

To compare genetically associated alleles with MPRA allelic effects, we analyzed common variants reported by Bellenguez et al.^5^ and rare variants included in the Prokopenko et al.^32^ whole-genome sequencing burden analysis. Bellenguez variants were matched to MPRA variants by rsID and exact allele pair. The AD risk allele was assigned from the reported effect estimate for the minor allele, with β > 0 indicating the minor allele and β < 0 indicating the major allele. Prokopenko variants were matched by hg38 genomic coordinate and exact allele pair to variants within each burden region. As the regional W statistic was oriented to the minor allele, W > 0 and W < 0 were used to assign the minor and major allele, respectively, as the association-aligned allele.

Analyses were restricted to SNP-centered or peak-centered constructs with significant allelic effects in brain tissue or THP-1 macrophages (FDR < 0.05). MPRA effect sizes were reoriented to the genetically associated allele, such that positive values indicated reduced reporter transcription and negative values indicated increased reporter transcription. For variants represented by multiple significant constructs, the effect direction was assigned by majority vote; variants with equal numbers of constructs in each direction were classified as ties. Variant-level effects were summarized by genetic locus and assay context.

### Machine Learning Model Training and Prediction

#### Data Preprocessing for Machine Learning

For each cell type, IDR optimal peaks (NarrowPeak format) were resized to a uniform length of 500 bp centered on the peak summit. The model input consisted only of the underlying DNA sequence of each resized peak. As the supervised target, we used the ATAC-seq peak intensity recorded in column 7 of the NarrowPeak file (signalValue). To enable supervised learning and calibration of background sequence preferences, we generated mononucleotide-distribution–matched negative sequences using BiasAway v3.3.0 (biasaway g --foreground {foreground} --nfold 1 --bgdirectory {background} --seed 19951124). Each positive peak sequence was paired with one GC-matched negative sequence, and negative sequences were required to have no overlap with positive peak intervals. Negative examples were assigned a target signal of 0. To minimize information leakage across genomic regions, sequences were split by chromosome into training, validation, and test sets: chromosome 4 was used for validation, chromosomes 8 and 9 were held out for testing, and all remaining chromosomes were used for training.

#### Convolutional Neural Network Training

Convolutional neural networks (CNNs) were trained separately for each cell type or state (one model per NarrowPeak file) using only ATAC-seq peak DNA sequences as input and the corresponding peak accessibility signalValue as the regression target. Each input DNA sequence is one-hot encoded as a 4 × 500 matrix, and used to train a CNN implemented in TensorFlow v2.7 to predict signalValue from sequence alone, thereby learning motif-like sequence patterns and motif-combination features associated with chromatin accessibility. Peak category annotations (e.g., immune, ubiquitous, low-ATAC) were not provided to the model and were not used in dataset construction, loss weighting, or model selection; these labels were assigned only for downstream stratified evaluation of model outputs. The architecture comprised sequential layers: Input, Conv1D, Dropout, Conv1D, Dropout, MaxPooling1D, Flatten, Dense, Dropout, Dense, Dropout, Dense, and Output. Hyperparameters are listed in Supplementary Table 13.

#### Predicting Chromatin Accessibility Using Machine Learning Models

Candidate sequences, derived from open chromatin peaks, were standardized to 500 bp centered on the peak summit. Peaks from differential analyses were either expanded or trimmed to 500 bp, centered on their middle point. Both forward and reverse complementary sequences of each input were one-hot encoded into 4×500 matrices. These matrices were then fed into the trained convolutional neural network. The model computed the predicted signal value for each input by averaging the peak signals of the forward and reverse complementary sequences.

#### Model Interpretation

To interpret model predictions at base-pair resolution, we employed SHapley Additive exPlanations (SHAP)^92^ v0.46.1 on both validation datasets and genomic sequences with genetic variants. The SHAP values were input in TF-MODISCO-Lite^93^ v1.0.0 and match TF motifs in the HOCOMOCO V12 Core motif database. The enriched motifs with q value < 0.05 were reported.

#### Allele-specific Differences in Predicted ATAC Accessibility and SHAP Contribution Scores

For each candidate variant, we extracted paired 500-bp genomic sequences centered on the variant position, differing solely at the major- or minor-allele nucleotide. Each allele-specific sequence was evaluated using the trained model in both forward and reverse-complement orientations. Strand-specific predictions were averaged to yield a single ATAC-seq accessibility score per allele, and the allelic effect was quantified as the difference between major- and minor-allele scores.

For each allele, we calculated the SHAP contribution score of each nucleotide across the 500-bp region and derived an allelic SHAP contrast by computing the major-minus-minor difference.

To establish a baseline null distribution, we selected GC-matched control sequences from non-peak genomic regions within a held-out validation set, ensuring the central nucleotide matched the major allele of the query candidate variant. We then mutated the central nucleotide to the minor allele nucleotide of the query variant. Differences in predicted accessibility and SHAP scores between these control sequence pairs defined the null distributions, reflecting expected variability in non-functional regions. All differences underwent log10 transformation. The transformed null distributions displayed substantial overlap, whereas candidate SNPs exhibited a marked rightward shift suggestive of functional relevance. Null distributions were modeled using a normal distribution, enabling computation of variant-specific P values. Variants exhibiting P values < 0.1 were prioritized and visualized using a heat map.

#### Estimating Prediction Uncertainty

To quantify predictive uncertainty, we adopted Monte-Carlo dropout. Specifically, we re-enabled all Dropout layers in the trained CNN and drew 64 stochastic forward passes per input, using 0.3 dropout rate to approximate sampling from the model’s posterior. The resulting ensemble of outputs was summarized by the mean prediction as well as its standard deviation, skewness, and kurtosis, providing a principled estimate of both aleatoric and epistemic uncertainty. To quantify the statistical confidence of allele-specific differences in predicted ATAC accessibility, we performed two-sided Wilcoxon signed-rank tests on the 64 Monte-Carlo dropout replicates per allele and controlled the false-discovery rate with Benjamini–Hochberg correction.

#### MPRA-hit Enrichment among Model-prioritized Variants

For each ATAC-trained sequence model, allele-specific predicted accessibility effects were calculated from paired major- and minor-allele 500-bp SNP-centered sequences and compared with matched negative-SNP predictions to obtain one-sided P values from a fitted log-normal null distribution. For each model group, the smallest P value across constituent models was used as the group-level variant score, and variants were ranked by −log10(P). MPRA hits were defined as variants with allelic-effect FDR < 0.05. We calculated the fraction of MPRA hits among variants ranked in the top 5–49% of model scores in 1% increments.

#### CRISPR Interference Plasmid Cloning

We constructed a dCas9-KRAB-MeCP2 plasmid by fusing the bipartite dCas9 repressor (Addgene: 122205) with a third-generation lentiviral backbone (Addgene: 83889). Fragments were PCR-amplified using NEB Q5 High-Fidelity DNA Polymerase and assembled with the NEBuilder HiFi DNA Assembly kit. sgRNA oligonucleotides, synthesized by Eurofins Genomics, were annealed via a thermal cycler and ligated into the direct-capture Perturb-seq sgRNA backbone (Addgene: pBA904), pre-digested with Thermo FastDigest Restriction Enzymes (BstXI and Bpu1102I), using NEB T4 DNA Ligase. Assembled plasmids were amplified in NEB 5-alpha competent E. coli and purified with the QIAprep Spin Mini Plasmid Prep kit.

#### Guide RNA Design

sgRNA spacers, designed to initiate with “G” to optimize mouse U6 promoter activity, targeted the promoters of *SEC63, OSTM1,* and the enhancer of interest using CRISPick (Broad Institute). Two safe-targeting and one non-targeting sgRNAs were selected from published sources^134,135^. All sgRNA sequences are provided in Supplementary Table 14.

#### Lentivirus Production

HIV-based lentiviral particles were produced by transfecting 0.75 × 10^6^ HEK293T cells in 2.5 ml DMEM with 3 µg total plasmid DNA comprising pMDLg/pRRE (Addgene plasmid #12251), pRSV-Rev (Addgene plasmid #12253), pMD2.G (Addgene plasmid #12259) and the lentiviral transfer plasmid at a mass ratio of 2:1:1:4 using Lipofectamine 3000. Cells were cultured at 37°C for 2-4 days, after which virus-containing supernatants were filtered through a 0.45 µm PVDF filter (Millipore Sigma) and either used immediately or stored at −80°C.

#### Stable Cell Generation

THP-1 dCas9-KRAB-MeCP2 line. THP-1 monocytes (3 × 10^5 cells per well, 6-well plate) were resuspended in 1 mL RPMI-1640 (10% FBS) containing polybrene (10 µg/mL); 1 mL dCas9-KRAB-MeCP2 lentivirus (prepared in DMEM, 10% FBS; polybrene maintained at 10 µg/mL) was added (total 2 mL per well). Cells were incubated 72 h at 37 °C; medium was replaced and cells were incubated 24 h; on day 5, blasticidin (15 µg/mL) was applied for 3 days. Medium was then replaced with antibiotic-free medium; stable pools were expanded in flasks with periodic maintenance selection.

sgRNA transduction. THP-1 dCas9-KRAB-MeCP2 cells (3 × 10^5 per well, 6-well plate) were resuspended in 1 mL RPMI-1640 (10% FBS) with polybrene (10 µg/mL); 1 mL sgRNA lentivirus (pBA904 backbone) was added (total 2 mL per well). After 72 h, medium was replaced and cells were incubated 24 h; on day 5, cells were selected for 3 days with puromycin (2.5 µg/mL) and blasticidin (15 µg/mL). Medium was then replaced with antibiotic-free medium; cells were allowed to recover ≥24 h before experiments.

#### Sample Collection for RNA-seq

Stable THP-1 cell lines were differentiated into macrophages via 3 days of PMA treatment followed by 1 day of resting. On day 4, macrophages were treated with or without LPS+IFN-γ for 4 hours, lysed in-plate, and total RNA was extracted for RNA-seq as described previously.

#### Genome-Wide Transcriptional Concordance and Pathway Enrichment Analysis

Differential-expression estimates for each CRISPRi perturbation were matched by Ensembl gene identifier. Genome-wide concordance between each CRE-targeting sgRNA (sgLeft, sgCenter and sgRight) and *SEC63*- or *OSTM1*-promoter CRISPRi was assessed across genes with a mean normalized count (baseMean) >= 50 and finite unshrunken log2 fold-change estimates in both comparisons using two-sided Spearman rank correlations. Concordance between *SEC63*- and *OSTM1*-promoter CRISPRi was also calculated as a reference. Untreated and LPS+IFN-γ-stimulated samples were analyzed separately. For pathway analysis, genes were ranked by the DESeq2 Wald statistic, defined as the unshrunken log2 fold change divided by its standard error, and preranked gene-set enrichment analysis was performed separately for each perturbation and condition against the MSigDB^136^ Hallmark 2020 collection using a weighted enrichment statistic. Gene sets containing 15–500 genes represented in the ranked list were tested using 1,000 gene-set permutations with a fixed random seed. Duplicate gene symbols were represented by the highest-ranking entry. Normalized enrichment scores and FDR q values were used for visualization; FDR < 0.05 and FDR < 0.25 were denoted by an asterisk and a middle dot, respectively.

To formally test whether the sgLeft transcriptional response was condition-dependent, we additionally fit a DESeq2 model for sgLeft that included a guide × stimulation interaction term. Genes were ranked by the interaction Wald statistic — equivalent to the difference between the LPS+IFN-γ and untreated log2 fold changes — and preranked gene-set enrichment analysis was performed against the MSigDB Hallmark 2020 collection^136^ using the parameters described above.

### Enhancer-promoter Mapping

#### Enhancer-promoter Mapping with the Activity-by-Contact Model

THP-1 enhancer-promoter interactions were predicted with the Activity-by-Contact (ABC) algorithm^96^ v1.1.2. All input data (Hi-C, ATAC-seq, and H3K27ac ChIP-seq for resting cells and for cells stimulated 4 h with LPS+IFN-γ) were obtained from the same GEO series, GSE201376 (Reed *et al.*^97^, hg19). The publicly supplied 5-kb Hi-C contact matrix was used as-is, whereas the raw ATAC-seq and ChIP-seq FASTQ files were re­processed with the ENCODE ATAC-seq v2.2.3 and ChIP-seq pipelines v2.2.2 (hg19 alignment, default parameters). ABC scores were computed from the re-processed activity tracks and the Hi-C contacts, and enhancer-promoter pairs with ABC ≥ 0.02 were retained as putative regulatory links.

#### ABC target genes versus nearest genes

emVars overlapping distal, non-promoter ABC cis-regulatory elements (CREs) were assigned to the gene linked by the highest ABC score. Overlapping CREs were merged to generate a nonredundant set. For each CRE, the ABC-linked gene was compared with the nearest protein-coding gene, defined by transcription start site proximity. CRE-to-TSS distances were calculated using hg38 coordinates.

#### Human brain single-nucleus chromatin co-accessibility

Precomputed chromatin co-accessibility tracks were examined using the human brain ATAC-seq browser in cATLAS^130^. The *SEC63–OSTM1* locus was queried using the GRCh38 genome assembly, and the microglial co-accessibility track was inspected to evaluate potential connections between the variant-containing CRE and nearby gene promoters. Co-accessibility links were displayed using the browser’s default settings and were not recomputed or statistically reanalyzed in this study.

## Supplementary Information

**Supplementary Table 1 | ATAC-seq promoter accessibility in THP-1 cells**

Genome-wide differential-accessibility statistics for all promoters (±1 kb from the annotated TSS) profiled in THP-1 monocytes and in macrophages that were unstimulated or stimulated with IFN-β, IFN-γ, or LPS+IFN-γ. For each stimulus-versus-resting comparison, the table lists genomic coordinates, base-mean accessibility, log₂-fold change, Wald statistic, and Benjamini–Hochberg-adjusted *P* value (*padj*).

**Supplementary Table 2 | RNA-seq gene-expression profiles in THP-1 cells**

Gene-level differential-expression statistics for THP-1 monocytes and macrophages exposed to IFN-β, IFN-γ, or LPS+IFN-γ, alongside the unstimulated macrophage state. For every annotated gene, the table reports transcript ID, gene symbol, base-mean expression, log₂-fold change relative to the baseline, Wald statistic, and Benjamini–Hochberg-adjusted *P* value (*padj*).

**Supplementary Table 3 | MPRA oligonucleotide library design**

A complete list of all variants and their corresponding oligonucleotides included in the MPRA designed to test regulatory variants associated with LOAD, with functional annotations for every element.

**Supplementary Table 4 | Enhancer activity in MPRAs**

Enhancer activities for each *cis*-regulatory element (CRE) measured by MPRA, based on 15 unique barcode replicates per CRE.

**Supplementary Table 5 | State-specific enhancer activities in immune-cell MPRAs**

Differential enhancer activities in THP-1 macrophages and HMC3 cells following IFN-β, IFN-γ, or LPS+IFN-γ stimulation, reported as log₂-fold change and false-discovery-rate-adjusted *P* values relative to the resting state.

**Supplementary Table 6 | Allelic MPRA effects (emVars)**

Allelic activity differences (reference versus alternate) for all expression-modulating variants (emVars) identified in the MPRA experiments, including effect size (log₂-fold change) and *padj*. Log₂-fold changes for major-versus-minor alleles need to be adjusted according to the ‘REFALT_Flip’ indicator listed in Supplementary Table 3.

**Supplementary Table 7 | Motif-shuffled effects in MPRAs**

Transcriptional differences of motif-shuffled CREs relative to the corresponding CREs harboring LOAD genetic variants.

**Supplementary Table 8 | State-specific allelic effects in MPRAs**

Allelic activity differences that are specific to immune stimulation in THP-1 macrophages and HMC3 cells, with interaction statistics for allele × stimulus effects.

**Supplementary Table 9 | Predicted variant effects from machine learning**

Allele-specific regulatory impact scores from convolutional neural network (CNN) models and splicing-impact scores from SpliceAI for every variant assayed.

**Supplementary Table 10 | Prioritized functional variants across cell types**

Integrated list of high-confidence functional regulatory variants in THP-1 macrophages, mouse brain tissue, HMC3 cells, and HEK293T cells, combining MPRA and computational predictions.

**Supplementary Table 11 | ABC promoter–enhancer links for candidate variants**

Activity-by-contact (ABC) model results linking candidate variants to putative target promoters in THP-1 macrophages, including ABC scores and genomic coordinates of interacting elements.

**Supplementary Table 12 | CRISPRi RNA-seq in THP-1 macrophages**

Differential gene-expression statistics from CRISPRi perturbations of candidate regulatory elements or their nearest-gene promoters in THP-1 macrophages, reported as log₂-fold change versus non-targeting control and Benjamini–Hochberg-adjusted *P* values.

**Supplementary Table 13 | CNN architecture and training hyper-parameters**

Layer-by-layer specification of the convolutional neural network used to predict ATAC-seq accessibility. Table details input shape, layer type, kernel size, stride, number of filters/units, activation function, dropout rate, optimizer, learning-rate schedule, batch size, and training epochs.

**Supplementary Table 14 | CRISPRi sgRNA designs**

Sequences and genomic targets of all sgRNAs used for enhancer and promoter knock-downs. For each guide RNA we list the target category (enhancer / promoter / non-targeting / safe-targeting), oligo sequences (5′→3′) and spacer sequences.

## Data Availability

Illumina sequencing data were deposited in Gene Expression Omnibus (GEO) with the accession number ***.

## Code Availability

Code used for MPRA analysis is publicly available at https://github.com/pfenninglab/context_ad_mpra

The CNN training, evaluation and variant-prediction pipeline is available at https://github.com/zihengc/cnn-transformer-pipeline

Code used for ATAC-seq, RNA-seq, CRISPRi and associated downstream analyses is available at https://github.com/pfenninglab/context_ad_multiomics/

## Acknowledgements

We thank Spencer Gibson for piloting the Monte Carlo dropout procedure that informed our assessment of model-prediction uncertainty. We are grateful to Jianglin Zhang for validating sysMPRA in gut tissues through molecular imaging that confirmed systemic transduction and assay performance. We acknowledge Xiantong Xin’s efforts in piloting ATAC-seq of THP-1 monocytes.

## Author Contributions

**Z.C.** and **A.R.P.** conceived the study. **Z.C.** performed the experiments, analyzed the data, and wrote the original draft of the manuscript. **Y.L.** assisted with the MPRA and CRISPRi experiments, cloned CRISPR plasmids, and, together with **L.G.**, produced lentiviral vectors. **A.R.B.** provided ATAC-seq and MPRA protocols and packaged the AAV vectors, and **B.N.P.** provided training and protocols for AAV injection and brain microdissection. **H.H.S.** developed and maintained the convolutional neural network training pipelines, and **I.K.** contributed to cross-species model predictions and CNN pipeline development. **S.W.** contributed to neuronal model development. **P.H.** advised on chromatin-interaction interpretation. **R.V.D.W.**, **K.E.S.**, and **R.A.G.** contributed to additional experiments and manuscript revision. **Z.C.**, **E.R.**, **B.N.P.**, **X.X.**, **D.P.**, **L.-H.T.**, **L.B.**, **W.H.**, **R.E.T.**, and **M.K.** contributed fine-mapped genetic-variant datasets. All authors reviewed, edited, and approved the manuscript.

---

**Extended Data Fig. 1.**
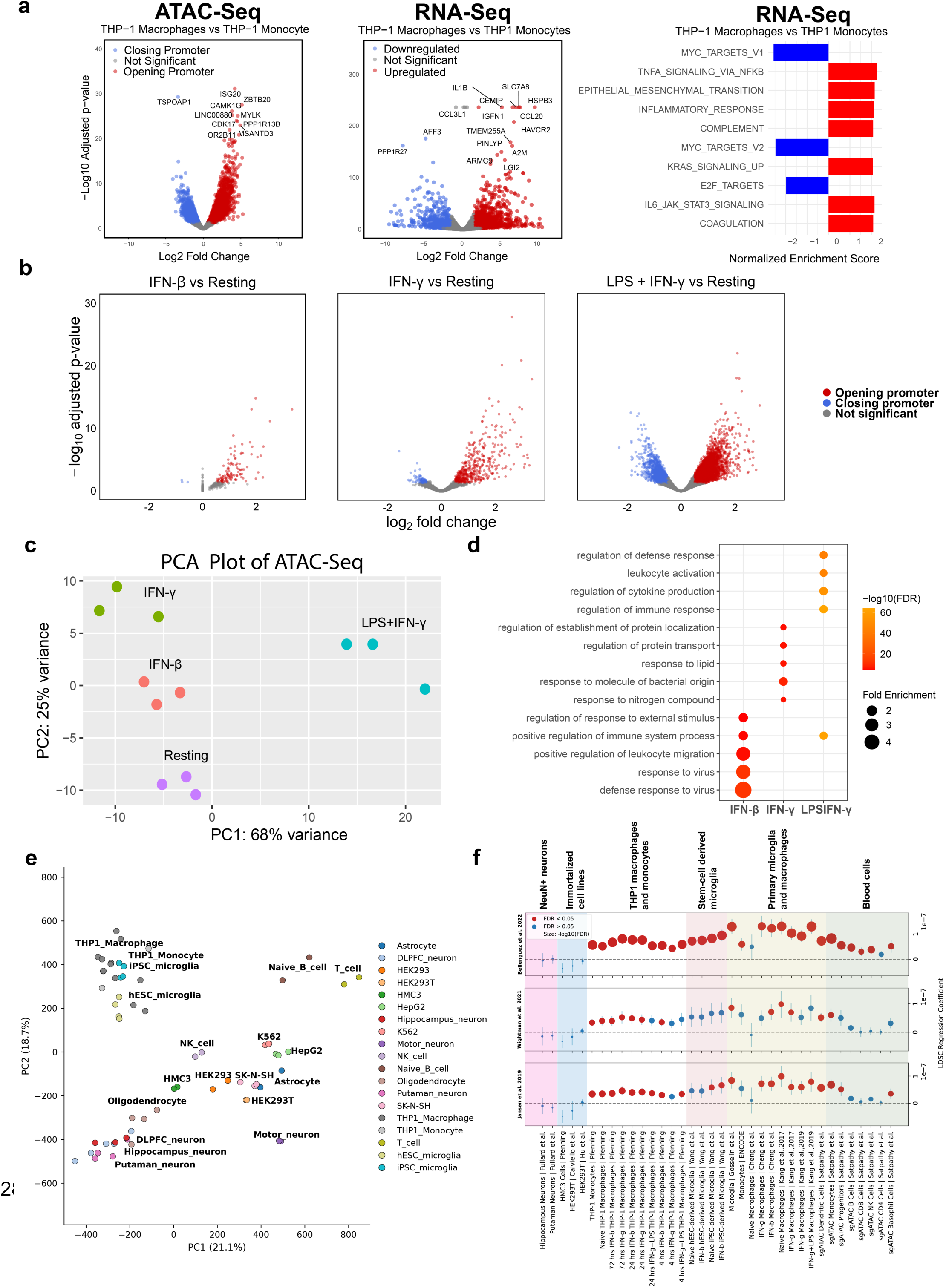
Epigenomic characterization of THP-1 macrophage differentiation and stimulation, with LOAD heritability enrichment. **(a)** Epigenomic and transcriptomic changes associated with THP-1 macrophage differentiation. Left, volcano plot showing differential chromatin accessibility at promoter regions in THP-1 macrophages relative to undifferentiated THP-1 monocytes. Red points indicate promoters with increased accessibility, blue points indicate promoters with decreased accessibility, and gray points indicate nonsignificant promoters. Middle, volcano plot showing differential gene expression in THP-1 macrophages relative to undifferentiated THP-1 monocytes. Red points indicate upregulated genes, blue points indicate downregulated genes, and gray points indicate nonsignificant genes. Labels identify selected promoters or genes. Right, gene set enrichment analysis (GSEA) of RNA-seq changes associated with macrophage differentiation. Bars indicate normalized enrichment scores (NES); red and blue indicate positively and negatively enriched gene sets, respectively. Differentiated THP-1 macrophages exhibited increased inflammatory and cytokine-associated programs, including TNF-α signaling via NF-κB, inflammatory response, complement, IL-6–JAK–STAT3 signaling, and coagulation, together with reduced MYC- and E2F-associated proliferative programs. **(b)** Volcano plots showing differential promoter accessibility following 4 h of stimulation with IFN-β, IFN-γ, or LPS + IFN-γ relative to resting THP-1 macrophages. Red points indicate promoters with increased accessibility, blue points indicate promoters with decreased accessibility, and gray points indicate nonsignificant promoters. **(c)** Principal-component analysis (PCA) of ATAC-seq profiles from resting and stimulated THP-1 macrophages. Axes indicate the percentages of variance explained by principal components 1 and 2. **(d)** Gene ontology (GO) enrichment analysis of genes associated with promoters that gained accessibility following stimulation. Columns indicate stimulation conditions, and rows indicate enriched biological processes. Dot color indicates −log₁₀(FDR), and dot size indicates fold enrichment. **(e)** PCA comparing THP-1 ATAC-seq profiles with publicly available chromatin-accessibility datasets from neuronal, immune, induced pluripotent stem cell-derived, and immortalized cell contexts. Axes indicate the percentages of variance explained by principal components 1 and 2. **(f)** Stratified linkage disequilibrium score regression (S-LDSC) analysis of late-onset Alzheimer’s disease (LOAD) GWAS heritability across open-chromatin annotations from neuronal, immortalized, THP-1, stem-cell-derived microglial, primary microglial/macrophage, and blood-cell datasets. The three panels show LDSC regression coefficients obtained using independent LOAD GWAS datasets. Red points indicate annotations significant at FDR < 0.05, whereas blue points indicate nonsignificant annotations. Point size indicates −log₁₀(FDR).

**Extended Data Fig. 2.**
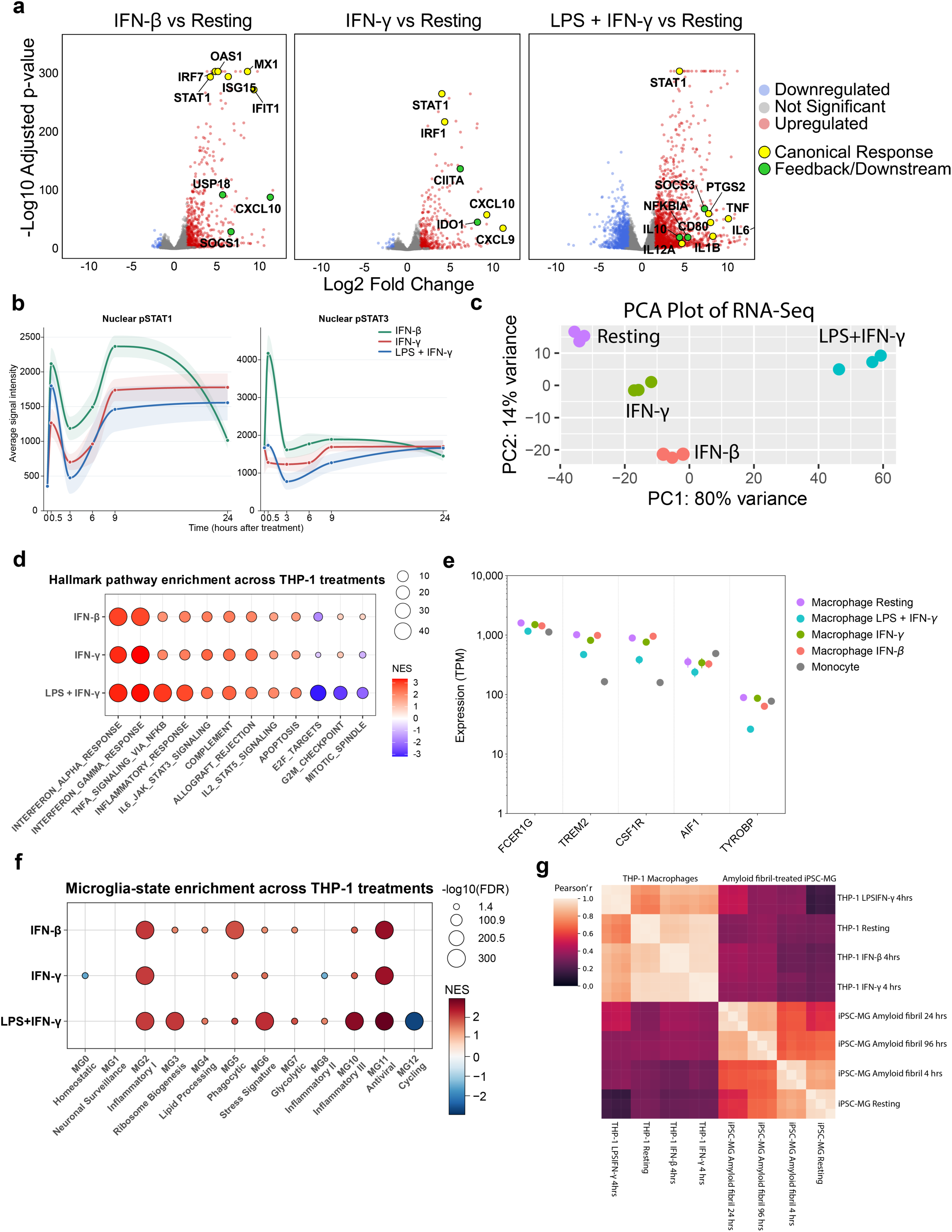
Signaling and transcriptomic characterization of stimulated THP-1 macrophages and their correspondence to human microglial states. **(a)** RNA-seq volcano plots showing differential gene expression in THP-1 macrophages stimulated with IFN-β, IFN-γ, or LPS + IFN-γ relative to resting THP-1 macrophages. The x-axis indicates log₂ fold change, and the y-axis indicates −log₁₀(adjusted *P* value). Red and blue points indicate significantly upregulated and downregulated genes, respectively, and gray points indicate nonsignificant genes. Yellow and green highlighted points indicate selected canonical-response and feedback or downstream-response genes, respectively. **(b)** Nuclear phosphorylated STAT1 (pSTAT1) and STAT3 (pSTAT3) signal intensities measured over time following IFN-β, IFN-γ, or LPS + IFN-γ stimulation. Points indicate the observed mean signal intensities at each time point, curves indicate PCHIP-smoothed temporal trajectories, and shaded bands indicate PCHIP-smoothed nominal 95% *t*-based confidence intervals. For stimulated time points, confidence intervals were calculated from cell-level measurements within each well; the 0-h baseline was calculated across pooled resting well means. **(c)** Principal-component analysis (PCA) of RNA-seq profiles from resting, IFN-β-stimulated, IFN-γ-stimulated, and LPS + IFN-γ-stimulated THP-1 macrophages. Axes indicate the percentages of variance explained by principal components 1 and 2. **(d)** Hallmark gene set enrichment analysis of transcriptional changes following IFN-β, IFN-γ, or LPS + IFN-γ stimulation relative to resting THP-1 macrophages. Dot color indicates the normalized enrichment score (NES), and dot size indicates −log₁₀(false discovery rate [FDR]). **(e)** Expression of selected myeloid and microglia-associated marker genes in THP-1 monocytes and resting or stimulated THP-1 macrophages. The y-axis indicates expression in transcripts per million (TPM) on a logarithmic scale. **(f)** Enrichment of human microglial-state gene signatures in THP-1 macrophages following IFN-β, IFN-γ, or LPS + IFN-γ stimulation relative to resting cells. Columns indicate previously defined microglial states. Dot color indicates NES, and dot size indicates −log₁₀(FDR). **(g)** Pearson correlation heatmap comparing transcriptomic profiles of resting and stimulated THP-1 macrophages with those of resting and amyloid-fibril-treated induced pluripotent stem cell-derived microglia (iPSC-MG). Color indicates the Pearson correlation coefficient (*r*).

**Extended Data Fig. 3.**
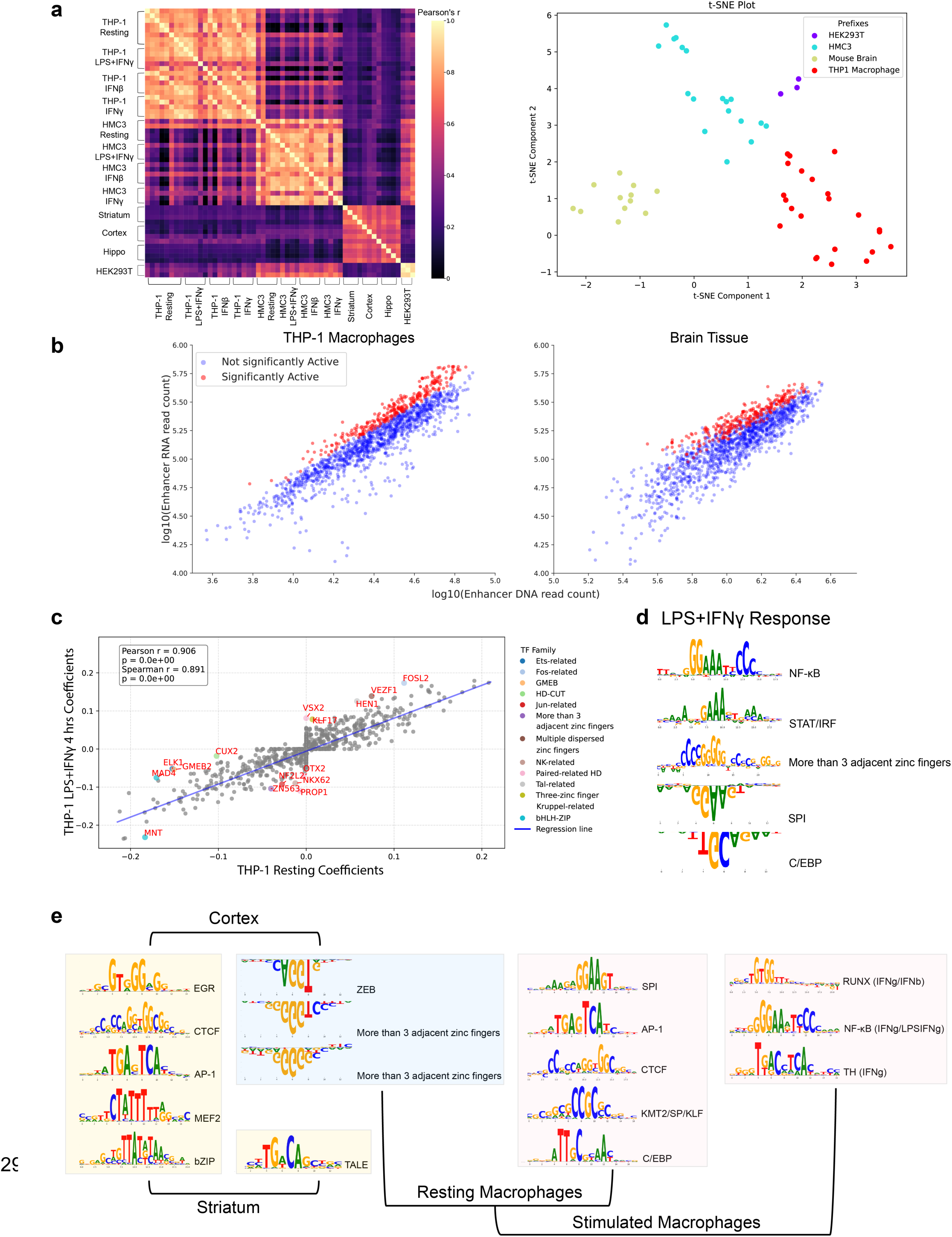
MPRA reproducibility, context-specific enhancer activity, and motif features associated with macrophage and brain regulatory programs. **(a)** Comparison of massively parallel reporter assay (MPRA) enhancer activity across biological replicates and assay contexts. Left, Pearson correlation heatmap of candidate *cis*-regulatory element (CRE) activity across resting and stimulated THP-1 macrophages, resting and stimulated HMC3 cells, mouse brain regions, and HEK293T cells. Color indicates the Pearson correlation coefficient (*r*). Right, t-distributed stochastic neighbor embedding (t-SNE) projection of CRE activity profiles, showing separation of samples according to assay context, including THP-1 macrophages, HMC3 cells, mouse brain tissue, and HEK293T cells. **(b)** Relationship between enhancer DNA abundance and reporter RNA abundance in THP-1 macrophages and mouse brain tissue. Each point represents a candidate CRE. The x- and y-axes show log₁₀-transformed enhancer DNA and RNA read counts, respectively. Red points indicate CREs classified as significantly active by one-sided MPRAnalyze quantitative analysis, and blue points indicate CREs not classified as significantly active. **(c)** Comparison of elastic-net regression coefficients for HOCOMOCO transcription factor (TF) motif features associated with MPRA enhancer activity in resting and LPS + IFN-γ-treated THP-1 macrophages. Each point represents a motif feature. The x-axis shows coefficients estimated in resting THP-1 macrophages, and the y-axis shows coefficients estimated after 4 h of LPS + IFN-γ stimulation. The blue line indicates the linear regression fit. Pearson and Spearman correlation coefficients are shown. Red labels identify selected motif features with large residuals from the regression fit. **(d)** TF-MoDISco motif logos derived from the LPS + IFN-γ-versus-resting differential chromatin-accessibility model. NF-κB- and STAT/IRF-related motifs were associated with increased predicted accessibility following stimulation, whereas SPI1- and C/EBP-related motifs were associated with decreased predicted accessibility. A motif assigned to a cluster containing more than three adjacent zinc-finger motifs is also shown. **(e)** Representative TF-MoDISco motif logos derived from convolutional neural network models trained on context-specific ATAC-seq profiles. Motifs are shown for cortex, striatum, resting macrophage, and stimulated macrophage models. Brackets indicate the model contexts in which each motif group was identified, highlighting shared and context-associated sequence features across neural and macrophage regulatory programs.

**Extended Data Fig. 4.**
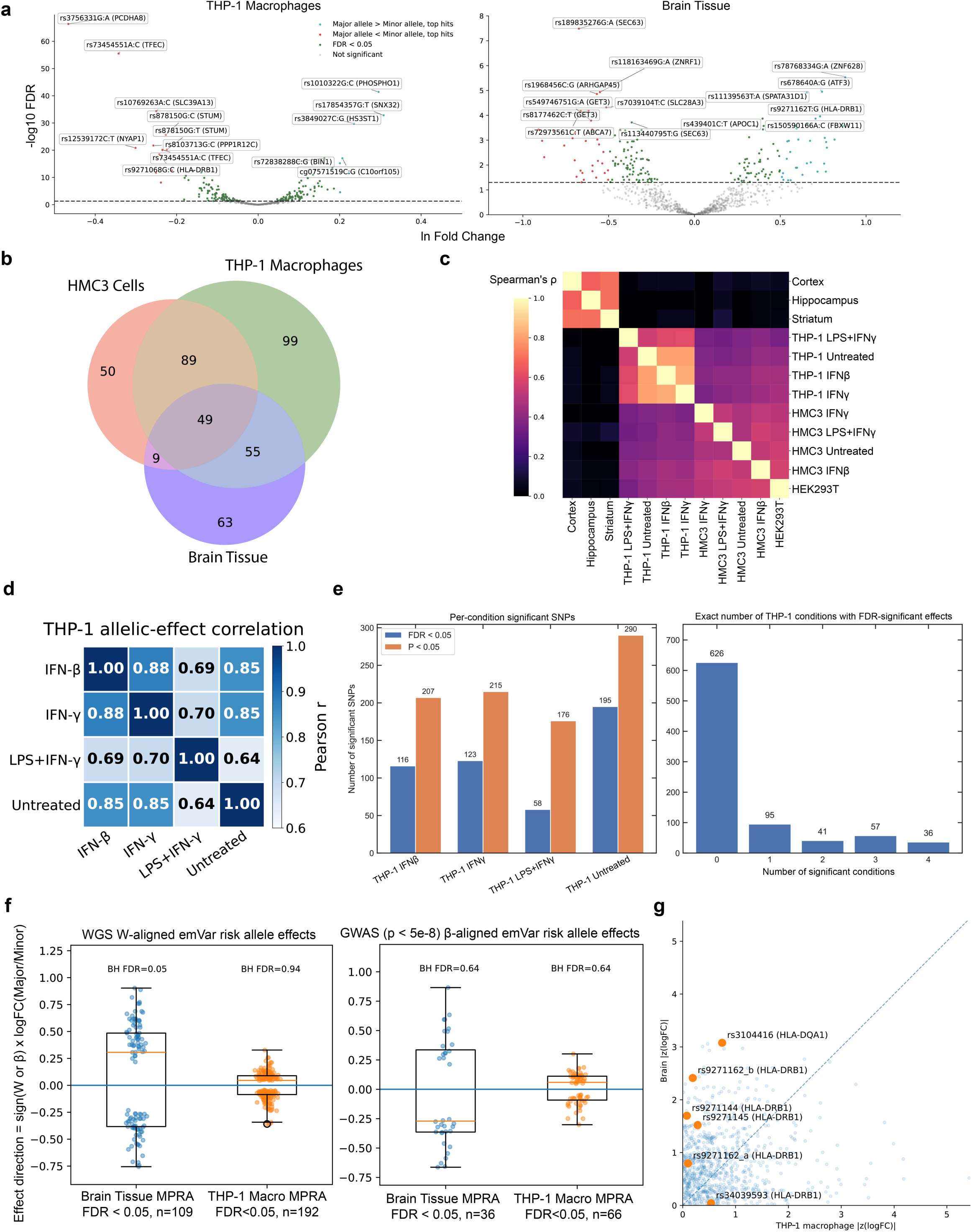
Context-dependent MPRA allelic effects of LOAD-associated variants across immune-cell and brain-tissue contexts. **(a)** Volcano plots showing massively parallel reporter assay (MPRA) allelic effects of late-onset Alzheimer’s disease (LOAD)-associated variants in THP-1 macrophages and mouse brain tissue. The x-axis shows the natural-log fold change between major- and minor-allele constructs, and the y-axis shows −log₁₀(false discovery rate [FDR]) from the MPRAnalyze comparative analysis. Green points indicate variants with significant allelic effects at FDR < 0.05, and gray points indicate variants without significant allelic effects. Selected top hits are highlighted according to allelic direction: cyan indicates greater transcriptional activity of the major allele than of the minor allele, and red indicates greater activity of the minor allele than of the major allele. Labels indicate variant identifiers and nearby or annotated genes. **(b)** Overlap of expression-modulating variants (emVars) identified in HMC3 cells, THP-1 macrophages, and mouse brain tissue. Each set represents variants with significant MPRA allelic effects in the indicated assay context. **(c)** Pairwise comparison of MPRA allelic effect sizes across assay contexts. The heatmap shows Spearman correlation coefficients (ρ) across mouse brain regions, THP-1 macrophage states, HMC3 cell states, and HEK293T cells. Correlations are displayed for context pairs with nominal *P* < 0.05; nonsignificant comparisons are shown in black. Higher values indicate greater concordance of allelic effects between contexts. **(d)** Cross-condition comparison of MPRA allelic effects among THP-1 macrophage states. The heatmap shows Pearson correlations of allelic effect estimates among untreated, IFN-β-stimulated, IFN-γ-stimulated, and LPS + IFN-γ-stimulated THP-1 macrophages. Values within cells indicate Pearson correlation coefficients (*r*). **(e)** Distribution of significant allelic effects across THP-1 macrophage conditions. Left, numbers of variants with significant allelic effects in each condition, defined using either FDR < 0.05 or nominal *P* < 0.05 in the MPRAnalyze comparative analysis. Right, numbers of variants with FDR-significant allelic effects in exactly 0, 1, 2, 3, or 4 THP-1 macrophage conditions. Numbers above the bars indicate variant counts. **(f)** Directionality of MPRA allelic effects after alignment to LOAD genetic risk. Left, WGS risk-allele-aligned emVar effects in mouse brain tissue and THP-1 macrophage MPRA. Right, GWAS β-aligned emVar effects for genome-wide significant variants (*P* < 5 × 10⁻^8^). Effect directions were oriented such that positive values indicate decreased reporter transcription by the WGS risk allele or by the GWAS risk-increasing allele defined by the sign of β, whereas negative values indicate increased transcription. Each point represents one emVar, and the horizontal blue line indicates zero effect. Benjamini–Hochberg-adjusted FDR values from tests of directional bias are shown above each group. **(g)** Comparison of absolute MPRA allelic effect sizes between THP-1 macrophages and mouse brain tissue. The x-axis shows the absolute z-scored allelic log fold change in THP-1 macrophages, and the y-axis shows the corresponding value in mouse brain tissue. Orange points indicate selected variants in the HLA region, labeled by variant identifier and nearby gene, and light-blue points indicate other tested variants. The dashed diagonal line indicates equal absolute allelic effect sizes in the two contexts.

**Extended Data Fig. 5.**
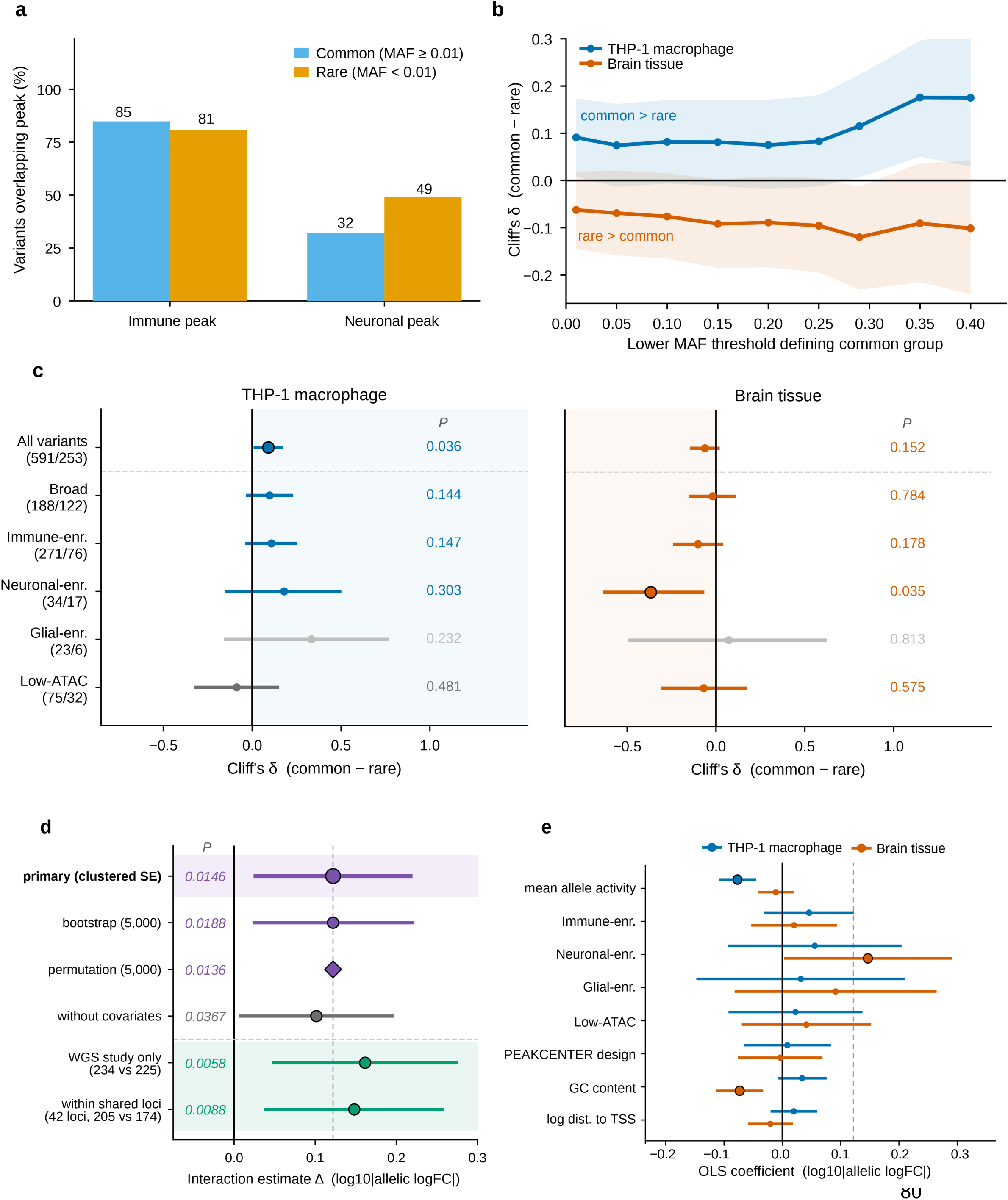
Robustness analyses of common-versus-rare differences in MPRA allelic effect size. Throughout, minor allele frequency (MAF) was taken as the European allele frequency, folded as min(AF, 1 − AF); common variants were defined as MAF ≥ 0.01 (n = 591 constructs) and rare variants as MAF < 0.01 (n = 253 constructs). Allelic effect size was defined as the absolute allelic log fold change estimated using MPRAnalyze. All *P* values are two-sided. **(a)** Overlap of common and rare variants with immune and neuronal open-chromatin peaks. The y-axis indicates the percentage of variants overlapping each peak category. Blue and orange bars represent common and rare variants, respectively, and numbers above the bars indicate percentages. The two sets overlap immune peaks at similar rates and differ mainly in neuronal peak overlap, consistent with the design of the region-based whole-genome sequencing candidate set. Values are descriptive; no inferential test was applied. **(b)** Common-versus-rare differences in allelic effect size across alternative MAF thresholds. The common-variant group was defined using the lower MAF threshold indicated on the x-axis, whereas the rare-variant group was fixed at MAF < 0.01. The y-axis shows Cliff’s δ calculated as common minus rare, with positive values indicating larger effects among common variants and negative values indicating larger effects among rare variants. Blue and orange points represent THP-1 macrophages and brain tissue, respectively, and shaded bands indicate 95% confidence intervals from 6,000 bootstrap resamples. Cliff’s δ remained positive in THP-1 macrophages and negative in brain tissue at every threshold, so the direction of the contrast did not depend on the threshold chosen. Successive thresholds are nested and share the same rare-variant group, and estimates at the highest thresholds rest on progressively smaller common-variant groups (n = 83 at MAF ≥ 0.40); the apparent increase in magnitude at high thresholds is therefore not interpreted as a trend with allele frequency. **(c)** Common-versus-rare differences in allelic effect size across assay for transposase-accessible chromatin using sequencing (ATAC-seq)-defined variant categories. Left, estimates for THP-1 macrophages; right, estimates for brain tissue. The x-axis shows Cliff’s δ calculated as common minus rare. Values in parentheses beside each category indicate the numbers of common and rare constructs, respectively. Points indicate Cliff’s δ estimates, and horizontal bars indicate 95% confidence intervals from 10,000 bootstrap resamples. Exact *P* values from two-sided Mann–Whitney *U*-tests are shown at right. Black-outlined points indicate *P* < 0.05, and gray points indicate categories containing fewer than ten rare variants. Background shading marks the side of zero corresponding to the direction observed in the complete variant set for each context. **(d)** Sensitivity analyses of the allele-frequency-class × assay-context interaction in allelic effect size. Rows are separated by a dashed horizontal line and distinguished by background shading into estimates obtained from the complete variant set (upper rows, purple and gray) and estimates obtained from ascertainment-matched subsets (lower rows, green); the row corresponding to the primary model is additionally highlighted. For the complete variant set, rows show the primary pooled model with construct-clustered standard errors, a construct-level bootstrap with 5,000 resamples, permutation of the rare-variant label with 5,000 permutations, and a model fitted without covariates. For the matched subsets, “WGS study only” indicates the interaction re-estimated using an identical model restricted to variants nominated by the region-based whole-genome sequencing analysis, which contributed both common (n = 234) and rare (n = 225) constructs; “within shared loci” indicates the interaction re-estimated among the 42 loci from that analysis at which both common (n = 205) and rare (n = 174) constructs were assayed, with locus fixed effects included so that common and rare variants are compared within the same locus and with standard errors clustered by locus. Values in parentheses beside each subset label indicate the numbers of common and rare constructs, respectively. The x-axis shows the estimated interaction coefficient, calculated as the difference between the rare-variant coefficients in brain tissue and in THP-1 macrophages. Circles indicate model or bootstrap estimates, horizontal bars indicate 95% confidence intervals, and the diamond indicates the permutation-based estimate, for which a *P* value rather than a confidence interval is obtained. The dashed vertical line marks the estimate from the primary model. Exact *P* values are shown at left. Statistical inference for the interaction rests on the single primary test reported in Fig. 3c; the analyses shown here are supporting and sensitivity analyses, and *P* values are unadjusted for multiple comparisons. **(e)** Covariate coefficients from context-specific ordinary least-squares models of allelic effect size (n = 844 constructs per context). Blue and orange points represent estimates for THP-1 macrophages and brain tissue, respectively. The x-axis shows the regression coefficient, and horizontal bars indicate 95% confidence intervals. Black-outlined points indicate *P* < 0.05. Covariates include mean allele reporter activity, ATAC-seq-defined variant category, construct design, GC content and log-transformed distance to the transcription start site. Continuous covariates were standardized before model fitting, so coefficients are per standard deviation, and ATAC-seq categories were modelled relative to the Broad category. The dashed purple line marks the interaction estimate from **d**, plotted on the same coefficient scale.

**Extended Data Fig. 6.**
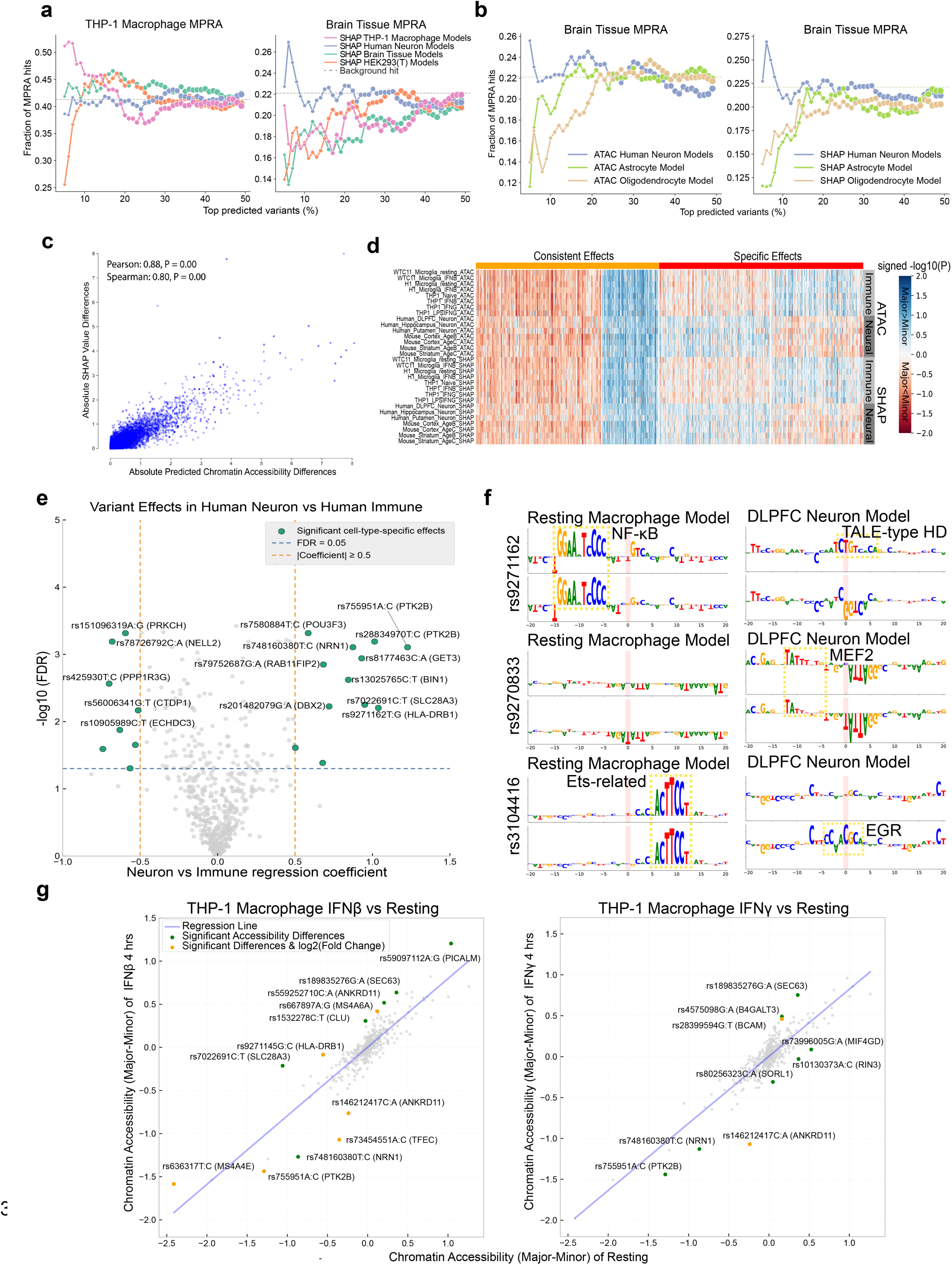
Machine-learning prioritization and interpretation of predicted cell-type- and immune-state-dependent variant effects. **(a)** Enrichment of massively parallel reporter assay (MPRA) hits among candidate variants ranked by SHapley Additive exPlanations (SHAP) scores from different model groups. Left, recovery of THP-1 macrophage MPRA hits; right, recovery of mouse brain-tissue MPRA hits. The x-axis indicates the percentage of top-ranked candidate variants, and the y-axis indicates the fraction identified as MPRA hits. Colored lines represent rankings derived from SHAP scores from THP-1 macrophage, human-neuron, brain-tissue, or HEK293T models. Gray dashed lines indicate the background MPRA hit rate. **(b)** Recovery of mouse brain-tissue MPRA hits using cell-type-specific chromatin-accessibility models. Left, variants ranked by predicted allelic accessibility effects from human-neuron, astrocyte, and oligodendrocyte models. Right, variants ranked by the corresponding SHAP-derived allelic effects. The x-axis indicates the percentage of top-ranked candidate variants, and the y-axis indicates the fraction identified as brain-tissue MPRA hits. **(c)** Relationship between model-predicted allelic chromatin-accessibility differences and SHAP-derived allelic attribution differences. Each point represents a candidate variant. The x-axis shows the absolute predicted accessibility difference between alleles, and the y-axis shows the absolute difference in SHAP attribution values. Pearson and Spearman correlation coefficients are shown; the corresponding *P* values were below numerical reporting precision. **(d)** Heatmap of signed variant-effect significance across assay for transposase-accessible chromatin using sequencing (ATAC-seq)-based predictions and SHAP attribution analyses. Columns indicate variants grouped into consistent-effect and context-specific-effect classes. Rows indicate immune- and neuronal-model comparisons based on predicted accessibility or SHAP attribution scores. Color indicates signed −log₁₀(*P*), with positive values indicating greater predicted accessibility or attribution for the major allele and negative values indicating greater values for the minor allele. **(e)** Identification of variants with differential predicted effects between human-neuron and human-immune models. The x-axis shows the neuron-versus-immune regression coefficient, and the y-axis shows −log₁₀(false discovery rate [FDR]). Green points indicate variants with significant cell-type-specific effects, whereas gray points indicate nonsignificant variants. The horizontal dashed line indicates FDR = 0.05, and the vertical dashed lines indicate absolute regression coefficients of 0.5. Labels identify selected variants and nearby or annotated genes. **(f)** Allele-specific, base-resolution SHAP importance-score logos for rs9271162, rs9270833, and rs3104416 in resting macrophage and dorsolateral prefrontal cortex (DLPFC) neuron models. Paired tracks show importance profiles for the two alleles. Pink shading marks the variant position, and dashed yellow boxes highlight selected local transcription factor motif contexts. Motif labels indicate representative matched motifs. **(g)** Comparison of predicted allelic chromatin-accessibility effects between resting and stimulated THP-1 macrophage models. Left, IFN-β stimulation for 4 h versus resting; right, IFN-γ stimulation for 4 h versus resting. The x-axis shows the predicted major-minus-minor accessibility difference in the resting model, and the y-axis shows the corresponding difference in the stimulated model. Blue lines indicate linear regression fits. Green points indicate variants with significant predicted allelic accessibility differences, orange points indicate variants that additionally show significant stimulation-dependent log₂ fold-change differences, and gray points indicate other tested variants. Labels identify selected variants and nearby or annotated genes.

**Extended Data Fig. 7.**
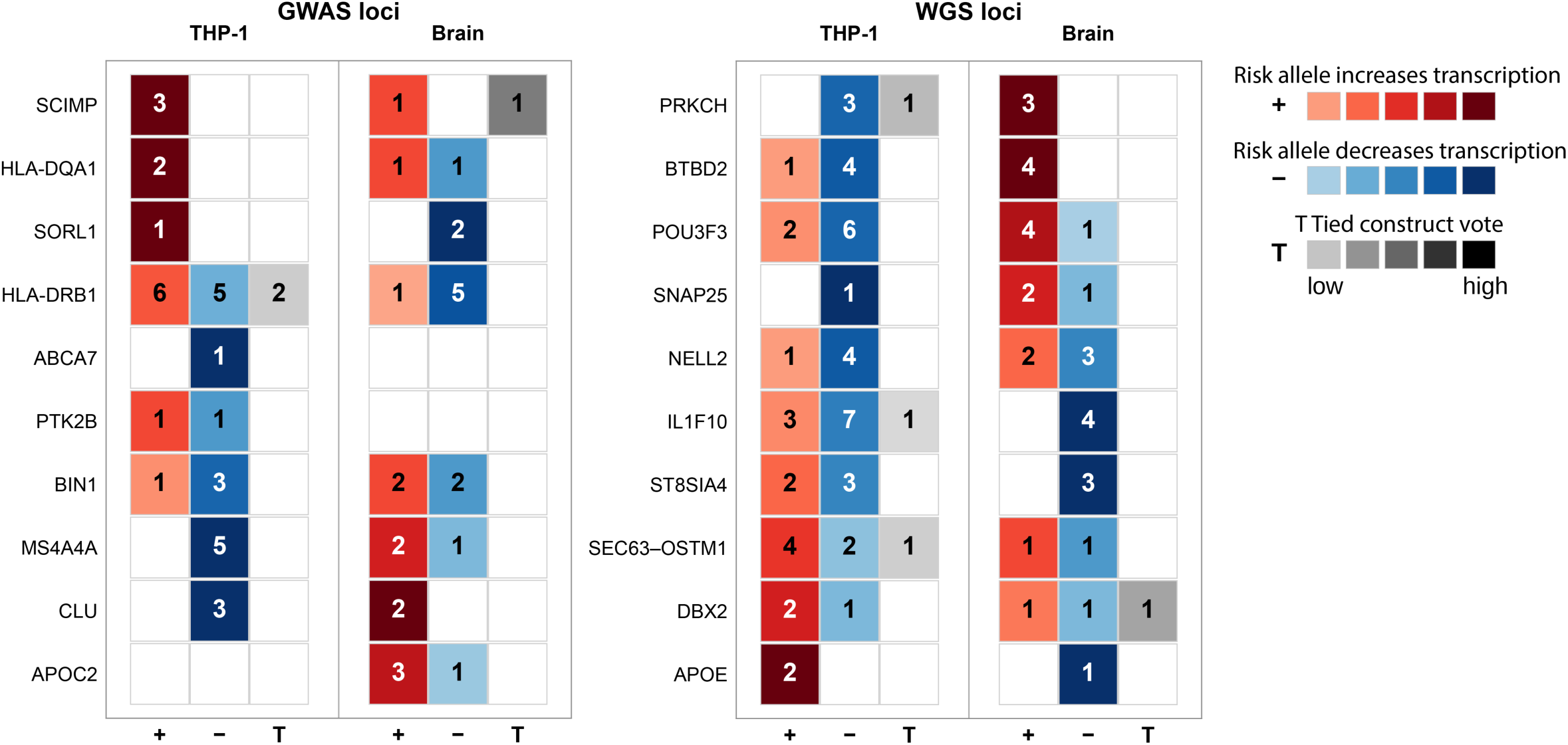
Association-aligned directionality of MPRA allelic effects across LOAD-associated GWAS and WGS loci. Locus-level summary of expression-modulating variants at LOAD-associated genome-wide association study (GWAS) and whole-genome sequencing (WGS) loci in THP-1 macrophages and mouse brain tissue. Rows indicate loci, and columns classify variants according to whether the risk allele increased transcription (+), decreased transcription (−), or produced a tied construct-level vote (T). Numbers within cells indicate the number of MPRA-significant variants in each category. Red, blue, and gray shading indicate transcription-increasing, transcription-decreasing, and tied effects, respectively, with darker shading indicating a stronger construct-level vote.

**Extended Data Fig. 8.**
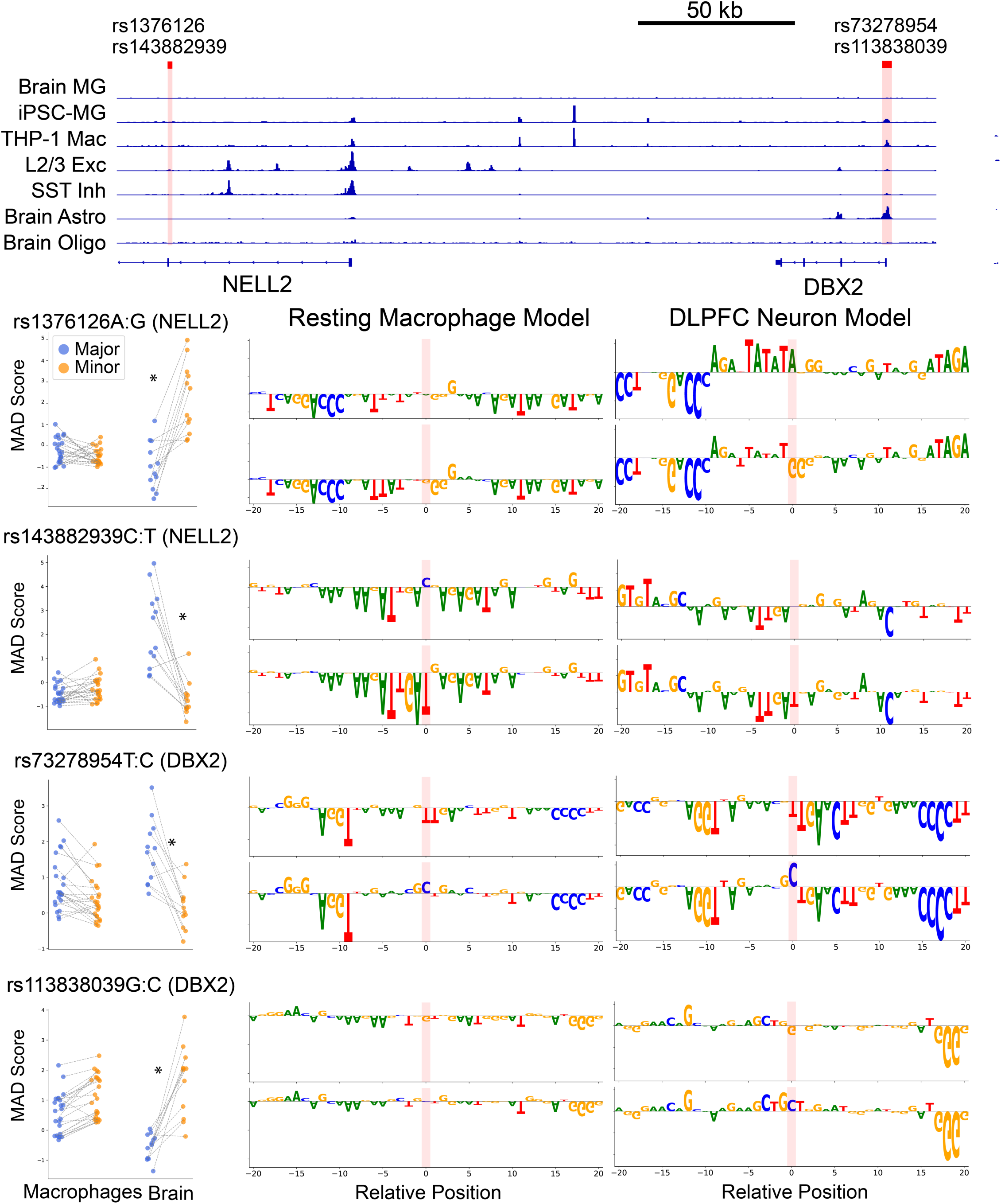
Context-dependent regulatory effects of rare variants across the *NELL2–DBX2* region. Top, chromatin-accessibility profiles across the *NELL2–DBX2* locus in brain microglia, induced pluripotent stem cell-derived microglia (iPSC-MG), THP-1 macrophages, layer 2/3 excitatory neurons, somatostatin-positive inhibitory neurons, brain astrocytes, and brain oligodendrocytes. Pink shading marks the positions of four rare variants selected for massively parallel reporter assay (MPRA) analysis: rs1376126 and rs143882939 near *NELL2*, and rs73278954 and rs113838039 near *DBX2*. Gene annotations are shown below the accessibility profiles, and the scale bar represents 50 kb. Bottom left, MPRA activity of the major- and minor-allele constructs for each variant in macrophage and mouse brain contexts. Activity is represented by median absolute deviation (MAD) scores. Blue and orange points indicate major- and minor-allele constructs, respectively, and gray lines connect paired measurements. Asterisks indicate significant allelic differences at false discovery rate (FDR) < 0.05. Bottom middle and right, allele-specific, base-resolution importance-score logos generated using the resting macrophage and dorsolateral prefrontal cortex (DLPFC) neuron sequence-to-function models, respectively. Paired tracks show the local importance profiles of the major- and minor-allele sequences. Pink shading marks the variant position at relative position 0.

**Extended Data Fig. 9.**
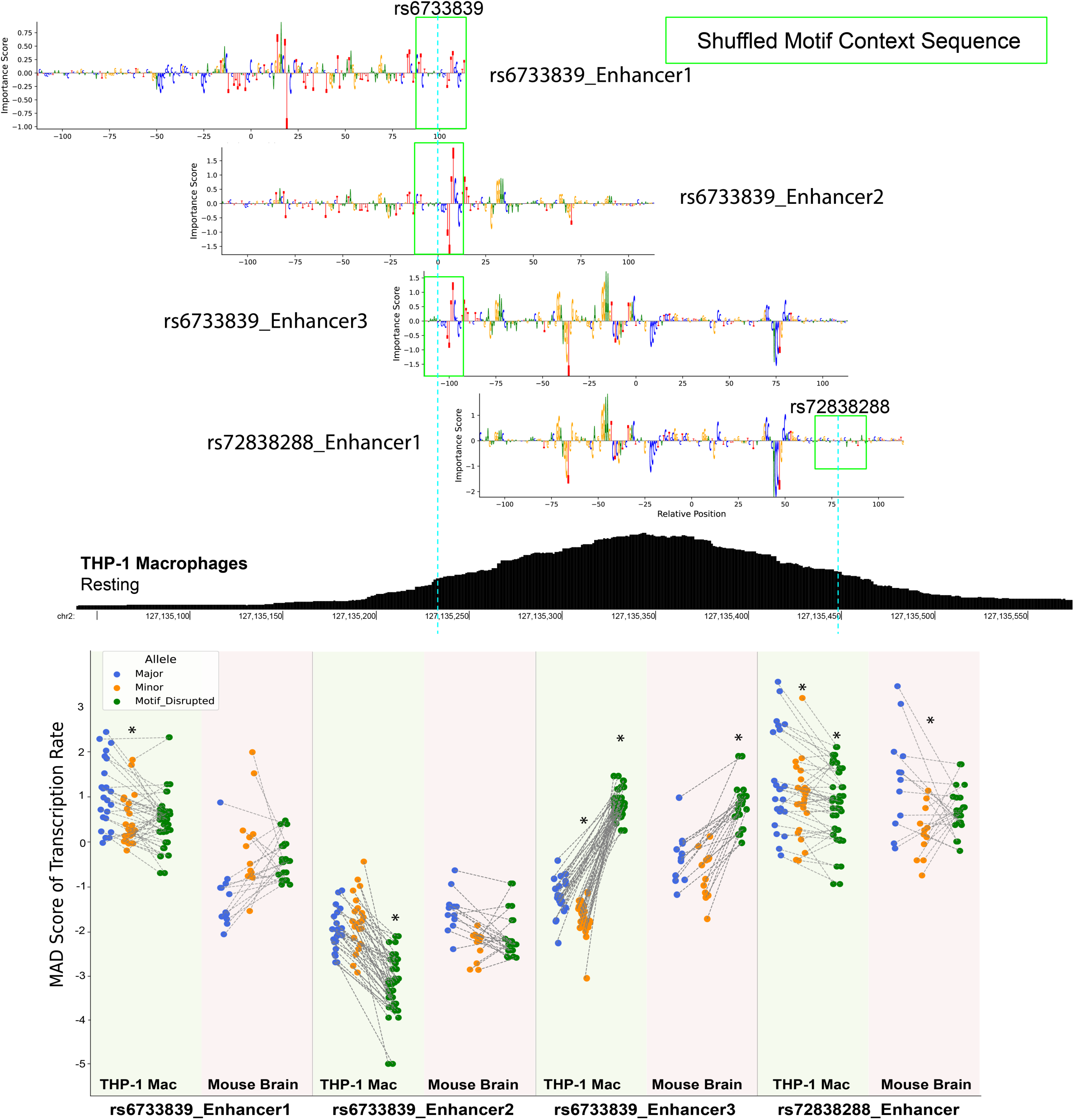
Tiling-based dissection and motif-context shuffling of a *BIN1* enhancer. Top, model-derived base-resolution importance-score tracks for tiled constructs spanning a *BIN1* enhancer containing rs6733839 and rs72838288. Three overlapping constructs are shown for rs6733839, together with one construct containing rs72838288. Green boxes indicate the sequence windows selected for motif-context shuffling, and cyan dashed vertical lines mark the genomic positions of rs6733839 and rs72838288. Middle, chromatin-accessibility signal from resting THP-1 macrophages across the corresponding *BIN1* enhancer. Bottom, massively parallel reporter assay (MPRA) transcriptional activity of major-allele, minor-allele, and motif-shuffled constructs in THP-1 macrophages and mouse brain tissue. Blue, orange, and green points indicate major-allele, minor-allele, and motif-shuffled constructs, respectively. Each point represents an independent biological replicate samples, and gray dashed lines connect matched measurements from the same biological replicate across construct types. Activity is represented by median absolute deviation (MAD) scores. Asterisks indicate significant pairwise differences at false discovery rate (FDR) < 0.05.

**Extended Data Fig. 10.**
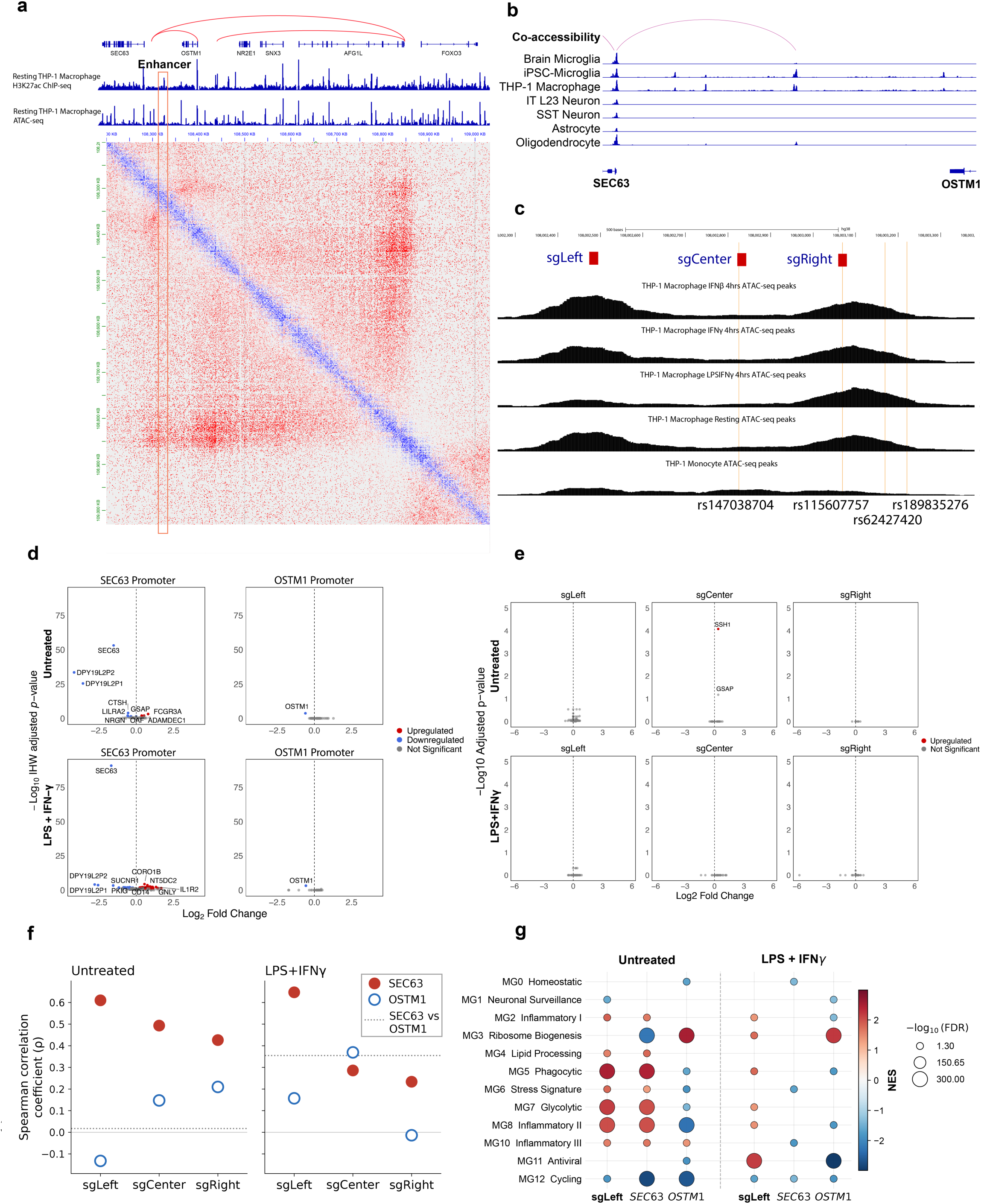
Chromatin architecture and transcriptional consequences of CRISPRi perturbation at the *SEC63–OSTM1* locus. **(a)** Local chromatin architecture surrounding the *SEC63–OSTM1* locus in resting THP-1 macrophages. H3K27ac ChIP–seq and ATAC-seq profiles are shown above the Hi-C contact map. The prioritized enhancer is outlined in orange, and red arcs indicate chromatin contacts between the enhancer-containing region and nearby genomic regions. Gene annotations include *SEC63*, *OSTM1*, *NR2E1*, *SNX3*, *AFG1L*, and *FOXO3*. **(b)** Chromatin co-accessibility and accessibility profiles across the *SEC63–OSTM1* locus in brain microglia, induced pluripotent stem cell-derived microglia (iPSC-MG), THP-1 macrophages, layer 2/3 excitatory neurons, somatostatin-positive inhibitory neurons, astrocytes, and oligodendrocytes. The pink arc indicates a co-accessible connection spanning the locus. **(c)** Design of CRISPR interference (CRISPRi) perturbations targeting the prioritized enhancer. The positions of sgLeft, sgCenter, and sgRight are shown relative to THP-1 ATAC-seq profiles obtained after 4 h of IFN-β, IFN-γ, or LPS + IFN-γ stimulation and under resting macrophage and monocyte conditions. The positions of rs147038704, rs115607757, rs62427420, and rs189835276 are indicated below the tracks. **(d)** Gene-expression changes following CRISPRi targeting of the *SEC63* or *OSTM1* promoter in untreated and LPS + IFN-γ-treated THP-1 macrophages. Columns indicate the targeted promoter, and rows indicate treatment condition. The x-axis shows log₂ fold change, and the y-axis shows −log₁₀(IHW-adjusted P value). Red and blue points indicate significantly upregulated and downregulated genes, respectively, and gray points indicate nonsignificant genes. Labels identify selected genes. **(e)** Gene-expression changes following CRISPRi targeting of the left, center, or right region of the prioritized enhancer in untreated and LPS + IFN-γ-treated THP-1 macrophages. Columns indicate the targeted enhancer region, and rows indicate treatment condition. Axes and point colors are defined as in **d**. **(f)** Concordance between transcriptomic responses to enhancer and promoter perturbation. Points show Spearman correlation coefficients (ρ) between genome-wide gene-expression changes following sgLeft, sgCenter, or sgRight perturbation and those following *SEC63* or *OSTM1* promoter perturbation in untreated and LPS + IFN-γ-treated THP-1 macrophages. Filled red circles indicate correlations with *SEC63* promoter perturbation, and open blue circles indicate correlations with *OSTM1* promoter perturbation. The dotted horizontal line indicates the correlation between the *SEC63* and *OSTM1* promoter-perturbation responses within each treatment condition. **(g)** Gene set enrichment analysis of microglial-state marker gene sets defined by Sun et al. following sgLeft enhancer perturbation and *SEC63* or *OSTM1* promoter perturbation in untreated and LPS + IFN-γ-treated THP-1 macrophages. Rows indicate the microglial states defined by Sun et al., and columns indicate perturbation targets within each treatment condition. Dot color indicates the normalized enrichment score (NES), and dot size indicates −log₁₀(false discovery rate [FDR]).

