## Supplementary Figure for "Context-dependent regulatory variants in Alzheimer’s disease"

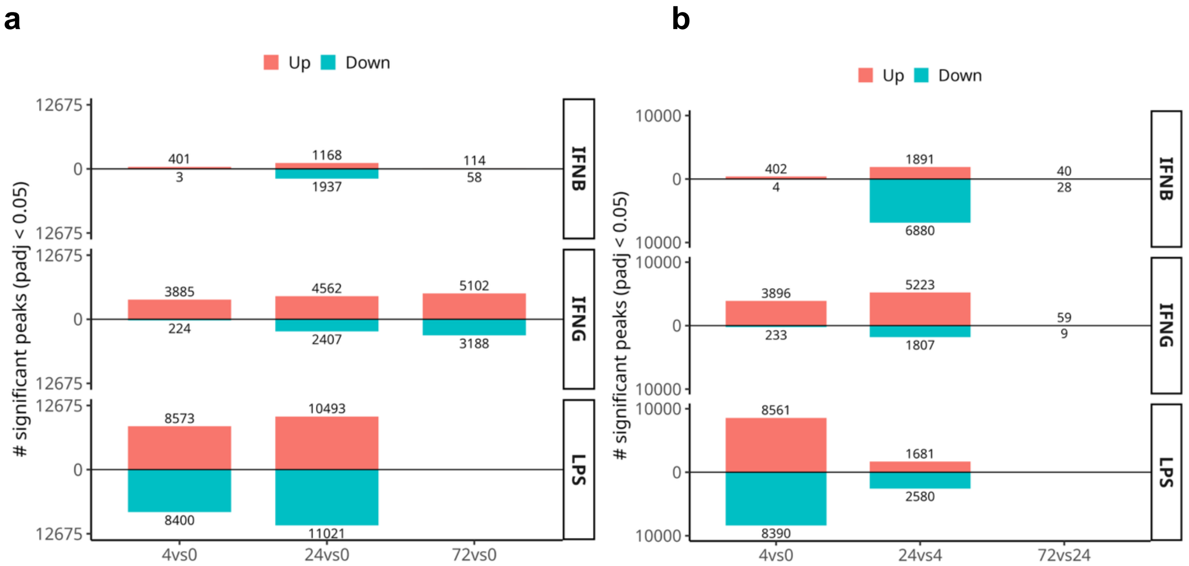


**Supplementary Fig. 1 | Numbers of differential chromatin-accessibility peaks across cytokine-stimulation time-course contrasts.** Bar plots show the numbers of significantly differential ATAC-seq peaks across IFN-β, IFN-γ and LPS+IFN-γ stimulation time courses. Red bars indicate peaks with increased accessibility, and teal bars indicate peaks with decreased accessibility. Numbers adjacent to the bars indicate the corresponding numbers of differential peaks. **a,** Comparisons between each stimulated time point and the unstimulated baseline, including 4 h versus 0 h, 24 h versus 0 h and 72 h versus 0 h. **b,** Sequential time-course comparisons, including 4 h versus 0 h, 24 h versus 4 h and 72 h versus 24 h. Rows indicate stimulation conditions. Significance was defined as adjusted $P < 0.05$.

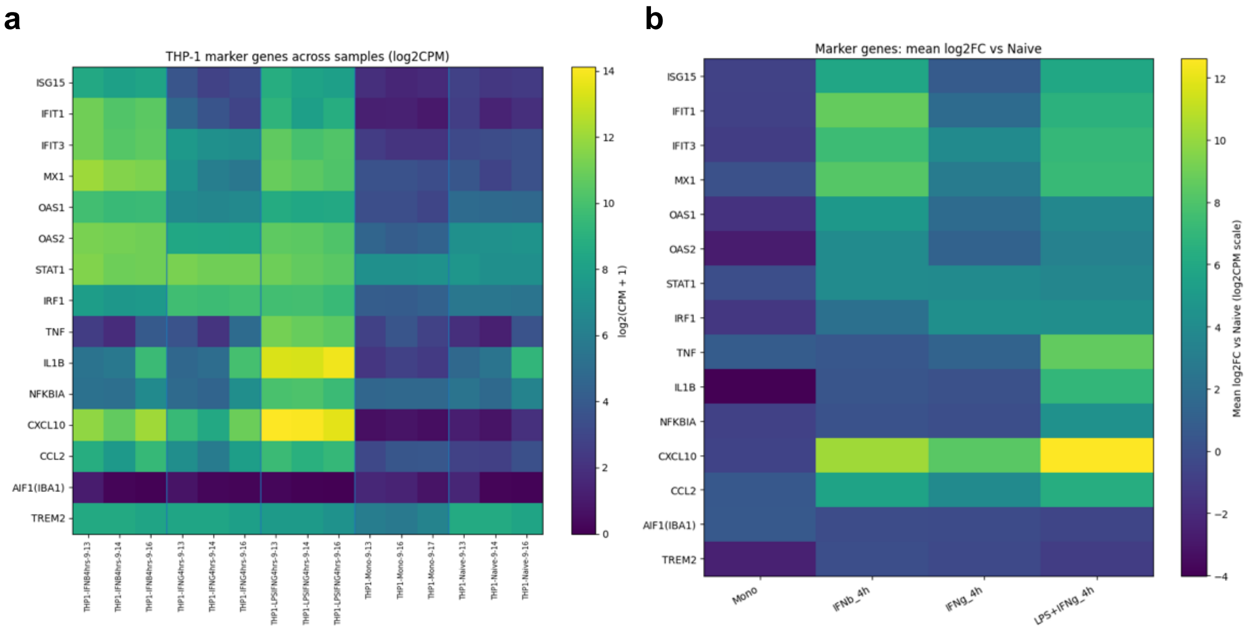


**Supplementary Fig. 2 | Marker-gene expression across untreated and stimulated THP-1 macrophage conditions. a,** Heatmap showing log2(CPM + 1) expression of canonical interferon-stimulated genes and inflammatory response markers across THP-1 samples (three biological replicates per condition). Samples are grouped by condition (IFN-β 4 h, IFN-γ 4 h, LPS+IFN-γ 4 h, monocytes, and naïve) and ordered by replicate date. Expression was computed from the gene-level count matrix by counts-per-million (CPM) normalization followed by log2(CPM + 1) transformation. **b,** Mean log2 fold-change (log2FC) of the same marker genes relative to the naïve (untreated) condition. For each gene, log2FC was calculated on the log2(CPM + 1) scale as the difference between the condition mean and the naïve (untreated) mean across biological replicates; plotted values represent the mean log2FC for each condition. Marker genes include interferon-response genes (e.g., *ISG15, IFIT1/3, MX1, OAS1/2, STAT1, IRF1*) and inflammatory/chemokine genes (e.g., *TNF, IL1B, NFKBIA, CXCL10, CCL2*), as well as microglia/macrophage markers (*AIF1/IBA1 and TREM2*).

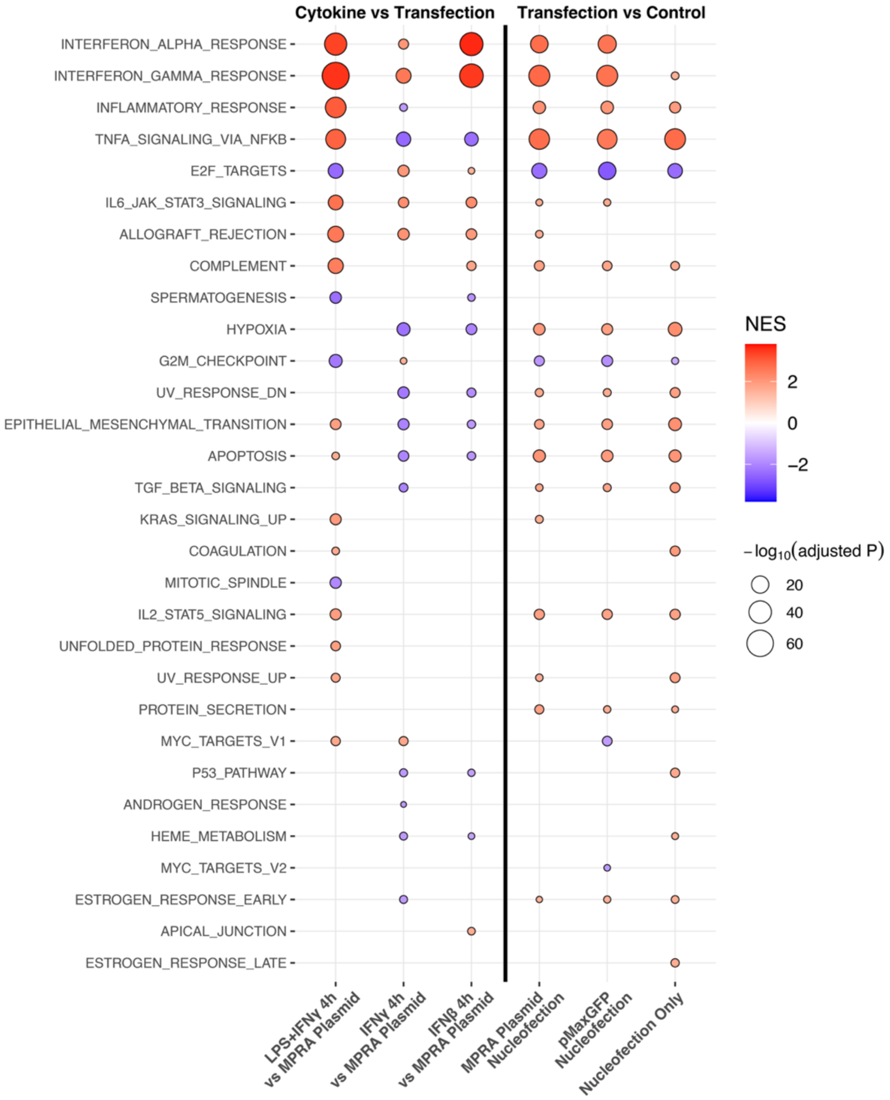


**Supplementary Fig. 3 | Hallmark pathway enrichment across cytokine-stimulation and transfection-related contrasts.** Gene set enrichment analysis of MSigDB Hallmark pathways across the indicated comparisons. The left panel shows cytokine-stimulated conditions relative to MPRA plasmid-transfected cells, including LPS+IFN-γ, IFN-γ and IFN-β stimulation. The right panel shows transfection- or nucleofection-related conditions relative to the corresponding control, including MPRA plasmid nucleofection, pMaxGFP nucleofection and nucleofection-only conditions. Rows indicate Hallmark pathways, and columns indicate pairwise contrasts. Dot color represents the normalized enrichment score (NES), with red indicating positive enrichment and blue indicating negative enrichment. Dot size represents $-\log_{10}$(adjusted $P$). Only selected significantly enriched pathways are shown.

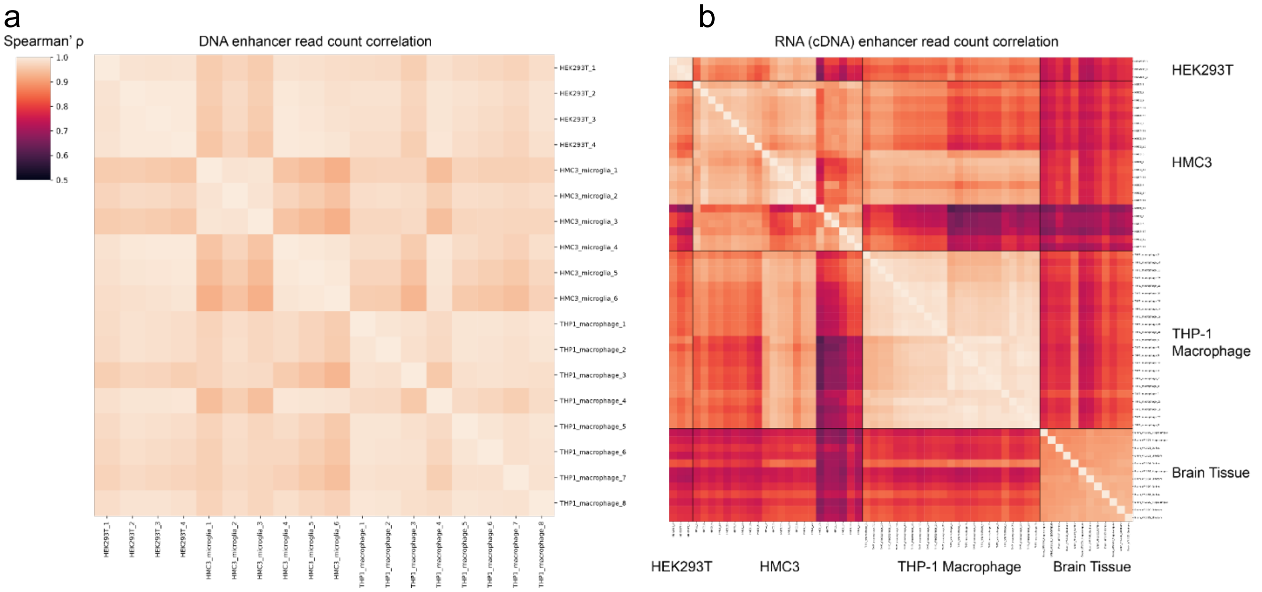


**Supplementary Fig. 4 | Reproducibility of MPRA enhancer-level read counts across DNA and RNA samples.** Heatmaps show pairwise Spearman correlation coefficients for enhancer-level read counts across MPRA samples. **a,** Correlation matrix of DNA plasmid-library read counts across HEK293T, HMC3 and THP-1 macrophage samples. **b,** Correlation matrix of RNA-derived cDNA read counts across HEK293T, HMC3, THP-1 macrophage and brain-tissue MPRA samples. Rows and columns represent individual samples. Black lines in **b** indicate assay-context boundaries. Color represents Spearman's ρ, with lighter colors indicating higher correlation and darker colors indicating lower correlation.

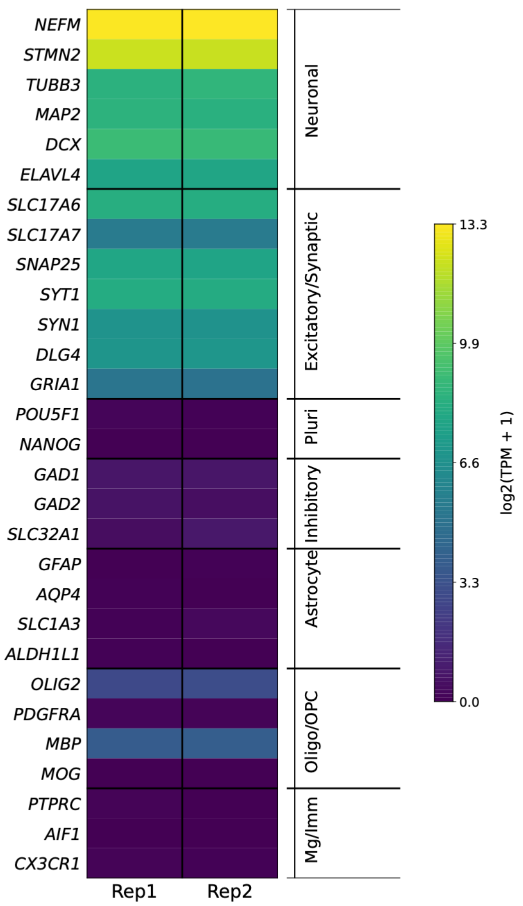


**Supplementary Fig. 5 | Marker-gene expression in iPSC-derived excitatory neurons.** Heatmap showing the expression of selected lineage- and cell-type marker genes in two biological replicates of iPSC-derived excitatory neurons. Rows indicate marker genes grouped by annotation category, including neuronal, excitatory and synaptic, pluripotency, inhibitory neuronal, astrocyte, oligodendrocyte and oligodendrocyte precursor cell (OPC), and microglial and immune markers. Columns indicate biological replicates. Color represents $\log_2$(TPM + 1) expression.

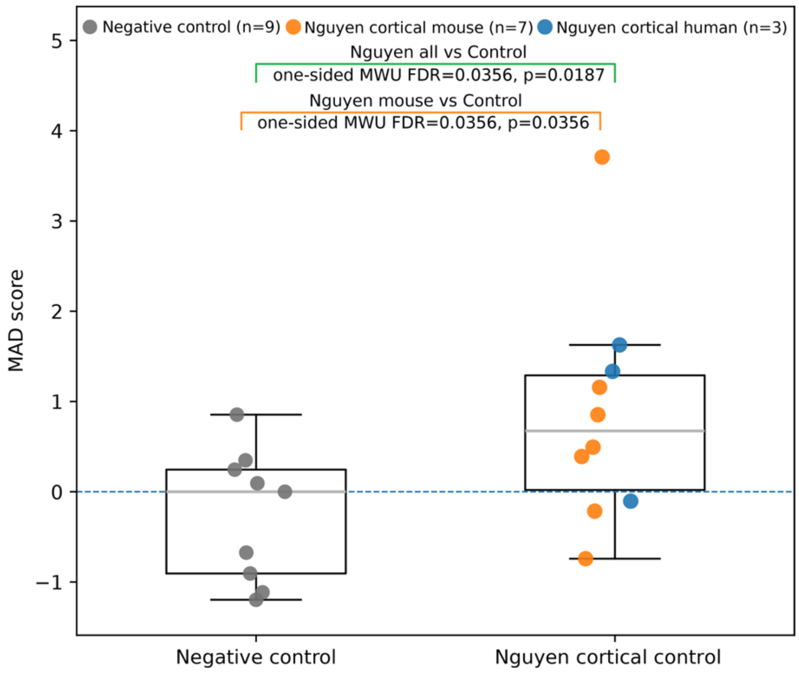


**Supplementary Fig. 6 | MPRA activity of Nguyen cortical positive-control sequences in human iPSC-derived excitatory neurons.** Boxplots show median absolute deviation (MAD) scores for negative-control sequences and Nguyen cortical positive-control sequences. Individual points represent individual sequences. Negative controls are shown in grey ($n = 9$), Nguyen mouse cortical sequences in orange ($n = 7$) and Nguyen human cortical sequences in blue ($n = 3$); the combined Nguyen cortical-control group includes both mouse and human sequences. The dashed horizontal line indicates a MAD score of 0. Statistical comparisons were performed using one-sided Mann–Whitney U-tests, with false-discovery rate (FDR)-adjusted *P* values shown for the combined Nguyen cortical-control group versus negative controls and for Nguyen mouse cortical sequences versus negative controls.

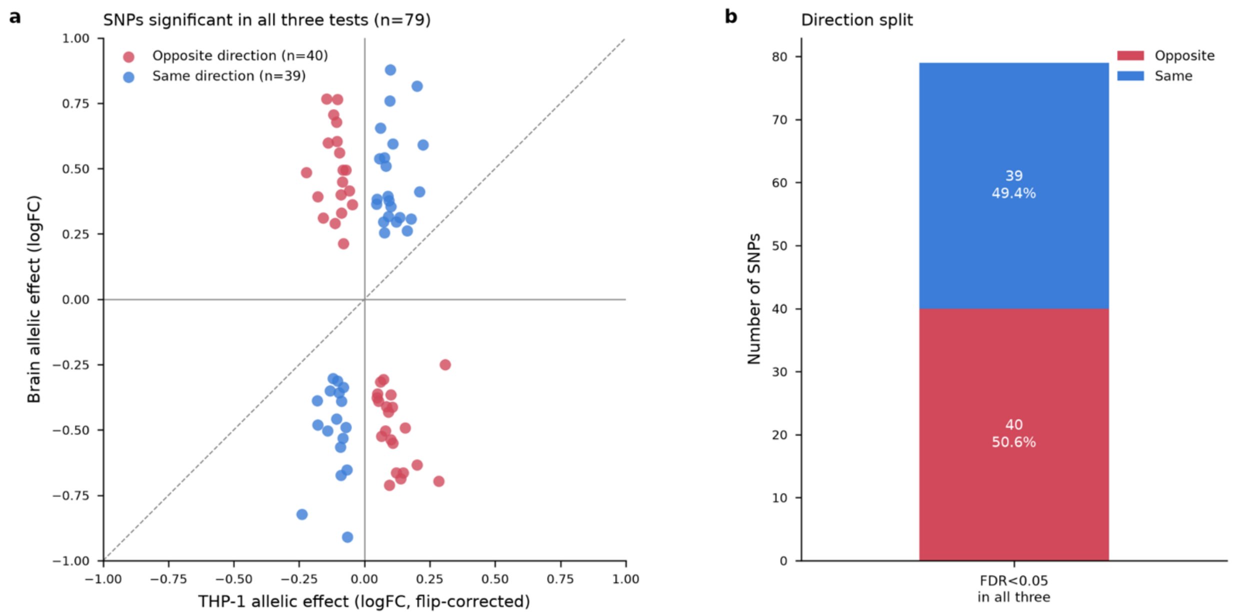


**Supplementary Fig. 7. Allelic effect directions of elements with significant context-dependent MPRA activity. a,** Scatter plot comparing allele-specific MPRA effects between THP-1 macrophages and brain tissue for 79 tested elements with significant allelic effects in both contexts and a significant context-by-allele interaction. Elements were required to pass FDR < 0.05 in THP-1 macrophages, FDR < 0.05 in brain tissue, and FDR < 0.05 for the THP-1 × brain interaction test; these 79 elements correspond to 77 unique SNPs. The x-axis shows flip-corrected allelic logFC in THP-1 macrophages, and the y-axis shows flip-corrected allelic logFC in brain tissue, with alleles oriented consistently across contexts. Red points indicate elements with opposite allelic effect directions between contexts (n = 40), and blue points indicate elements with concordant directions (n = 39). Solid gray lines indicate zero effect in each context, and the dashed line indicates the identity line (y = x). Points in the second and fourth quadrants represent direction reversals. **b,** Proportion of elements with opposite or concordant allelic effect directions among the 79 elements passing the three significance filters. Opposite-direction effects were detected for 40 elements (50.6%), whereas concordant-direction effects were detected for 39 elements (49.4%). Direction was assigned based on the sign of the flip-corrected allelic logFC in each context.

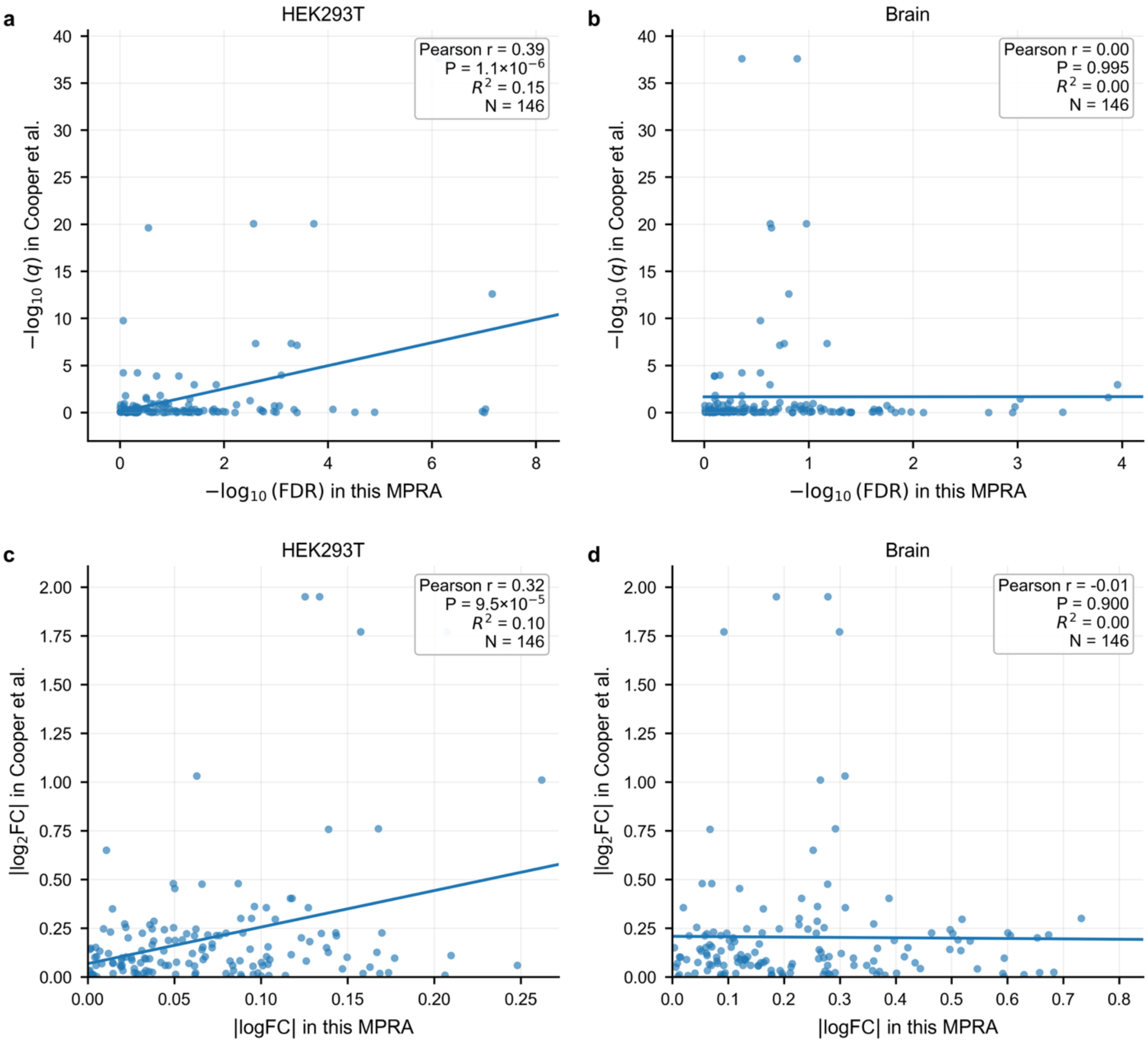


**Supplementary Fig. 8 | Comparison of allelic effects measured in HEK293T cells and brain tissue with Cooper *et al.* MPRA data.** Scatter plots compare allelic-effect measurements for 146 variants shared between this study and the Cooper *et al.* MPRA dataset. **a,b,** Comparison of statistical significance, shown as $-\log_{10}$(FDR) in this study and $-\log_{10}(q)$ in Cooper *et al.*, for the HEK293T (**a**) and brain-tissue (**b**) MPRAs. **c,d,** Comparison of absolute allelic effect sizes, shown as $|\log_2$ fold change$|$ in this study and Cooper *et al.*, for the HEK293T (**c**) and brain-tissue (**d**) MPRAs. Each point represents one overlapping variant, and blue lines indicate linear regression fits. Pearson's *r*, corresponding two-sided *P* values, $R^2$ values and sample sizes are shown in each panel.

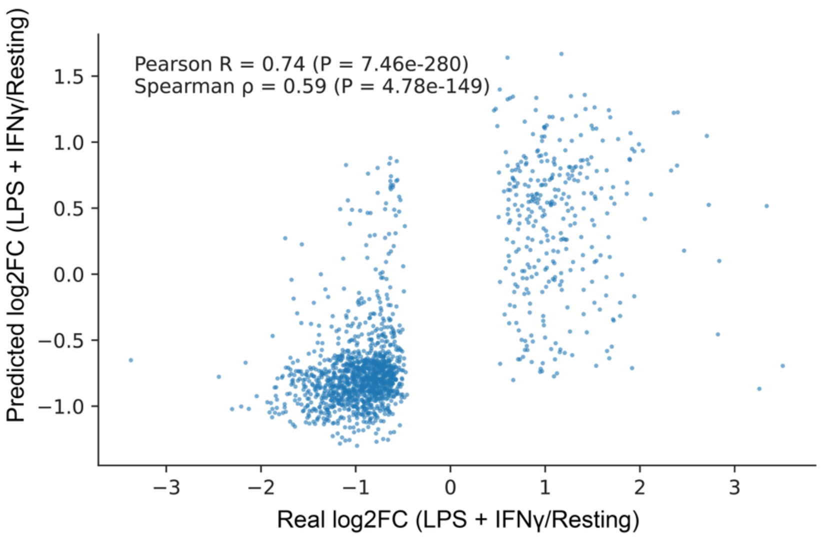


**Supplementary Fig. 9 | Observed and CNN-predicted accessibility changes at LPS+IFN-γ-responsive peaks in THP-1 macrophages.** Scatter plot comparing observed and convolutional neural network (CNN)-predicted $\log_2$ fold changes for ATAC-seq peaks responsive to LPS+IFN-γ stimulation. The x axis shows the observed $\log_2$ fold change between LPS+IFN-γ-stimulated and untreated THP-1 macrophages, and the y axis shows the corresponding CNN-predicted $\log_2$ fold change. Models were trained on LPS+IFN-γ-responsive differentially accessible regions identified from ATAC-seq data using DESeq2 (FDR < 0.05). Each point represents one peak. Pearson's *r*, Spearman's ρ and the corresponding *P* values are shown.

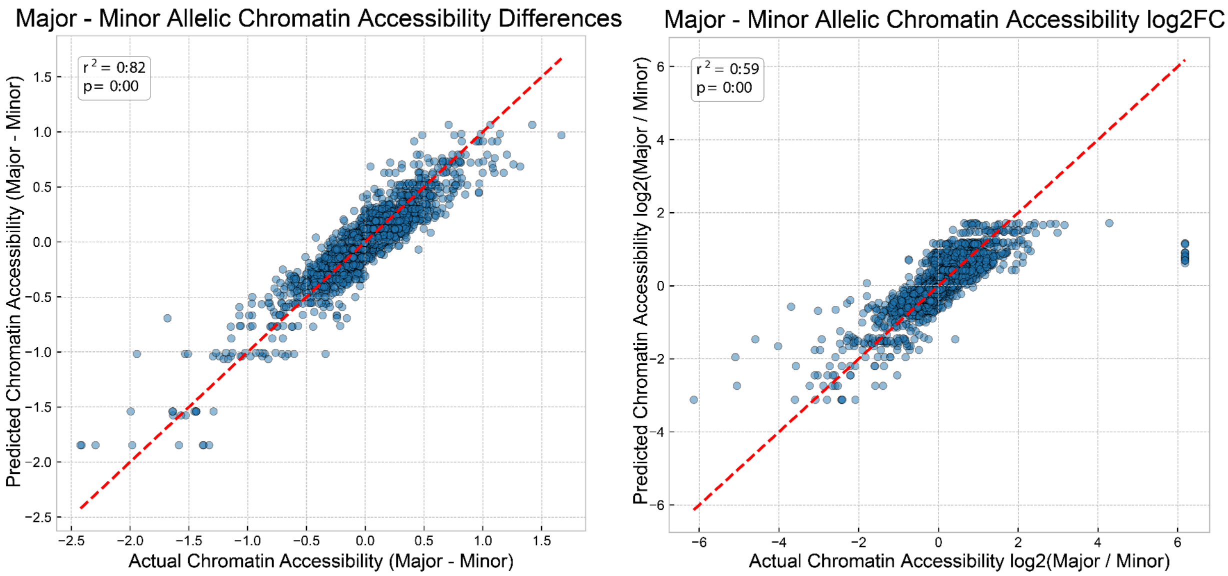


**Supplementary Fig. 10 | Agreement between observed and model-predicted major–minor allelic differences in chromatin accessibility.** Scatter plots compare observed and sequence-model-predicted chromatin-accessibility differences between the major and minor alleles. **a,** Major-minus-minor accessibility differences on the original prediction scale. **b,** Major-minus-minor accessibility differences expressed as $\log_2$ fold changes. Each point represents one allelic comparison. Red dashed lines indicate the identity line ($y = x$). Coefficients of determination ($R^2$) and corresponding *P* values are shown in each panel.

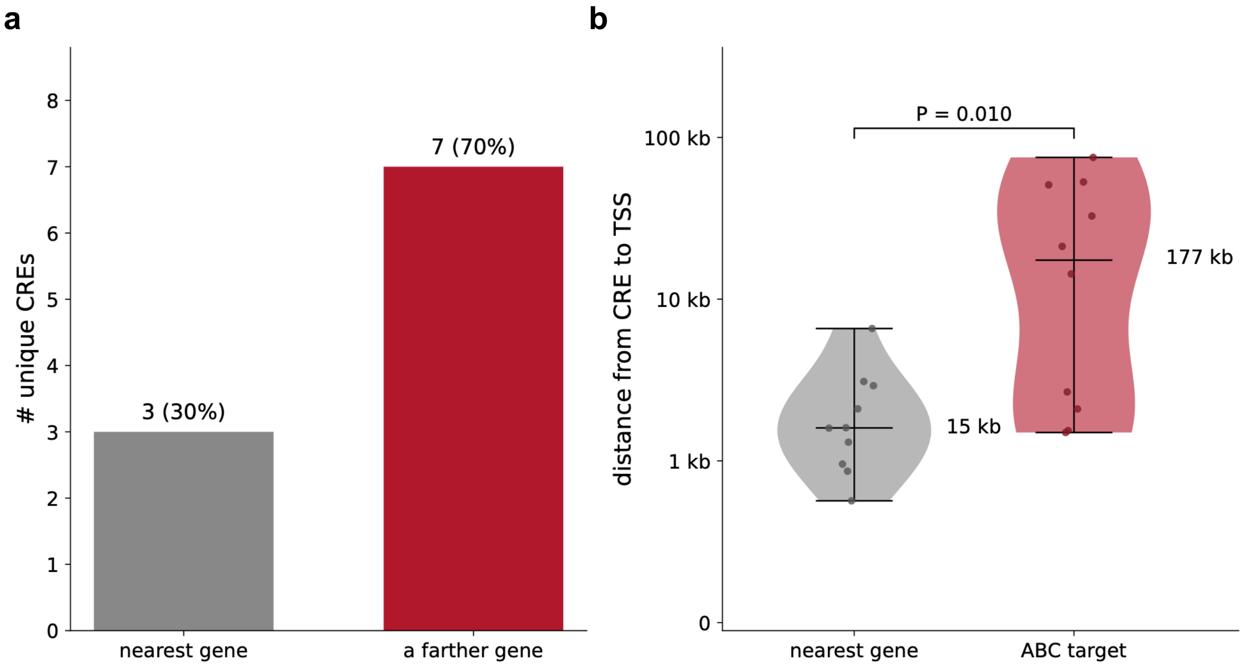


**Supplementary Fig. 11 | ABC-nominated target genes for MPRA-active distal CREs frequently differ from nearest-gene assignments.** Fine-mapped AD-associated variants showing significant allelic effects in the THP-1 macrophage MPRA (FDR < 0.05) and annotated to distal cis-regulatory elements (CREs) were mapped to Activity-by-Contact (ABC) enhancer peaks. ABC peaks overlapping by genomic coordinates across five macrophage or monocyte contexts—untreated, IFN-β-stimulated, IFN-γ-stimulated and LPS+IFN-γ-stimulated THP-1 macrophages, and monocytes—were merged, collapsing 22 variants and 18 ABC peaks into 10 unique CREs. For each CRE, the ABC-nominated target gene was defined as the linked gene with the highest ABC score, whereas the nearest gene was defined as the protein-coding gene with the closest transcription start site (TSS). **a,** Numbers of unique CREs for which the ABC-nominated target was the nearest protein-coding gene (grey) or a more distal gene (red). Seven of ten CREs (70%) were linked to a non-nearest gene. **b,** Distances from each CRE to the nearest-gene TSS (grey) and to the ABC-nominated target-gene TSS (red). Points represent individual CREs ($n = 10$, paired), and horizontal bars indicate medians. ABC-nominated target genes were located a median of 177 kb from the corresponding CREs, compared with 15 kb for the nearest genes, representing an 11.8-fold difference ($P = 0.010$, two-sided Wilcoxon signed-rank test).

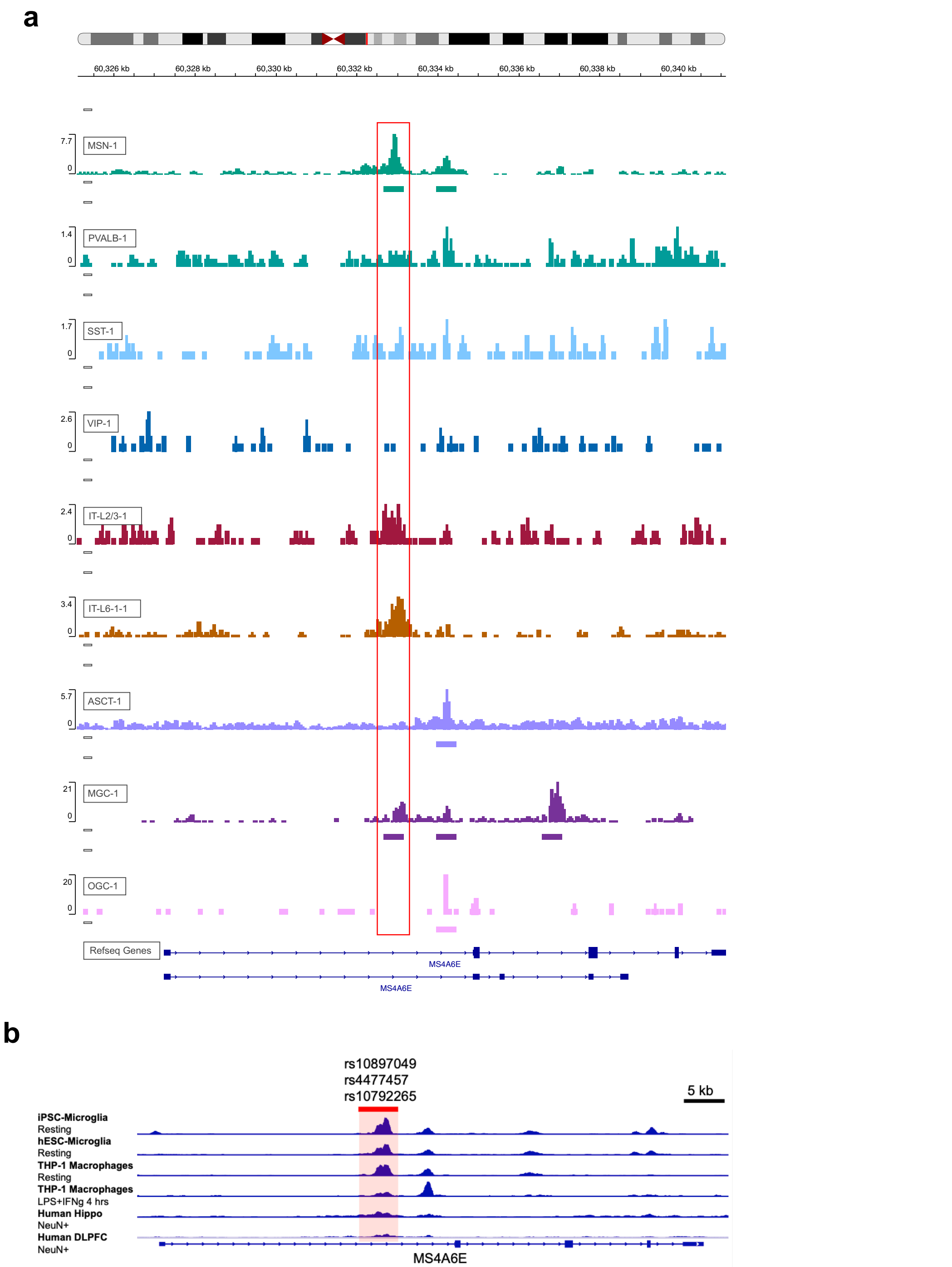


**Supplementary Fig. 12 | Chromatin accessibility at the *MS4A6E* intronic CRE across brain cell subtypes and cellular models.**

**a,** Genome-browser tracks show chromatin accessibility across the *MS4A6E* locus in the indicated neuronal and glial cell subtypes. The red box highlights the intron 1 CRE examined in Fig. 6c. Horizontal bars beneath individual tracks indicate called accessible regions. Track-specific signal ranges are shown on the left, genomic coordinates are shown above, and RefSeq gene annotations are shown below. MSN, medium spiny neuron; PVALB, parvalbumin interneuron; SST, somatostatin interneuron; VIP, vasoactive intestinal peptide interneuron; IT-L2/3 and IT-L6, intratelencephalic neurons from cortical layers 2/3 and 6, respectively; ASCT, astrocyte; MGC, microglia; OGC, oligodendrocyte. **b,** Chromatin-accessibility tracks at the same *MS4A6E* intronic CRE in resting iPSC-derived microglia, resting hESC-derived microglia, resting and LPS+IFN-γ-stimulated THP-1 macrophages, and $NeuN^+$ nuclei from the human hippocampus and dorsolateral prefrontal cortex (DLPFC). The red-shaded interval marks the CRE containing rs10897049, rs4477457, and rs10792265. The genomic scale is indicated at the upper right.

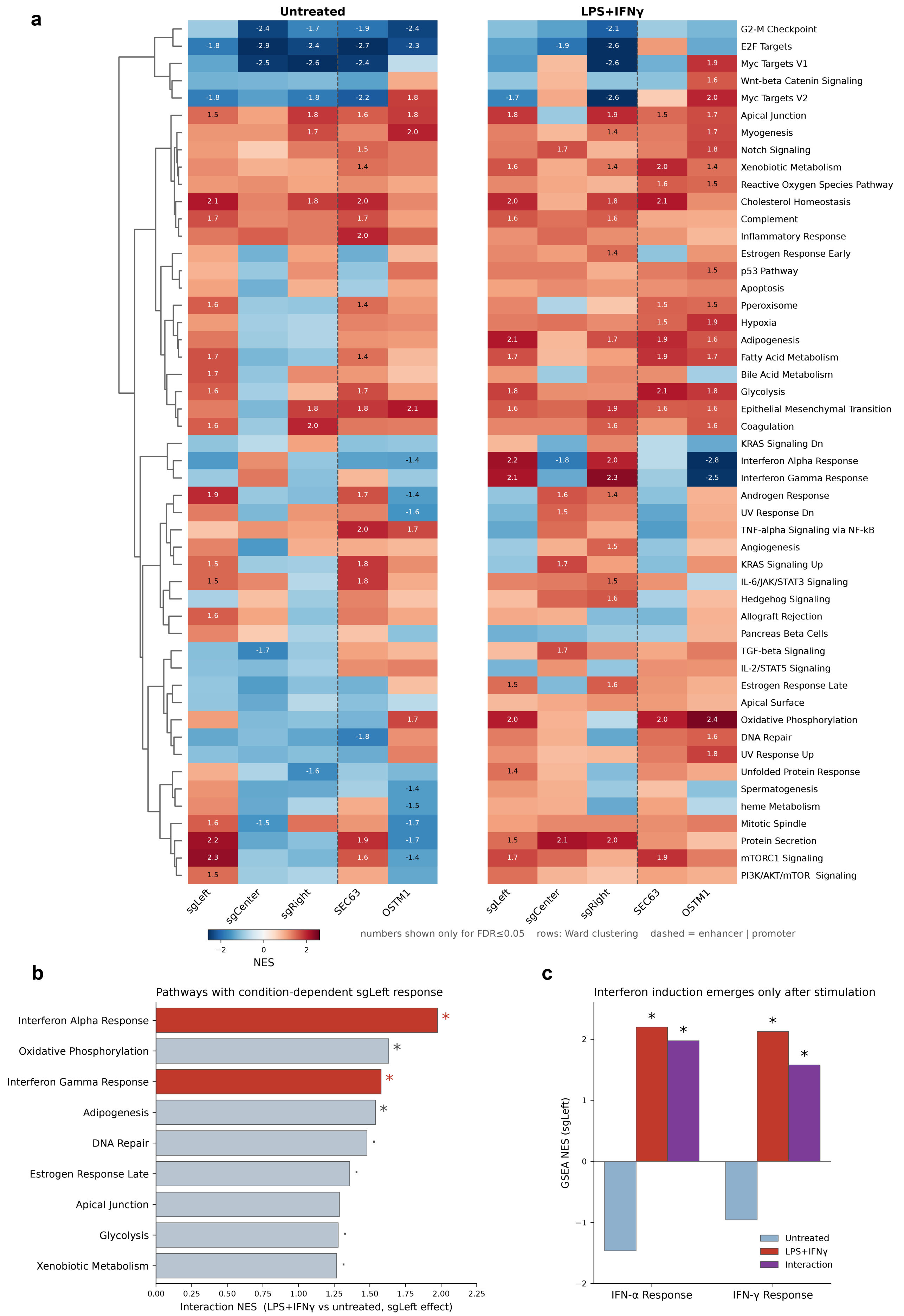
a
Untreated
LPS+IFNγ
G2-M Checkpoint
E2F Targets
Myc Targets V1
Wnt-beta Catenin Signaling
Myc Targets V2
Apical Junction
Myogenesis
Notch Signaling
Xenobiotic Metabolism
Reactive Oxygen Species Pathway
Cholesterol Homeostasis
Complement
Inflammatory Response
Estrogen Response Early
p53 Pathway
Apoptosis
Pperoxisome
Hypoxia
Adipogenesis
Fatty Acid Metabolism
Bile Acid Metabolism
Glycolysis
Epithelial Mesenchymal Transition
Coagulation
KRAS Signaling Dn
Interferon Alpha Response
Interferon Gamma Response
Androgen Response
UV Response Dn
TNF-alpha Signaling via NF-kB
Angiogenesis
KRAS Signaling Up
IL-6/JAK/STAT3 Signaling
Hedgehog Signaling
Allograft Rejection
Pancreas Beta Cells
TGF-beta Signaling
IL-2/STAT5 Signaling
Estrogen Response Late
Apical Surface
Oxidative Phosphorylation
DNA Repair
UV Response Up
Unfolded Protein Response
Spermatogenesis
heme Metabolism
Mitotic Spindle
Protein Secretion
mTORC1 Signaling
PI3K/AKT/mTOR Signaling
sgLeft
sgCenter
sgRight
SEC63
OSTM1
NES
numbers shown only for FDR≤0.05 rows: Ward clustering dashed = enhancer | promoter
b
Pathways with condition-dependent sgLeft response
Interferon Alpha Response
Oxidative Phosphorylation
Interferon Gamma Response
Adipogenesis
DNA Repair
Estrogen Response Late
Apical Junction
Glycolysis
Xenobiotic Metabolism
Interaction NES (LPS+IFNγ vs untreated, sgLeft effect)
c
Interferon induction emerges only after stimulation
GSEA NES (sgLeft)
Untreated
LPS+IFNγ
Interaction
IFN-α Response
IFN-γ Response

**Supplementary Fig. 13 | Pathway-level transcriptional effects of CRISPRi perturbation at the *SEC63–OSTM1* regulatory element.**

**a,** Heatmaps show normalized enrichment scores (NES) from gene set enrichment analysis of MSigDB Hallmark pathways following CRISPR interference (CRISPRi) targeting the *SEC63–OSTM1* regulatory element with sgLeft, sgCenter or sgRight, or targeting the *SEC63* or *OSTM1* promoter, in untreated and LPS+IFN-γ-stimulated THP-1 macrophages. sgLeft, sgCenter and sgRight target the left ATAC-seq subpeak, intervening accessibility valley and right subpeak, respectively. Rows indicate Hallmark pathways ordered by Ward hierarchical clustering; the dashed vertical line separates enhancer- and promoter-targeting perturbations. Color indicates NES relative to the corresponding non-targeting control, with red indicating positive enrichment and blue indicating negative enrichment. Numerical values are shown only for pathways with FDR ≤ 0.05. **b,** Hallmark pathways showing condition-dependent responses to sgLeft perturbation. Bars indicate interaction NES values comparing the sgLeft effect under LPS+IFN-γ stimulation with that under untreated conditions. Asterisks indicate FDR < 0.05 and dots indicate FDR < 0.25. **c,** NES values for the interferon-α and interferon-γ response pathways following sgLeft perturbation under untreated and LPS+IFN-γ-stimulated conditions, together with the treatment-by-perturbation interaction effect. Asterisks indicate FDR < 0.05.
